# Novel roles for pirin proteins and a 2-ketoglutarate: ferredoxin oxidoreductase ortholog in *Bacteroides fragilis* central metabolism and comparison of metabolic mutants susceptibility to metronidazole and amixicile

**DOI:** 10.1101/2024.04.11.589080

**Authors:** Andrea M. Gough, Anita C. Parker, Patricia J. O’Bryan, Terence R. Whitehead, Sourav Roy, Brandon L. Garcia, Paul S. Hoffman, C. Jeffrey Smith, Edson R. Rocha

## Abstract

The regulation of the central metabolism and fermentation pathways and its effect on antimicrobial susceptibility in the anaerobic pathogen *Bacteroides fragilis* is not completely understood. In this study we show that *B. fragilis* encodes for two iron-dependent redox-sensitive regulatory pirin protein genes, *pir1* and *pir2,* whose mRNA expression are upregulated following oxygen exposure and growth in iron-limiting conditions. *Δpir1* and *Δpir2* mutants showed altered short-chain fatty acids production compared to the parent strain. Overexpression of Pir1 and Pir2 increased susceptibility to metronidazole (MTZ), and to amixicile (AMIX), a novel inhibitor of pyruvate:ferredoxin oxidoreductase (PFOR) in anaerobes. We hypothesized these observations were a result of a modulatory effect of Pir1 and Pir2 proteins. Consistent with this, we showed that Pir1 forms direct protein-protein interactions with PFOR. In addition, AlphaFold2-based structural analysis predicts that Pir1 and Pir2 form stable interactions with several enzymes of the central metabolism including the 2-ketoglutate:ferredoxin oxidoreductases (KFOR), Kor1AB and Kor2CDAEBG. A series of metabolic mutants and electron transport chain inhibitors were used to show a wide-ranging effect of bacterial metabolism on MTZ and AMIX susceptibility. There was no cross-resistance between MTZ resistant strains with AMIX susceptibility. Furthermore, we show that AMIX is an effective antimicrobial against *B. fragilis* in an experimental model of intra-abdominal infection. This investigation led to the discovery that the *kor2AEBG* genes are essential for growth, and we present evidence that *kor2AEBG* genes have dual-functions including the reductive synthesis of 2-ketoglutarate via reverse TCA cycle. Support for a novel Kor2AEBG activity comes from the findings showing that addition of compounds containing oleic acid stimulated growth of the *Δkor2AEBG* mutant. However, the metabolic activity that bypasses KorAEBG function remains to be defined. Collectively our investigation reveals new information on *B. fragilis* central metabolism and its modulatory control by pirin proteins. These data provide new genetic and metabolic knowledge which may be leveraged for the future development of new narrow-spectrum antimicrobials.

## INTRODUCTION

Among the anaerobic pathogenic bacteria causing human infections, *Bacteroides fragilis* is the most frequent isolate, and multi-drug-resistant strains are on the rise accounting for most treatment failures (Byun et al., 2019; Hartmeyer et al., 2012; Jasemi et al., 2021; Nagy et al., 2011; Schuetz, 2014, Snydman et al., 2017). Metronidazole (MTZ) remains the antibiotic of choice for the management of infections caused by anaerobes and resistance to MTZ is generally still low, however, resistant strains have been reported in regional survey studies (Alauzet et al., 2019; Hartmeyer et al., 2012; Jasemi et al., 2021, Nagy et al., 2011, Shafquat e al., 2019; Shilnikova & Dmitrieva, 2015). MTZ, is a derivative of the 5-nitroimidazole class of prodrugs requiring (2 pairs of 2 e^−^) or 4 1e^−^ reductions of the nitro group for activation to produce a reactive radical species accountable for its lethal DNA mutagenic and strand fragmentation activity (Alauzet et al., 2019; Dingsdag & Hunter, 2018; Ghotaslou et al., 2018; Sisson et al., 2000). Therefore, in view of the continuous increase in *B. fragilis* multi-drug resistance (MDR) traits and the steady decrease in the number of new alternative antibiotics, there is a need to find alternative anaerobic therapeutics. In this regard, recent studies have reported on a novel antimicrobial, amixicile, that exhibits excellent *in vitro* and *in vivo* activity against oral anaerobic bacteria pathogens grown on biofilm or internalized by host cells (Hutcherson et al., 2017; Gui et al., 2019; Gui et al., 2020; Gui et al., 2021; Reed et al., 2018).

Amixicile (AMIX), a novel second generation of nitazoxanide (Hoffman, 2020), selectively inhibits the action of thiamine-diphosphate (ThPP) cofactor in pyruvate:ferredoxin oxidoreductase (PFOR) and related members of the 2-ketoacid ferredoxin oxidoreductase (KFOR) superfamily found in obligate anaerobic bacteria, anaerobic human intestinal parasites, and in members of the epsilonproteobacteria such as *Campylobacter* and *Helicobacter* (Hoffman et al., 2014; Hoffman, 2020; Kennedy et al., 2016; Warren et al., 2012). Unlike MTZ, AMIX does not undergo redox electron transfer, it is not mutagenic (Ballard et al., 2011; Hoffman et al., 2007; Warren et al., 2012), and furthermore it exhibits no cross-resistance with MTZ (Hoffman et al., 2007). Also important, AMIX targets are not found in humans, mitochondria or in aerobic and facultative anaerobes which utilize pyruvate dehydrogenase (PDH) in the catabolism of pyruvate (Hoffman, 2020). Pre-clinical studies show that AMIX is effective against *Clostridium difficile* and *Helicobacter pylori* in animal models (Hoffman et al., 2014; Warren et al., 2012) as well as oral anaerobic pathogens (Gui et al;., 2019; Gui et al., 2020, Gui et al., 2021; Hutcherson et al., 2017; Reed et al., 2018), anaerobic protozoan *Trichomonas vaginalis* (Jain et al., 2022), and exhibits no effect on the intestinal microbiota of healthy mice (Hoffman et al., 2014). However, very little is known about AMIX activity against *B. fragilis*.

PFOR is a major metabolic enzyme in the oxidative decarboxylation of pyruvate pathway in anaerobes and the best studied member of the KFOR superfamily (Gibson et al., 2016; Ragsdale, 2003). However, PFOR is not essential for *B. fragilis* growth *in vitro*, and though PFOR has been shown to be a major player in MTZ activation, and a target for AMIX activity, the lack of PFOR does not significantly increase resistance to MTZ (Diniz et al., 2004) nor does it alter susceptibility to AMIX (this study). This indicates that other metabolic pathways containing ThPP-binding enzymes that catalyze the cleavage or formation of carbon-carbon bonds of 2-ketoacids or 2-hydroxyketones (Gibson et al., 2016; Prajapati et al., 2022) may play important roles in MTZ and AMIX susceptibility. Nearly all ThPP-dependent enzymes can perform one-electron redox reaction steps that occur in the 2-electron process in oxidoreductases, and the low potential (~ −500mV) electrons generated can reductively activate pro-drugs such as MTZ (Chen et al., 2019; Gibson et al., 2016; Reed et al., 2012).

In addition to PFOR, *B. fragilis* encodes three other ThPP-binding KFOR members: two 2-ketoglutarate:ferredoxin oxidoreductases (KGOR); the *kor1AB* (BF638R_4321-4322) and the *kor2ABG* genes encompassed in the *kor2CDABEG* putative operon (BF638R_1660-1655), and the indolepyruvate ferredoxin oxidoreductase, *iorAB* (BF638R_1606-1605). *B. fragilis* Kor1AB and Kor2ABG subunits are orthologs to *Magnetococcus marinus MmOGOR*, KorAB (Chen et al., 2019), *Hydrogenobacter thermophilus HtOGORs*, KorAB, and ForDABGE (Yamamoto et al., 2006; Yamamoto et al., 2010; Yun et al., 2001; Yun et al., 2002), KGOR of *Thermococcus litoralis* α- and β-subunits (Mai and Adams 1996), and KGOR of *Tharnea aromatica* KorAB (Dörner & Boll, 2002) which catalyze the reductive carboxylation of succinyl-CoA to form 2-ketoglutarate (2-KG) in the reverse (reductive) TCA cycle. However, there is a paucity of information regarding the contribution of Kor1AB and Kor2CDAEBG in *B. fragilis* reverse reductive TCA cycle mode.

Interestingly, several studies have demonstrated that *B. fragilis* has a bifurcated TCA cycle containing the heme-dependent reductive branch oxaloacetate to succinate pathway, and the heme-independent citrate/isocitrate to 2-KG oxidative pathway branch but lacking reductive synthesis of 2-KG via succinate (Baughn & Malamy, 2002; Baughn & Malamy, 2003; Chen & Wolin, 1981; Harris & Reddy, 1977; Macy et al., 1975; Macy et al., 1978). However, there is strong evidence that *B. fragilis* and other anaerobic bacteria synthesize 2-KG from succinate via reductive carboxylation of succinyl-CoA. Using [1,4-^14^C]-succinate, or D-[U-^14^C]-glucose it was demonstrated that radiolabeled carbon was incorporated into glutamate via succinate but not via the citrate/isocitrate pathway (Allison & Robinson, 1970, Allison et al., 1979). In the closely related organism *Bacteroides thetaiotaomicron*, metabolomic studies monitoring carbon flux with radiolabeled [U-^13^C]-glucose also showed that glutamate and its precursor 2-KG are synthesized via succinate but not via citrate (Schofield et al., 2018). Assimilation of [U-^13^C]-acetate in *B. thetaiotaomicron* results in propagation of labeled citrate, but not to glutamate indicating that 2-KG is not formed by the oxidative branch (Schofield et al., 2018), though it is assumed it occurs in *B. fragilis* under heme-limiting conditions (Baughn & Malamy, 2002). Nonetheless, no genetic evidence nor enzymatic activity responsible for the reductive formation of 2-KG in *B. fragilis* has been described. In this regard, we have initiated studies to understand the regulation and metabolic role of Kor1AB and Kor2CDAEBG in *B. fragilis* central metabolism.

Pirins are a highly conserved redox-sensitive iron-binding proteins belonging to the functionally diverse cupin protein superfamily. Pirins contribute to the control and modulation of a diverse range of regulatory and metabolic activities in archaea, prokaryotes and eukaryotes. (Agarwal et al., 2009; An et al., 2004; Dunwell, et al., 2000; Dunwell et al., 2001; Dunwell, et al., 2004; Hihara et al., 2004; Lapik & Kaufman, 2003, Orzaez et al., 2001; Pang et al., 2004, Yoshikawa et al., 2004; Wendler et al., 1997). In the bacterium *Serratia marcescens*, pirin interacts with the pyruvate dehydrogenase (PDH) E1 subunit and modulates the PDH activity determining the direction of the pyruvate metabolism toward the TCA cycle or the fermentation pathway (Soo et al., 2007). The involvement of pirin in modulating central metabolism of prokaryotes was also demonstrated in *Streptomyces ambofaciens* (Talà et al., 2018) and *Aliivibrio salmonicida* (Hansen et al., 2012). In *Acinetobacter baumanii*, pirin regulates expression of adaptative efflux mediated antibiotic resistance (Young et al., 2023). In *Bacteroides thetaiotaomicron,* recombinant pirin exerts immunomodulating actions preventing the induction of pro-inflammatory interleukins in animal model of Crohn’s disease (Delday et al., 2019).

In this study we show that *B. fragilis* central metabolism activities are controlled, at least partially, by pirin proteins. Using a series of metabolic mutant strains and electron transport inhibitors, we show that resistance to MTZ involves a complex accumulation of mutations affecting metabolic pathways and electron transfer in energy conservation mechanisms. In addition, we carry out experiments to understand the metabolic role of Kor1AB and Kor2CDAEBG in *B. fragilis* central metabolism and show evidence that *kor2AEBG* genes are essential for growth and may play a role in the formation of 2-KG. Lastly, we demonstrate the novel ThPP-dependent enzyme cofactor inhibitor AMIX has antimicrobial activity against *B. fragilis* in an experimental model of intra-abdominal infection.

## MATERIALS and METHODS

### Strains and growth conditions

The bacterial strains and plasmids used in this study are listed in Table 1. *Bacteroides* species Strains were routinely grown on BHIS (brain heart infusion) supplemented with L-cysteine (1 g/liter), hemin (5 mg/liter), and NaHCO_3_ (20 mL of a 10% solution per liter). A chemically defined medium was formulated as follows: KH_2_PO_4_ (1.15 g/liter); (NH4)_2_SO_4_ (0.4 g/liter); NaCl (0.9 g/liter); L-methionine (75 mg/liter); MgCl_2_.6H_2_O (20 mg/liter); CaCl_2_.2H_2_O (6.6 mg/liter); MnCl_2_.4H_2_O (1 mg/liter); CoCl_2_.6H_2_O (1 mg/liter); resazurin (1mg/liter); L-cysteine (1 g/liter); hemin (5 mg/liter); and D-glucose (5 g/liter) or otherwise stated in the text. Final pH was 6.9. Rifampin (20 μg/mL), gentamicin (100 μg/mL), erythromycin (10 μg/mL), tetracycline (5 μg/mL), cefoxitin (25 μg/ml), 5-fluor-2’-deoxyuridine, FUdR, (200 μg/mL), or anhydrotetracycline, aTC (100 ng/ml) were added to the media when required. *E. coli* strains were routinely grown on Luria-Bertani (LB) media with appropriate antibiotics.

**Table 1.**
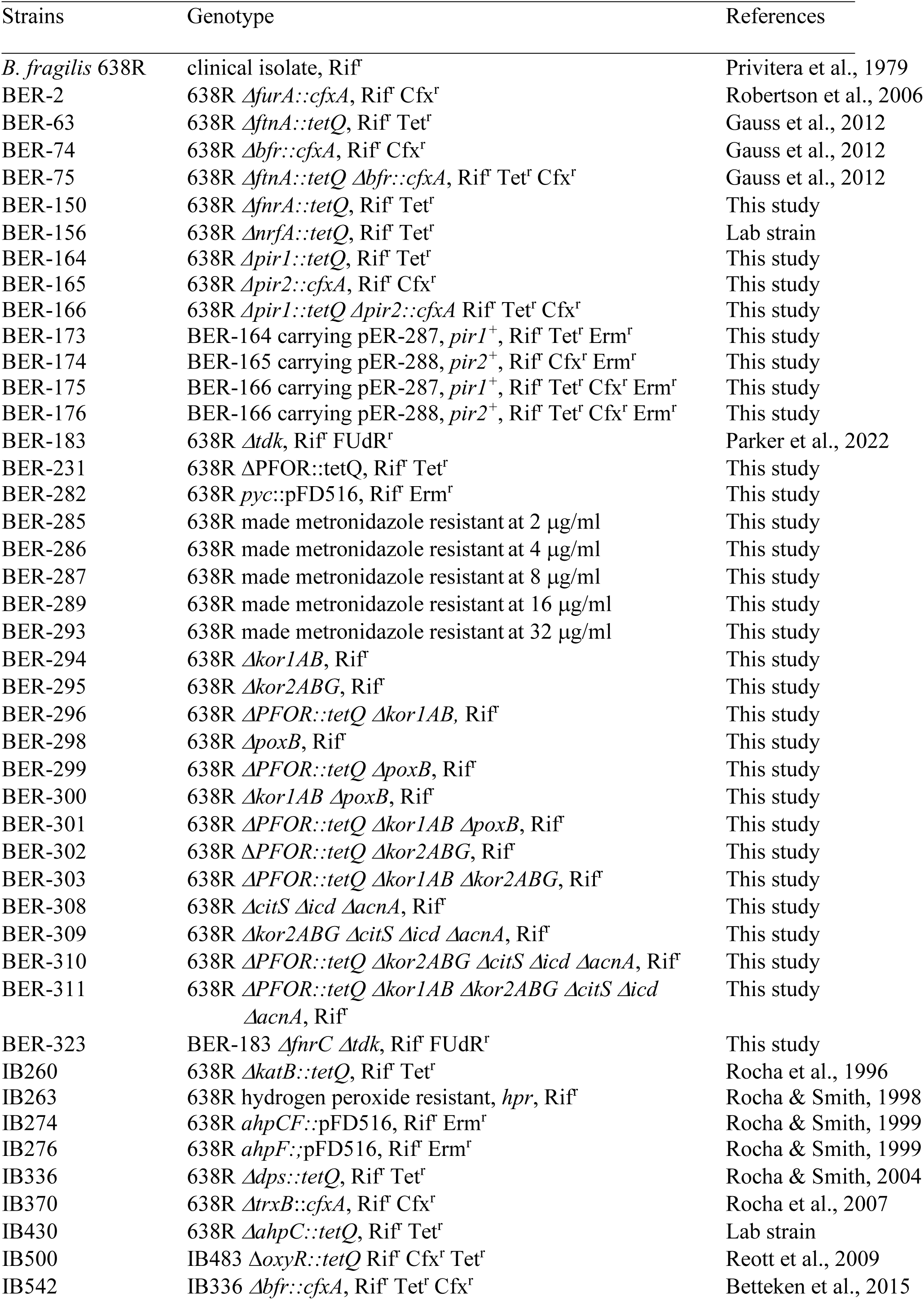
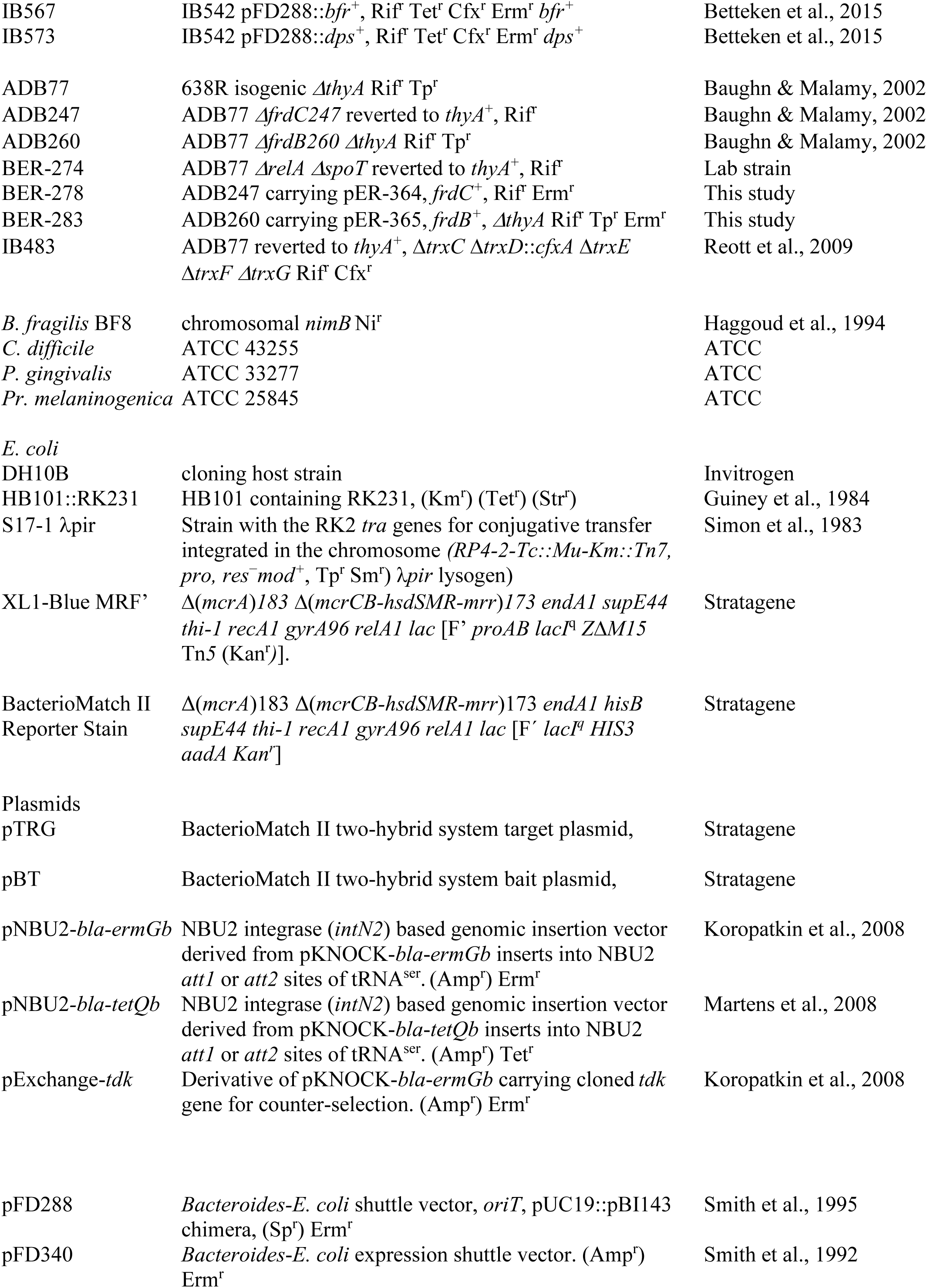
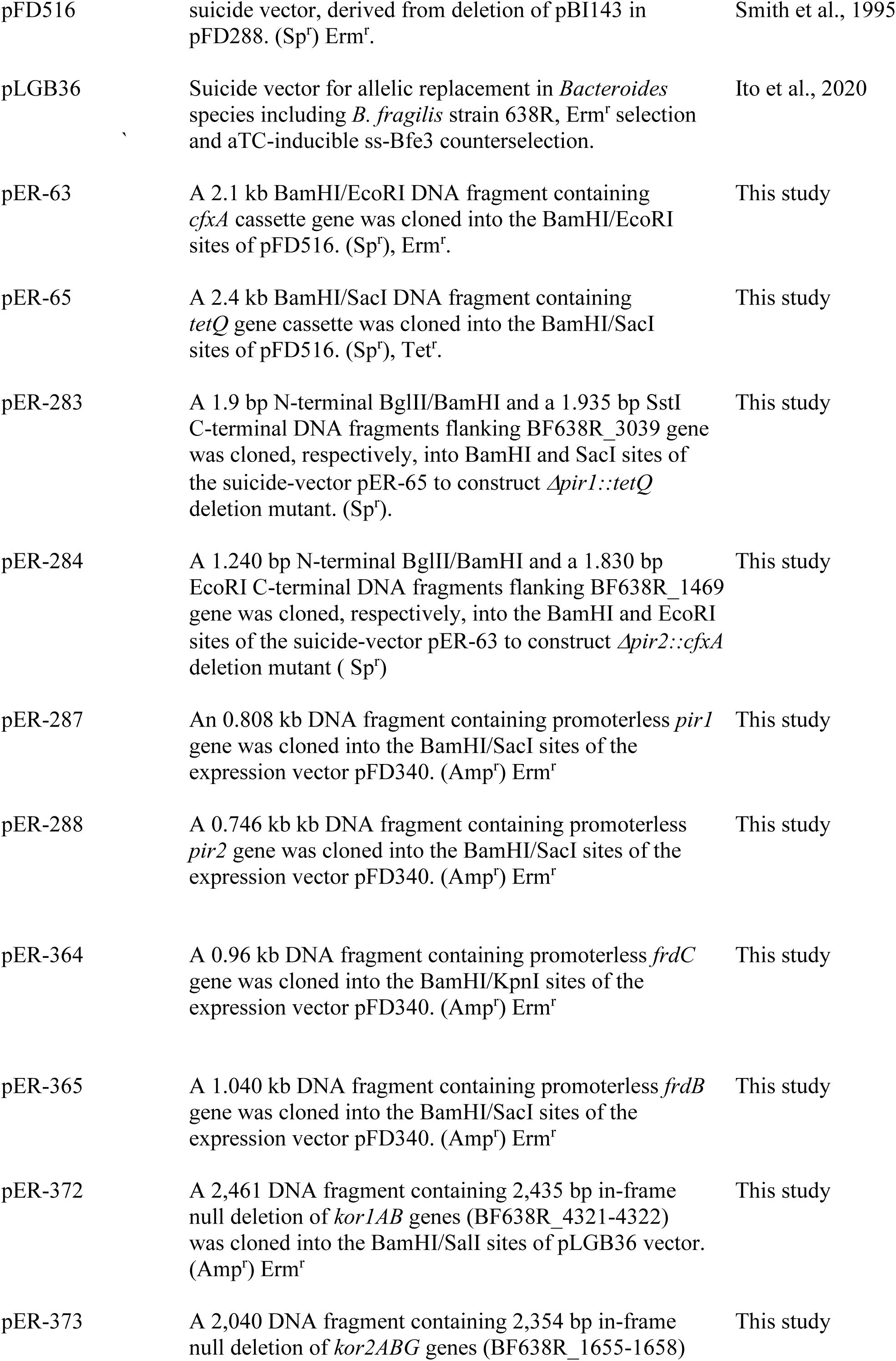
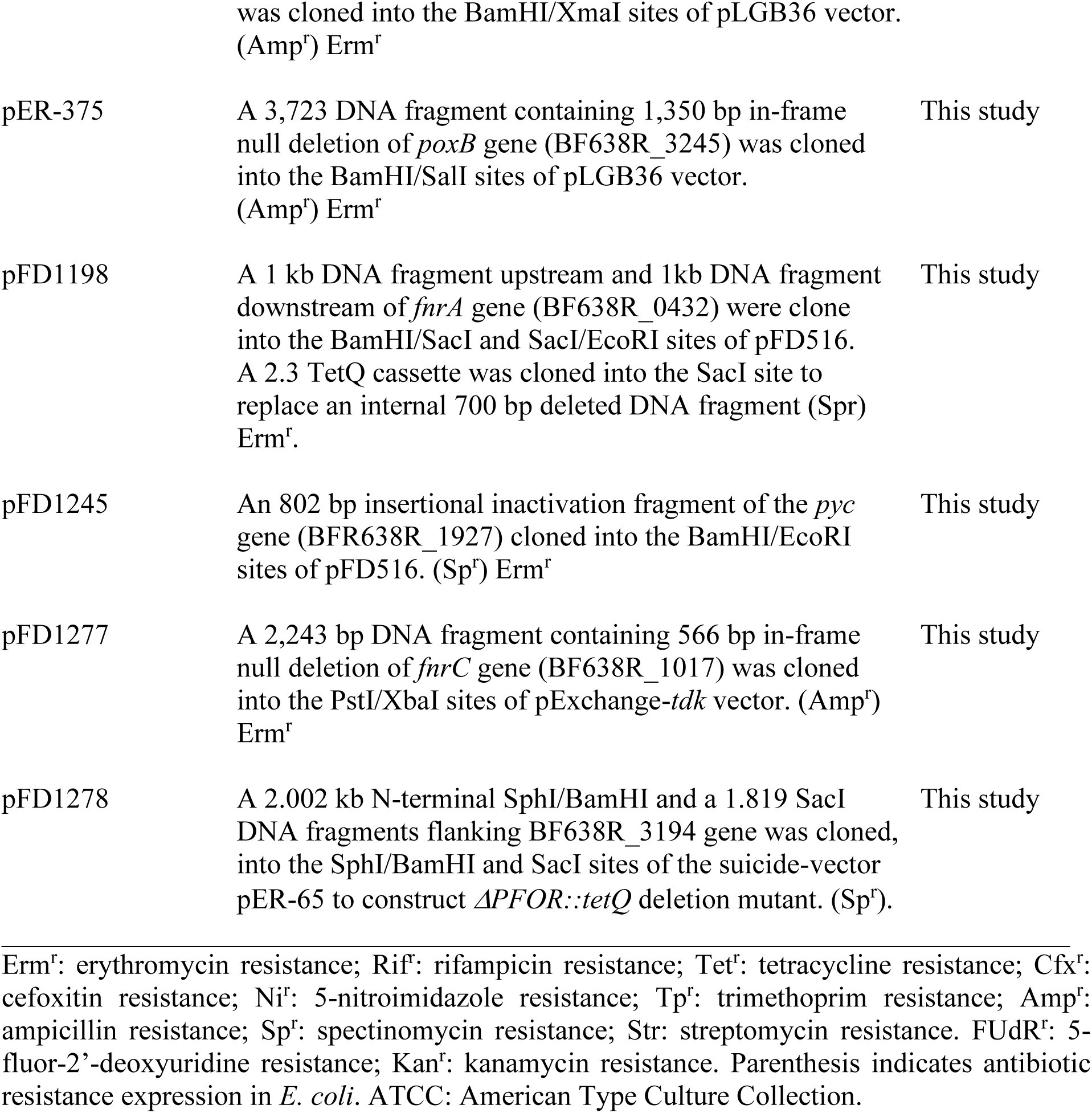
Bacterial strains and plasmids used in this study.

### Bacterial two-hybrid system assay

#### Screening partial chromosomal library for protein-protein interactions with Pir1 or Pir2

The chromosomal DNA from *B. fragilis* 638R was partially restricted with Sau3AI to obtain fragments average size within the range of 0.5 kb to 5 kb as previously described (Rocha & Smith, 1995). A genomic library was constructed by ligating the partial Sau3AI fragments into the unique BamHI site downstream of the RNAP α-subunit in the bacterial two-hybrid system (THS) target plasmid pTRG (BacterioMatch II; Stratagene, La Jolla, CA) and amplified in *E. coli* XL1-Blue MRF’ kan strain (Stratagene). The pirin 1, Pir1, (GenBank locus-tag BF638R_3039) or pirin 2, Pir2, (BF638R_1469) ORFs were amplified with primers described in Supplemental Table S1 and cloned in-frame with the Lambda repressor gene (*λcl*) under the control of the *lac-UV5* promoter in the bait plasmid pBT according to manufacturer’s instructions. The new constructs, pBT/Pir1 or pBT/Pir2 were cotransformed with the pTRG/genomic library into the *E*. *coli* THS reporter strain and plated on selective screening medium exactly as previously described (Robertson et al., 2006). The isolated two-hybrid system positive transformants were plated on dual-selective medium for validation as previously described (Robertson et al., 2006). The nucleotide sequence of the DNA fragment inserted into the TRG plasmid from each of the selected THS transformant was obtained using the pTRG forward primer and the deduced amino acid sequences were obtained.

#### Protein-protein interaction of Pir1 or Pir2 with PFOR and Zn-ADH

The entire ORF of PFOR and Zn-ADH were amplified by PCR and cloned in-frame with the RNA α-subunit of pTRG plasmid to construct pTRG/PFOR and pTRG/ADH, respectively. The pTRG/PFOR and pBT/Pir1, the pTRG/PFOR and pBT/Pir2, the pTRG/ADH and pBT/Pir1, or the pTRG/ADH and pBT/Pir2 plasmids were cotransformed into the *E*. *coli* THS reporter strain, respectively, and selected on non-selective medium, selective medium, and dual-selective medium exactly as previously described (Robertson et al., 2006).

#### Prediction of protein-protein interactions using computational modelling

Structure-based computational modelling of protein-protein interactions was used to assess the potential contributions of side-chain atoms in the interactions of Pir1 and Pir2 with PFOR, Zn-ADH, and with 140 enzyme units of the central metabolism and energy conservation of the TCA cycle, pyruvate metabolism, branched-chain amino acid aminotransferase (BCAAT), ThPP-binding enzymes, carboxylases/decarboxylases, oxidoreductases and dehydrogenases. The 3-dimensional structure of heteromeric protein-protein interactions were predicted using Alphafold2-Multimer https://colab.research.google.com/github/sokrypton/ColabFold/blob/main/AlphaFold2.ipynb (Evans et al., 2021; Jumper et al., 2021; Mirdita et al., 2022). Amino acid sequences of Pir1 and Pir2 along with other target enzymes were used as primary inputs for Alphafold2-Multimer. For models involving the interaction of Pir1 and Pir2 with PFOR, final models were also subjected to 2000 steps of energy minimization using an AMBER force field and in these specific cases the final energy relaxed models were used for interface analysis. Resulting models were analysed using Protein Interaction Z-Score Assessment (PIZSA) webserver http://cospi.iiserpune.ac.in/pizsa (Roy et al., 2019) using a distance threshold default 4Å to define interface residues contacts for the potential contributions of side chain atoms in the protein-protein interactions. A Z-score threshold ≥ 1.500 defined stable association. The interface area (Å^2^) and the buried surface area (Å^2^) of the interacting residues were calculated using Proteins, Interfaces, Structures and Assemblies (PDBePISA) webserver https://www.ebi.ac.uk/msd-srv/prot_int/pistart.html (Krissinel and Henrick, 2007).

#### HPLC analysis of short-chain fatty acids

Bacteria were grown to mid-logarithmic phase (OD_550nm_ to 0.3 – 0.4) or for 24h anaerobically in peptone yeast extract basal medium containing 0.5% D-glucose (PYG) prepared as described previously (Holdeman et al., 1977). Anaerobic mid-log cultures were split, and one half were exposed to atmospheric air for 1h or 24h in aerobic shaker incubator at 37 C. For iron restriction, 2,2’-dipyridyl (50 μM) was added to the medium. After pelleting cultures, the clear supernatants were passed through 0.22 μm filters. Samples were analysed for SCFAs using a BioRad HPLC organic acid system with AMINEX 87H, 300×7mm column with 5 mM sulfuric acid eluant at 0.6 ml/min, 65°C, with refractive index detector. Uninoculated media were used as blank and media background was subtracted except for glucose peak. The HPLC analysis were performed at the USDA Agricultural Research Service, Peoria, IL.

#### RNA extraction and RT-PCR analysis

Bacteria were grown in chemically defined medium supplemented with 100 μM ferrous sulphate or 50 μM 2,2’-dipyridyl to mid logarithmic phase anaerobically and exposed to atmospheric air for 1h. Total RNA was extracted from bacterial pellet using hot-phenol method as described previously (Rocha and Smith, 1997), and cleaned using RNeasy Mini kit (Qiagen, Valencia, CA) according to manufacturer instructions. RNA was DNAse treated using the Ambion DNAfree protocol (Ambion, Inc.). First strand cDNA synthesis was carried out from 1 μL total RNA at 1 μg/μL with random hexamer primers and Superscript III RT kit (Invitrogen Inc., Carlsbad, CA) according to manufacturer’s instructions. Real-time PCR quantification of each pirin mRNA was performed with 1 μL cDNA sample diluted 1:10 and forward and reverse primers described in Table S1. Real-time PCR efficiencies were performed for each primer set. The 16S rRNA was used as reference to normalize gene expression to a housekeeping gene. Relative expression values were calculated using the Pfaffl method (Pfaffl, 2001). Fold induction relative to the wild type in anaerobic conditions was determined for each gene using 16S RNA as the reference gene and all results were the average of at least two independent experiments with freshly isolated RNA.

#### Antibiotic susceptibility assays

The agar dilution method for minimal inhibitory concentration (MIC) determination and the disc inhibition assays were performed with BHI agar (20 ml/plate) containing heme (5 μg/ml). Overnight bacterial cultures in BHIS were diluted in PBS to a density approximately to 0.5 MacFarland scale. Five μl of suspension was applied on plates containing different concentrations of MTZ or AMIX. Disc inhibition was performed by spreading bacterial suspension with a swab. Five μl of 1 mg/ml MTZ solution or 10 μl of 1 mg/ml sterile AMIX in aqueous solution was applied on top of a sterile 6 mm disc paper. Plates were inoculated in duplicate. One set was incubated for 48h anaerobically at 37°C, and the other set was incubated in aerobically at 37°C for 24 before anaerobic incubation for 48h. Thymine (50 μg/ml) and sodium succinate (20 mM) were added to the media when required for growth of the *frdB* mutant, and the BF638R and isogenic ADB77 strains were used as control. Zone of inhibition around the disc was measured in mm. Electron transport system inhibitors or redox cycling agents were added to the medium when required as described in the text.

#### Construction of mutant strains

Deletion mutants with an antibiotic cassette replacing the gene internal DNA deleted fragment were constructed using pFD516 as suicide vector to mobilize mutated DNA fragments from *E. coli* DH10B into *B. fragilis* 638R for recombinant genetic exchange as described previously (Robertson et al., 2006; Rocha & Smith, 2004; Rocha et al., 2007). The forward and reverse primers used to PCR amplify DNA fragments to construct a deleted DNA fragment are shown in Supplemental Table S1. The construction of null deletion mutants in *B. fragilis* 638R was carried out using pLGB36 suicide vector for allelic replacement as described previously (Ito et al., 2020). Briefly, the pLGB36 constructs in *E. coli* S17-1 λpir strain were mobilized to *B. fragilis* 638R by biparental mating and transconjugants were selected on BHIS plates containing rifampicin (20 μg/ml), gentamycin (100 μg/ml) and erythromycin (10 μg/ml). A colony of erythromycin resistant first crossed-over recombinant strain was grown on 5 ml BHIS containing rifampicin (20 μg/ml), gentamycin (100 μg/ml), and without erythromycin, until OD_550 nm_ of 0.3-0.4. Then, 100 ng/ml aTC was added and the culture was incubated for approximately 2-3h to induce the ss-Bfe3 killer protein for counterselection. Ten μl aliquots were removed and spread on four plates of BHIS containing rifampicin (20 μg/ml), gentamycin (100 μg/ml), and aTC (100 ng/ml). After incubation for 3-4 days, colony PCR was performed using forward and reverse primer sets described in Supplemental Table S1 to identify transconjugants with chromosomal deletion fragment compared to the parent strain. Erythromycin susceptibility was performed to confirm loss of the suicide vector.

#### Intra-abdominal infection

All procedures involving animals followed the guidelines given by the National Research Council’s *Guide for the Care and Use of Laboratory Animals* (National Research Council, 2011) and approved by the Institutional Animal Care and Use Committee of East Carolina University. The rat tissue cage model of intra-peritoneal infection was performed exactly as described previously (Lobo et al., 2013) to test the *in vivo* efficacy of AMIX against *Bacteroides* 638R strain. Four groups of three Sprague-Dawley rats were infected with 4 ml of approximately 1 x 10^5^ CFU suspension in PBS into the tissue cage. One group was administered AMIX 20mg/kg once daily intraperitoneally (IP) at day 1 through day 7 post-infection. The second group received AMIX 0.5 mg injected intra-cage to obtain approximately 20 μg/ml final concentration. This expected intra-cage concentration corresponds to the AMIX concentration reached in rat serum receiving AMIX 20mg/kg/day via oral (Hoffman et al., 2014). Intra-cage injection of AMIX was administered once daily at day 1 through day 7 post-infection. The other two groups received saline administered IP or intra-cage as control. Intra-cage tissue fluid was aspirated at day 1, day 2, day 4, and day 8 post infection, serially diluted, and plated on BHIS. After 3 to 4 days of incubation in an anaerobic chamber at 37°C, colonies were counted and normalized to CFU/ml of tissue cage fluid. The limit of detection was 1 x 10^1^ CFU/ml.

## RESULTS

### The expression of *pir1* and *pir2* genes in response to oxygen and iron limitation

*B. fragilis* 638R contains two pirin-like proteins, Pir1 (BF638R_3039) and Pir2 (BF638R_1469). A genome transcription microarray of iron and heme regulated genes showed that both *pir1* and *pir 2* genes are up-regulated by iron limitation and following oxygen exposure (NCBI GEO DataSets GSE241210 and GSE241676, unpublished). Real Time RT-PCR using total RNA confirmed that *pir1* and *pir2* mRNAs were induced over 6-fold and 15-fold in the presence of oxygen, respectively (Fig. 1A), and over 5-fold and 8-fold under limiting iron conditions compared to parent strain, respectively (Fig. 1B). Furthermore, the *pir1* and *pir2* iron regulated expression is Fur-independent.

**Fig. 1.**
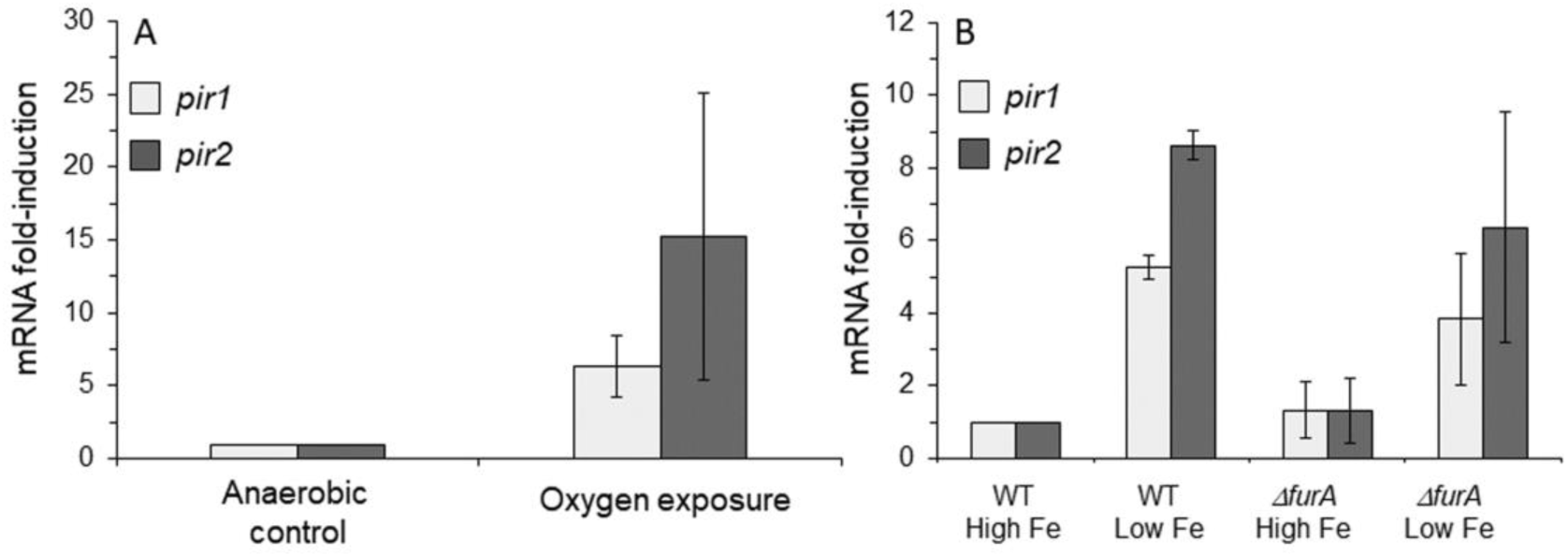
Fold-induction of *pir1* and *pir2* mRNA following (A) oxygen exposure or (B) growth in iron limiting conditions. In panel A, the BF638R parent strain was grown to mid-logarithmic phase in BHIS and exposed to atmospheric air for 1h. In panel B, the BF638R parent strain and its isogenic *ΔfurA* strain were grown to mid-logarithmic phase in defined medium with protoporphyrin IX supplemented with 100 μM FeSO_4_ (High Fe) or with 50 μM 2,2’-dipyridyl (Low Fe). For each condition, RNA was isolated and real-time RT-PCR was performed in triplicate. The 16S rRNA gene was used as an internal standard, and the results are expressed as fold induction relative to levels in the control condition. The values are means of fold induction compared to control from two independent experiments. The error bars indicate standard deviations.

A phylogenetic unrooted tree constructed from multiple amino acid alignments showed that the *Bacteroides* species Pir1 and Pir2 orthologs are grouped into two distinct clusters in a branch divergent from other members of the Bacteroidetes phylum (Supplementary file S1). The multiple amino acid sequence alignment revealed that the N-terminal domain residues His56, His58, His101, and Glu103 ligands of the iron centre of human pirin (Liu et al., 2013; Pang et al., 2004) are conserved in both Pir1 and Pir2 His58, His60, His102, and Glu104, respectively (Supplemental file S2).

### Two-hybrid system identifies Pir1 and Pir 2 protein-protein interactions with PFOR and a Zn-ADH

An *E. coli* THS assay (BacterioMatch II; Stratagene, La Jolla, CA) was used to screen a *B. fragilis* 638R partial Sau3A genomic library cloned in the target plasmid (pTRG) for expression of peptides forming protein-protein interactions with either Pir1 or Pir2 protein in the bait plasmid (pBT). Over fifty thousand co-transformed colonies were plated on selective THS media, and twelve colonies carrying pTRG/cloned insert interacting with pBT/Pir1 and 11 colonies carrying pTRG/insert interacting with pBT/Pir2 grew on selective media and confirmed by growth on dual-selective media. This indicated that peptides expressed from the pTRG/insert were positively interacting with Pir1 or Pir2. The deduced amino acid sequence in-frame with the C-terminal region of the RNAP α-subunit of each pTRG/insert interacting with pBT/Pir1 or pBT/Pir2 is shown in Supplemental file S3. One of these deduced peptide sequence showed homology to PFOR (BF638R_3194) interacting with Pir1, and two clones showed homology to a zinc-binding alcohol dehydrogenase, Zn-ADH (BF638R_1292), one interacting with Pir1 and the other with Pir2. To confirm these findings, the entire PFOR ORF (BF638R_3194) or Zn-ADH ORF was cloned in-frame with the RNA-α subunit into the pTRG vector. The new constructs, pTRG/PFOR and pTRG/ADH, were co-transformed with the pBT/Pir1 or pBT/Pir2 into the two-hybrid system reporter strain as described above. The THS assays showing protein-protein interaction of PFOR with Pir1 is shown in Fig. 2C, and Zn-ADH with Pir1 or Pir2 is shown in Supplemental file S4.

**Fig. 2.**
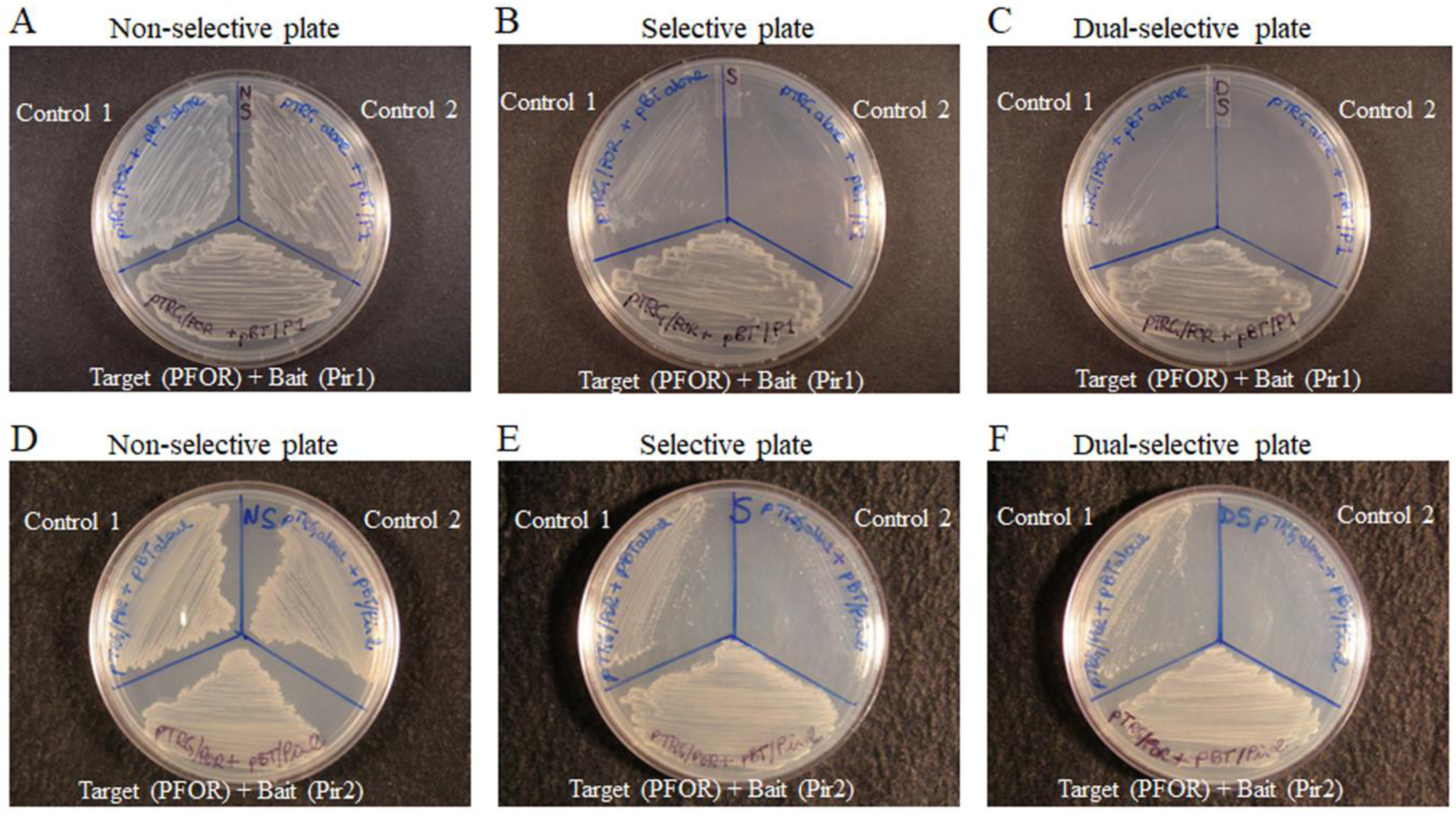
Bacterial two-hybrid system (THS) assay showing protein-protein interaction of PFOR with Pir1 (panels A, B, and C) and PFOR with Pir2 (panels D, E, and F). *E*. *coli* THS reporter strain cotransformed with pBT/Pir1 (bait) and pTRG/PFOR (prey) constructs are shown in panels A, B and C. *E*. *coli* THS reporter strain cotransformed with pBT/Pir2 (bait) and pTRG/PFOR (prey) constructs are shown in panels D, E and F. Bacteria were grown on a control non-selective plate (A and D), selective plate (B and E), and dual selective plate (C and F). Self-activation controls are *E*. *coli* THS reporter strain carrying the following constructs: Control 1, empty “bait” (pBT alone) cotransformed with loaded “prey” (pTRG/PFOR) in panels A to F; Control 2, loaded “bait” (pBT/Pir1) cotransformed with empty “prey” (pTRG alone) in panels A, B, and C, or loaded “bait” (pBT/Pir2) cotransformed with empty “prey” (pTRG alone) in panels D, E, and F See Materials and Methods for details on the bacterial two-hybrid system assay.

### AlphaFold2-Multimer-based modelling of protein-protein interactions between Pir1 or Pir2 with metabolic enzymes

To model protein-protein interactions of Pir 1 and Pir2 with PFOR, we used AlphaFold2-Multimer to predict 3D structures of Pir1:PFOR and Pir2:PFOR complexes. In agreement with the THS assay, the final relaxed AlphaFold2 models showed stable interactions of PFOR with Pir1 as judged by PISZA analysis (Z-score of 1.710 (stable association >1.5; Fig. 3A, Supplemental file S5A). Although, we did not obtain any genomic library clone indicating interactions of PFOR with Pir2, AlphaFold2 also predicted stable interactions of PFOR with Pir2 with a Z-score of 1.949 (stable association >1.5; Fig. 3B, Supplemental file S5A). To test this prediction, a THS assay was carried out using pTRG/PFOR and pBT/Pir2 constructs to co-transform *E. coli* THS reporter strain. Indeed, this experiment strongly supports that a direct interaction is formed between Pir2 and PFOR (Fig. 2F).

**Fig. 3.**
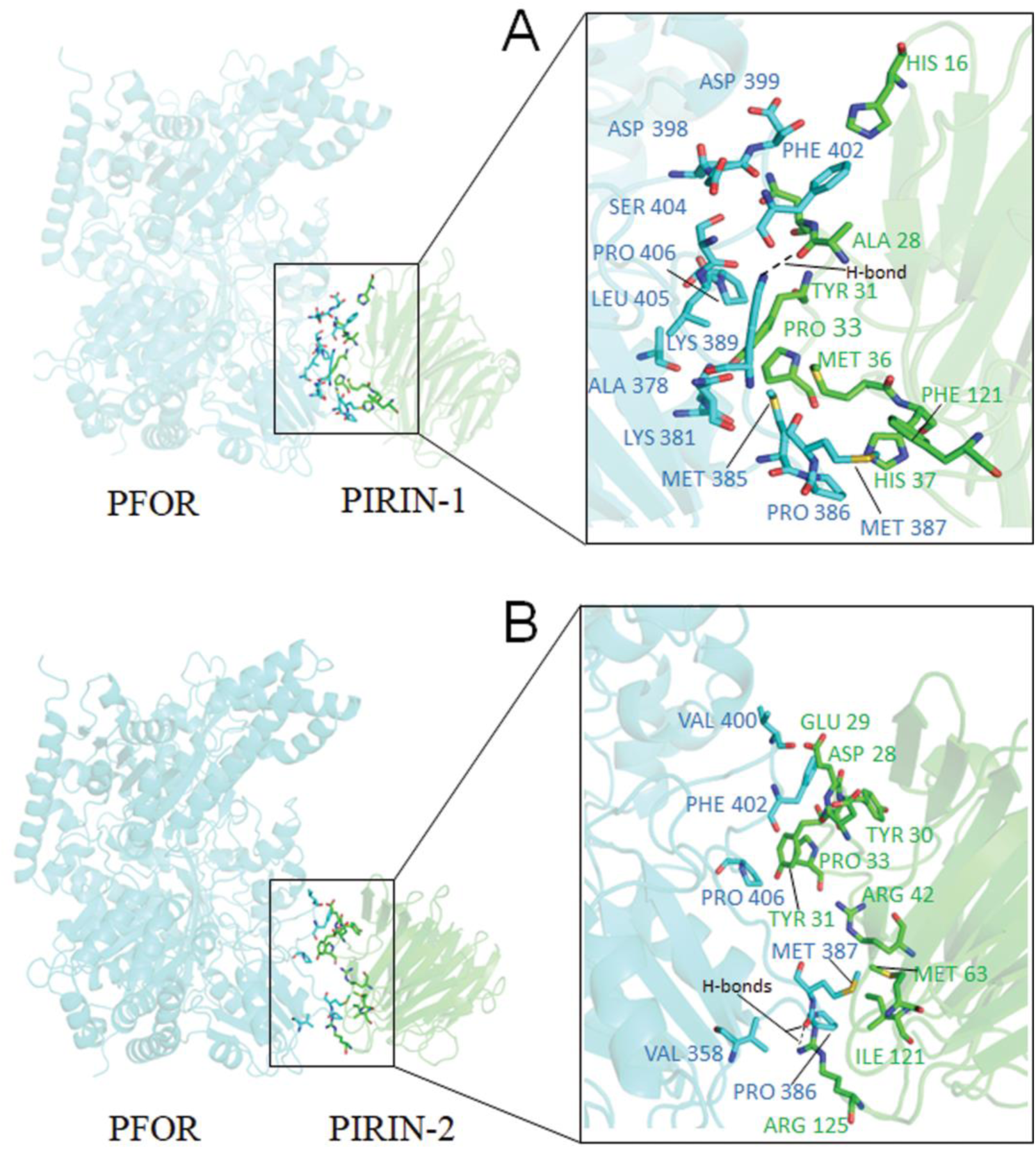
Ribbon cartoon diagram of AlphaFold2 Multimer 3D-structure modelling of the (A) Pirin1:PFOR and (B) Pirin2:PFOR complexes. Pirin proteins are shown in green and PFOR in cyan. Residue interaction pairs are drawn with stick models (Inset). Amino acids making contact in the protein-protein interface are depicted with font colour corresponding to each chain. A list of the amino acid interaction pairs and their relative buried surface area is shown in Supplemental file S5A. See Material and Methods section for additional details on the computational modelling methodology. Images were generated using PyMol molecular graphics system version 2.5.8 (Schrödinger, LLC).

These findings prompted us to perform additional 3D-structural modelling using Pir1 or Pir2 to understand if stable association with other metabolic enzymes of the central metabolism and energy conservation processes are also predicted. For this purpose, AlphaFold2-Multimer predictions using unrelaxed mode was used to broadly screen putative protein-protein interactions of Pir1 and Pir2 with enzymes involved in the biochemical pathways depicted in Supplemental file S5, including Zn-ADH. The resulting 3D structural models were then used to calculate the PISZA-interface Z-score and the buried surface area of the interacting amino acids with Pir1 or Pir2 (Supplemental file S5). In agreement with the THS screen (Supplemental file S4), stable interactions of Zn-ADH with Pir1 or Pir2 were predicted using this approach (Supplemental file S5A). Interestingly, 15 of the additional 140 tested enzymes were predicted to form stable protein-protein interactions with Pir1, 17 with Pir2, and 17 with both Pir1 and Pir2 (Supplemental file S5A), while 91 of the enzymes screened did not form stable protein-protein interactions with neither Pir1 nor Pir2 (Supplemental file S5B). Of the 49 proteins interacting with Pir1, Pir2 or Pir1 and Pir2, 32 enzymes belong to the oxidoreductase functional class, 9 are transferases, 2 are lyases, 2 are ligases, 2 proteins contain tetratricopeptide repeat (pfam13424 and pfam00515), 1 isomerase, and 1 conserved hypothetical protein. Interestingly, the amino acid residues of Pir1 and Pir2 that are predicted to coordinate stable interactions in 50% of the stable complexes are shared, (*i.e.*, Ala28, Asn29, Tyr31, Pro59 of Pir1 and Asp28, Glu29, Tyr31 of Pir2 (Supplemental file S2), indicating that a common binding site may be used.

### *pir1* and *pir2* deletion mutants have altered production of specific SCFAs

The results above suggested that Pir1 and Pir2 may have wide ranging interactions with key enzymes of central metabolism. To investigate this further we focused our investigation into the effects of *pir1* and *pir2* gene deletions on cellular metabolic activities by measuring fermentation of SCFA products and antimicrobial susceptibility to MTZ and AMIX. Analysis of the SCFA products of fermentation showed that in mid-log growth phase under anaerobic conditions the Δ*pir1* and Δ*pir2* mutants increase production of lactate and propionate compared to undetectable amount in the parent strain. Butyrate production increased over 3-fold in the *Δpir1 Δpir2* double mutant compared to parent strain. In contrast, the amounts of isovalerate and phenylacetate found in the parent strain were abolished in *Δpir1 Δpir2* double mutant strain in mid log anaerobic cultures (Supplemental file S6). When bacteria were grown to mid-log phase under iron limiting conditions, the amount of propionate decreased nearly 4-fold in the *Δpir1* single mutant, decreased 25-fold in the *Δpir2 single* mutant, and 2-fold in the *Δpir1 Δpir2* double mutant compared to parent strain. Under the same mid-log growth iron limiting conditions, there was no production of isobutyrate in the parent strain, but its production was observed in all mutant strains. Butyrate was produced only in the *Δpir1 Δpir2* double mutant strain. In contrast, isovalerate production was abolished in the mutants compared to the amount found in the parent strain. The amount of propionate found in the parent strain in cultures exposed to oxygen for 1h was abolished in the *Δpir1* mutant but not in the *Δpir2* mutant, and it was elevated over 2-fold in the *Δpir1 Δpir2* double mutant.

There was no significant difference in SCFA products in anaerobic cultures grown for 24 h except for minor production of butyrate in the *Δpir1 Δpir2* double mutant strain compared to its absence in the single mutants and parent strain. In cultures exposed to oxygen for 24h, butyrate was produced in the *Δpir1* and in the *Δpir1 Δpir2* double mutant strain but not in the *Δpir2* mutant or in the parent strain. Isovalerate was only produced in the *Δpir2* mutant but not in other strains in cultures exposed to oxygen for 24 h. Phenylacetate increased over 3-fold in the *Δpir1*, *Δpir2*, and *Δpir1 Δpir2* double mutant strains compared to the parent strain level. In iron-limiting cultures exposed to oxygen for 24 h, butyrate was produced in the mutant strains but not in the parent strain. Taken together, these findings indicate that Pir1 and Pir2 have a wide ranging modulatory effect on multiple metabolic pathways involved in the central carbon redox balance mechanisms and energy conservation processes for fermentation products formation.

### The effect of *Δpir1* and *Δpir2* mutations and constitutive expression of *pir1* and *pir2* genes in susceptibility to MTZ and AMIX

In *B. fragilis*, MTZ resistance is linked to redox cycling processes that occur during carbon redox steps that flows through its different fermentation pathways (Paunkov et al., 2023). To examine if the effect of Pir1 and Pir2 on cellular metabolism described above could alter susceptibility to MTZ and AMIX, MIC determination and disc inhibition assays were performed (Table 2 and Fig. 4). Disc diffusion assay showed that *Δpir1* and *Δpir* were significantly more sensitive to MTZ following oxygen exposure compared to the parent strain and anaerobic kept cultures (Fig. 4A). The lack of *pir* genes did not significantly affect sensitivity to AMIX in either condition (Fig. 4B). The MIC for MTZ and AMIX in the *Δpir1*, *Δpir2*, and *Δpir1 Δpir2* double mutant strains were not altered in anaerobic conditions or in cultures exposed to oxygen compared to the parent strain (Table 2).

**Fig. 4.**
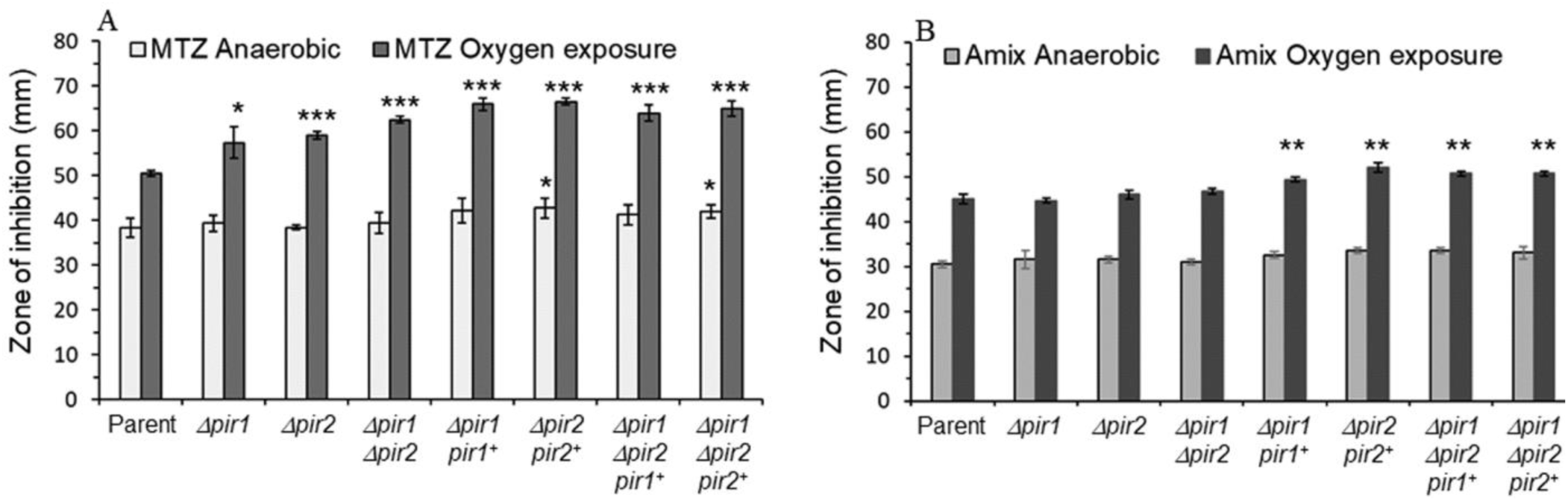
Disc diffusion assay sensitivity for A) metronidazole (MTZ) and B) Amixicile (AMIX). *B. fragilis* strains are depicted in each panel. Each bar represents the average zone of inhibition (mm) of at least three independent biological replicates. Vertical error bars denote standard deviation of the means from two independent experiments in triplicate. The significance of the *P* value was calculated between parent strain control and each mutant strain following an unpaired *t* test (parametric and two-tailed) two groups is shown above the horizontal bars. *p<0.05; **p<0.01; ***p<0.001.

**Table 2.** Agar dilution determination of minimal inhibitory concentration (MIC μg/ml) of metronidazole (MTZ) and amixicile (AMIX) for *Bacteroides species* and *B. fragilis* 638R mutant strains.

| Strains | MTZ<br>Anaerobic | MTZ O<sub>2</sub><br>exposure | AMIX<br>Anaerobic | AMIX O<sub>2</sub><br>exposure |
| --- | --- | --- | --- | --- |
| BF638R | 0.5 | 0.5 | 2 | 1 |
| BF638R isolated at 2 $\mu\text{g/ml}$ MTZ | 4 | 2 | 2 | 1 |
| BF638R isolated at 4 $\mu\text{g/ml}$ MTZ | 8 | 8 | 4 | 4 |
| BF638R isolated at 8 $\mu\text{g/ml}$ MTZ | 16 | 16 | 4 | 4 |
| BF638R isolated at 16 $\mu\text{g/ml}$ MTZ | 32 | 32 | 4 | 4 |
| *B. fragilis* ADB77 $\Delta\text{thyA}$ (638R isogenic) | 0.5 | 0.5 | 2 | 1 |
| *B. fragilis* BF8 | 16 | 2 | 2 | 1 |
| *B. fragilis* ATCC 25285 | 1 | 0.5 | 1 | 1 |
| *B. thetaiotaomicron* VPI 5482 | 1 | 1 | 1 | 1 |
| *B. vulgatus* ATCC 8482 | 1 | 0.5 | 1 | 0.0625 |
| *B. fragilis* 638R mutant strains |  |  |  |  |
| $\Delta\text{pir1}::\text{tetQ}$ | 0.5 | 0.5 | 2 | 1 |
| $\Delta\text{pir2}::\text{cfxA}$ | 0.5 | 0.5 | 2 | 1 |
| $\Delta\text{pir1}::\text{tetQ } \Delta\text{pir2}::\text{cfxA}$ | 0.5 | 0.5 | 2 | 1 |
| $\Delta\text{pir1}::\text{tetQ}/\text{pir1}^+$ | 0.25 | 0.125 | 2 | 0.5 |
| $\Delta\text{pir2}::\text{cfxA}/\text{pir2}^+$ | 0.25 | 0.125 | 2 | 0.5 |
| $\Delta\text{pir1}::\text{tetQ } \Delta\text{pir2}::\text{cfxA}/\text{pir1}^+$ | 0.25 | 0.125 | 2 | 0.5 |
| $\Delta\text{pir1}::\text{tetQ } \Delta\text{pir2}::\text{cfxA}/\text{pir2}^+$ | 0.25 | 0.125 | 2 | 0.5 |
| BF638R $\text{pir1}^+$ | 0.5 | 0.0625 | 2 | 1 |
| BF638R $\text{pir2}^+$ | 0.5 | 0.0625 | 2 | 1 |
| $\Delta\text{PFOR}::\text{tetQ}$ | 1 | 1 | 2 | 1 |
| $\Delta\text{kor1AB}$ | 0.5 | 0.5 | 2 | 1 |
| $\Delta\text{kor2ADBG}$ | 1 | 1 | 4 | 2 |
| $\Delta\text{PFOR}::\text{tetQ } \Delta\text{kor1AB}$ | 1 | 1 | 2 | 2 |
| $\Delta\text{poxB}$ | 1 | 0.5 | 4 | 1 |
| $\Delta\text{PFOR}::\text{tetQ } \Delta\text{poxB}$ | 1 | 1 | 4 | 1 |
| $\Delta\text{kor1AB } \Delta\text{poxB}$ | 1 | 0.5 | 2 | 1 |
| $\Delta\text{PFOR}::\text{tetQ } \Delta\text{kor1AB } \Delta\text{poxB}$ | 1 | 1 | 2 | 2 |
| $\Delta\text{PFOR}::\text{tetQ } \Delta\text{kor2ADBG}$ | 1 | 1 | 2 | 2 |
| $\Delta\text{PFOR}::\text{tetQ } \Delta\text{kor1AB } \Delta\text{kor2ADBG}$ | 1 | 1 | 4 | 2 |
| $\text{pyc}::\text{pFD516}$ | 0.5 | 0.5 | 1 | 1 |
| $\Delta\text{frdC247}$ | 0.25 | 0.25 | 1 | 0.5 |
| $\Delta\text{frdB260}$ | 0.25 | 0.0625 | 0.5 | 0.5 |
| $\Delta\text{frdC}/\text{frdC}^+$ | 0.5 | 0.25 | 2 | 1 |
| $\Delta\text{frdB260}/\text{frdB}^+$ | 0.5 | 0.25 | 1 | 1 |
| $\Delta\text{citS } \Delta\text{icd } \Delta\text{acnA}$ | 1 | 1 | 4 | 2 |
| $\Delta\text{kor2ADBG } \Delta\text{citS } \Delta\text{icd } \Delta\text{acnA}$ | 1 | 1 | 2 | 2 |
| $\Delta\text{PFOR}::\text{tetQ } \Delta\text{kor2ABG } \Delta\text{citS } \Delta\text{icd } \Delta\text{acnA}$ | 4 | 2 | 2 | 1 |
| $\Delta\text{PFOR}::\text{tetQ } \Delta\text{kor1AB } \Delta\text{kor2ADBG } \Delta\text{citS } \Delta\text{icd } \Delta\text{acnA}$ | 4 | 2 | 4 | 2 |
| Hydrogen peroxide resistant strain, *hpr* | 0.5 | 0.25 | 2 | 1 |
| $\Delta\text{katB}::\text{tetQ}$ | 0.5 | 0.25 | 2 | 1 |
| $\text{ahpCF}::\text{pFD516}$ | 0.5 | 0.25 | 2 | 1 |
| $\text{ahpF}::\text{pFD516}$ | 0.5 | 0.25 | 2 | 1 |
| $\Delta\text{ahpC}::\text{tetQ}$ | 0.5 | 0.25 | 2 | 1 |
| *ΔftnA::tetQ* | 0.5 | 0.25 | 2 | 1 |
| *Δbfr::cfxA* | 0.5 | 0.25 | 2 | 1 |
| *Δdps::tetQ* | 0.5 | 0.25 | 2 | 1 |
| *Δbfr::cfxA ΔftnA::tetQ* | 0.5 | 0.25 | 2 | 1 |
| *Δbfr Ddps* | 0.25 | 0.25 | 2 | 0.5 |
| *Δbfr::cfxA Δdps::tetQ/bfr<sup>+</sup>* | 0.25 | 0.25 | 1 | 0.5 |
| *Δbfr::cfxA Δdps::tetQ/dps<sup>+</sup>* | 0.25 | 0.125 | 1 | 0.5 |
| *ΔfnrA::tetQ* | 0.5 | 0.5 | 2 | 1 |
| *ΔnrfA::tetQ* | 0.5 | 0.5 | 2 | 1 |
| *ΔfnrC* | 0.5 | 0.5 | 1 | 1 |
| *ΔrelA ΔspoT* | 0.5 | 0.5 | 2 | 1 |
| *ΔtrxB::cfxA* | 0.25 | 0.25 | 1 | 0.5 |
| *ΔtrxC ΔtrxD::cfxA ΔtrxE ΔtrxF ΔtrxG* | 0.5 | 0.5 | 2 | 2 |
| *ΔtrxC ΔtrxD::cfxA ΔtrxE ΔtrxF ΔtrxG ΔoxyR::tetQ* | 0.5 | 0.25 | 2 | 0.5 |

Anaerobic: Inoculated plates were immediately incubated anaerobically. O<sub>2</sub> exposure: Duplicate plates were incubated for 16-20 h in aerobic incubator at 37° C prior to anaerobic incubation at 37° C.

The genetically complemented strains constitutively expressing *pir1* or *pir2* genes also showed a significant increase in MTZ and AMIX sensitivity following oxygen exposure but not in anaerobic plates as determined by disc inhibition assay (Fig. 4 A, B). In addition, the complemented *pir1* and *pir2* mutant strains showed 2-fold reduction in MTZ susceptibility (MIC 0.25 μg/ml) in anaerobic cultures, and 4-fold reduction (MIC 0.125 μg/ml) in cultures exposed to oxygen compared to the parent strain, respectively (Table 2). The MTZ MIC of the parent strain overexpressing *pir1* or *pir2* genes remained unaltered in anaerobic conditions (MIC 0.5 μg/ml) but when these strains were exposure to oxygen, they showed an 8-fold decrease in MTZ susceptibility (MIC 0.0625 μg/ml) compared to parent strain (Table 2). In addition, the complemented strains expressing *pir1* or *pir2* genes had no effect on AMIX MIC anaerobically and it caused a small reduction 2-fold (0.5 μg/ml) in oxygen exposed cultures compared to parent strain under the same conditions. The parent strain overexpressing *pir1* or *pir2* genes showed no changes in amixicile MIC anaerobically or following oxygen exposure (Table 2). Taken together, these findings support the role of Pir1 and Pir2 in altering metabolic activities though the specific pathways, potentially by forming protein-protein interactions with one or more enzymes that remain to be defined. Moreover, to investigate if changes in metabolic activities could also affect the mechanisms that might contribute to MTZ or AMIX susceptibility, a variety of cellular function mutants were used to carry out analysis of their antimicrobial susceptibility.

### Different functional metabolic mutants tested for susceptibility to MTZ and AMIX

A series of mutant strains available in our laboratory collection encompassing different metabolic and physiological functions such as: ThPP-binding 2-ketoacid oxidoreductases, carboxylases, TCA cycle enzymes, oxidative and redox stress responses, stringent response, nitrate reductase ortholog, anaerobic Fnr-like regulators, and iron-storage proteins listed in Table 1 were tested for MTZ and AMIX susceptibility. The genomic organizations and the deletion construct diagrams of four members of the ThPP-dependent 2-ketoacid oxidoreductases: the *PFOR*, *kor1AB*, and *kor2CDAEBG* which use ferredoxin as electron acceptor, and the putative inner membrane enzyme that catalyses the oxidative decarboxylation of pyruvate, *poxB*, used in this study are shown in Fig. 5. The results are presented in Table 2, Fig. 6, and Supplemental File S7. Of particular interest, the *Δkor2ABG*, *ΔpoxB* and *PFOR::tetQ* single mutant strains, the *ΔPFOR::tetQ Δkor2ABG* double mutant, the *ΔPFOR::tetQ Δkor1AB ΔpoxB* triple-mutant, and the ΔPFOR::tetQ *Δkor1AB Δkor2ABG* triple mutant strains showed 2-fold increase in MTZ resistance (MIC 1 μg/ml) compared to the parent strain (MIC 0.5 μg/ml). The *Δkor1AB* single mutant had no change in MTZ susceptibility (Table 2). However, the *ΔPFOR::tetQ Δkor2ABG ΔcitS Δicd ΔacnA* quintuple mutant had an 8-fold increase in MTZ resistance (MIC 4 μg/ml) anaerobically and 4-fold increase (MIC 2 μg/ml) following exposure to oxygen compared to parent strain (Table 2). The *ΔcitS Δicd ΔacnA* triple mutant showed a 2-fold MIC increase for MTZ and AMIX compared to the parent strain (Table 2).

**Fig. 5.**
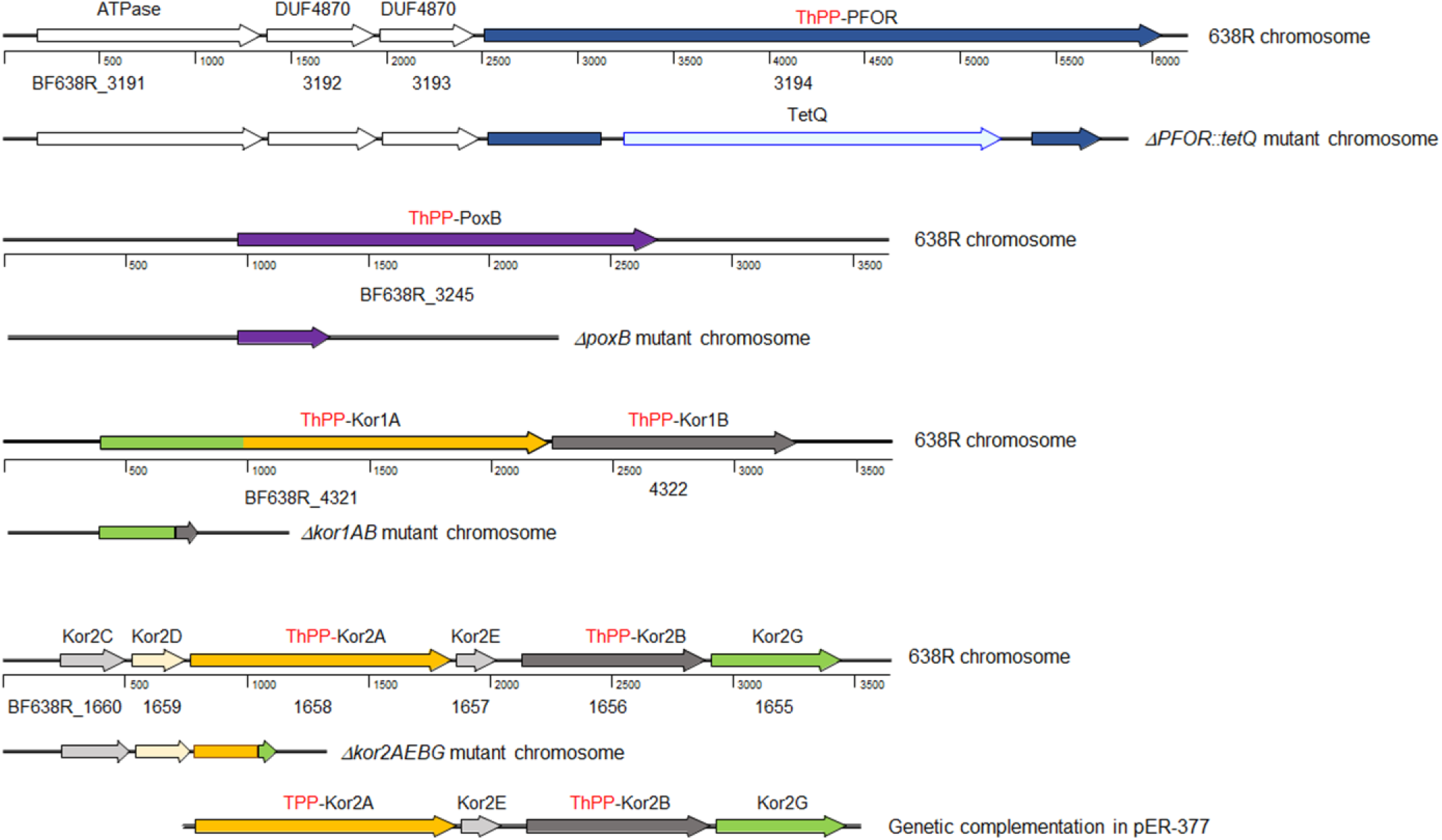
Schematic representation of *B. fragilis* 638R chromosomal regions for PFOR, PoxB, Kor1AB, and Kor2AEBG as shown in the panels. Each locus_tag is depicted below the respective deduced ORF symbolized by an arrow. The designation of the predicted peptide product is depicted above each open arrow gene region respectively. Arrow direction depicts the transcription orientation. Arrows filled with colour represent the functional annotation group assigned to PFOR (dark blue), PoxB (purple), KorA (orang), KorB (dark grey), KorG (light green), KorD (gold) or, KorC (light grey) orthologs, respectively. The deletion construct representation of each chromosomal region mutant is shown below the native chromosome region, respectively. The DNA fragment containing the promoterless *kor2AEBG* genes cloned into the expression vector pFD340 (pER-377) was used for genetic complementation studies. **ATPase**: predicted ATPase AAA+ superfamily. **DUF4870**: putative membrane protein of unknown function. **ThPP**: thiamine diphosphate cofactor. **PFOR**: pyruvate:ferredoxin oxidoreductase (GenBank accession number CBW23670), **PoxB**: putative pyruvate dehydrogenase (CBW23720), **Kor1A**: 2-ketoglutarate ferredoxin oxidoreductase subunit α (CBW24739), Kor1B: 2-ketoglutarate ferredoxin oxidoreductase subunit β (CBW24740), **Kor2A**: 2-ketoglutarate ferredoxin oxidoreductase subunit α (CBW22186), **Kor2B**: 2-ketoglutarate ferredoxin oxidoreductase subunit β (CBW22184), **Kor2C**: conserved hypothetical protein containing tetratricopeptide repeat (CBW22188), **Kor2D**: ferredoxin, 2-ketoglutarate-acceptor oxidoreductase subunit δ (CBW22187), Kor2E: hypothetical protein (CBW22185), and **Kor2G**: 2-ketoglutarate ferredoxin oxidoreductase subunit γ (CBW22183).

**Fig. 6.**
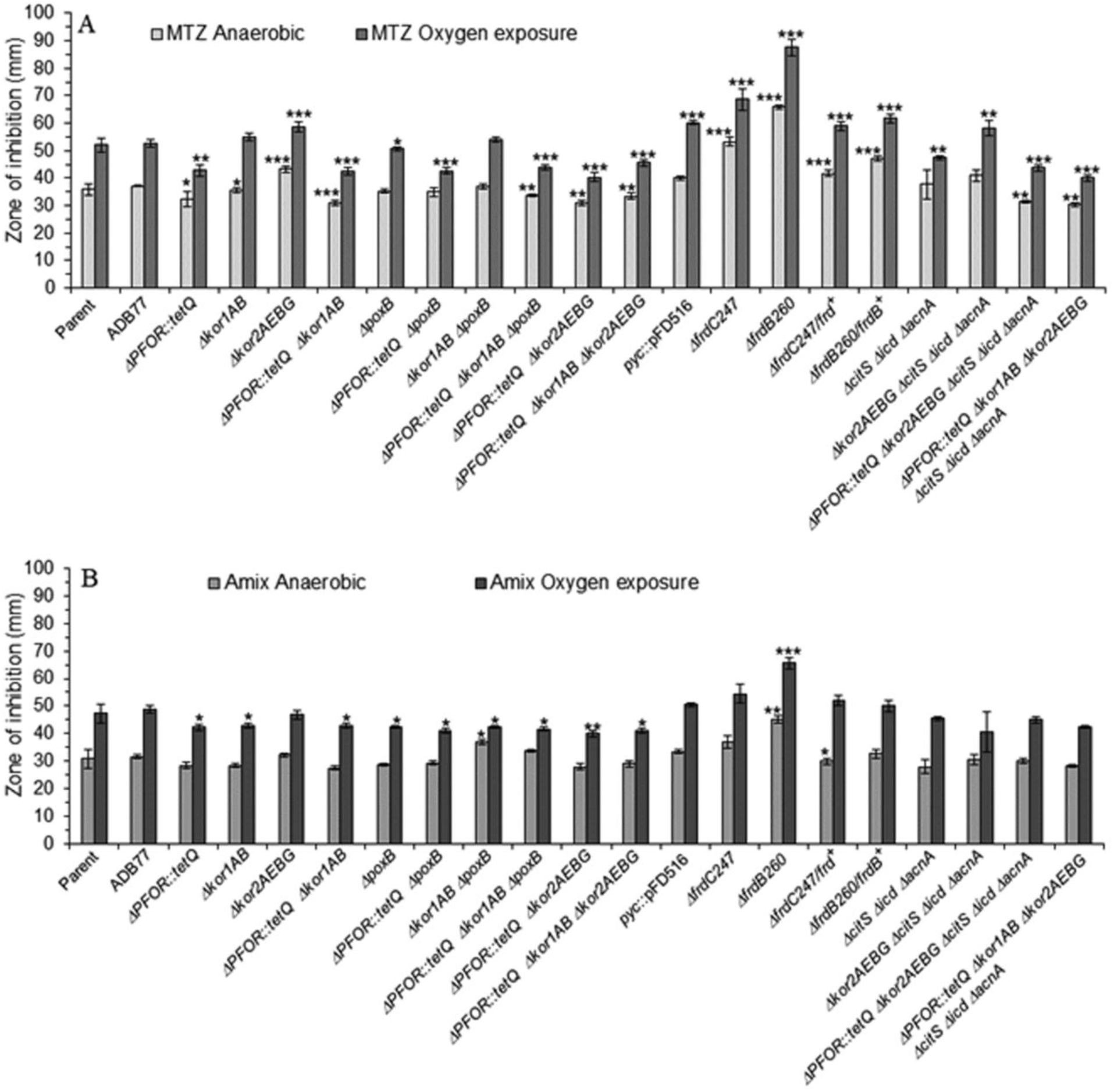
Disc diffusion assay sensitivity for A) metronidazole (MTZ) and B) Amixicile (AMIX). *B. fragilis* strains are depicted in each panel. Each bar represents the average zone inhibition (mm) of at least three independent biological replicates. Vertical error bars denote standard deviation of the means from two independent experiments in triplicate. The significance of the *P* value was calculated between parent strain control and each mutant strain following an unpaired *t* test (parametric and two-tailed) two groups is shown above the horizontal bars. *p<0.05; **p<0.01; ***p<0.001.

In contrast, the Δ*frdB* mutant had a 2-fold decrease in MTZ susceptibility (MIC 0.25 μg/ml) compared to parent strain. Interestingly, following oxygen exposure, the Δ*frdB* mutant had an 8-fold decrease in MTZ MIC (0.0625 ug/ml) (Table 2). The effect of the lack of the *frdB* subunit of the fumarate reductase on MTZ and AMIX susceptibility is also remarkedly seen in the disc inhibition assays (Fig. 6). The genetically complemented strains, *ΔfrdB260/frdB^+^* and *ΔfrdC/frdC^+^* brought MTZ and AMIX susceptibility close, but not identical, to the parent strain levels as determined by MIC and disc inhibition assays (Table 2 and Fig. 6). Deficiency in *pyc*, *PFOR*, *kor1AB, kor2AEBG,* or *citS icd acnA* genes did not significantly affect AMIX susceptibility (Table 2 and Fig 6B). A strain overexpressing *dps* gene showed significant increase in MTZ sensitivity compared to the parent strain but it did not affect AMIX susceptibility (Supplemental file S7B).

These findings appear to correlate deficiencies in the oxidative TCA cycle branch with an increase in MTZ resistance while deficiencies in the reductive TCA branch with an increase in MTZ susceptibility. These alterations in the oxidative and reductive balance of the central metabolism could ultimately lead to dysregulation of the NADH/NAD^+^ redox processes and cellular bioenergetics. To test this assumption, the inhibitors of *B. fragilis* NADH:electron acceptor transport coupling system (ETS) for fumarate reductase reduction of fumarate to succinate were used as described previously (Harris and Reddy, 1977).

### The effect of ETS inhibitors on MTZ and AMIX susceptibility

The following ETS inhibitors: acriflavine, rotenone, 2-heptyl-hydroxyquinoline-N-oxide (HQNO), and antimycin A were used as previously described (Harris & Reddy, 1977). Closantel, a halogenated salicylanilide antimicrobial whose mechanism of action is not completely understood but decouples oxidative phosphorylation and leads to inhibition of ATP synthesis (Rajamuthiah et al., 2014; Tran et al., 2016; Van Den Bossche et al., 1979; Williamson and Metcalf, 1967), and 2-Mercaptopyridine-N-oxide (2-MPNO), an NADH:fumarate reductase inhibitor (Turrens et al. 1999) were also included in this study. The ETS inhibitors were added to BHI plates at concentrations 2 to 4-fold lower than the amount needed to cause growth inhibition of *B. fragilis* 638R as indicated in the text. The BF638R strain susceptibility to both MTZ and AMIX were significantly increased in the presence of closantel, acriflavine, and HQNO under both anaerobic and oxygen exposed conditions (Fig. 7A, D). Antimycin A and rotenone did not cause significant changes in neither MTZ nor AMIX susceptibility. In the presence of 2-MPNO, there was a significant increase in MTZ susceptibility as determined by disc inhibition assays anaerobically and in oxygen exposed conditions while 2-MPNO did not alter AMIX susceptibility compared to control plates.

**Fig. 7.**
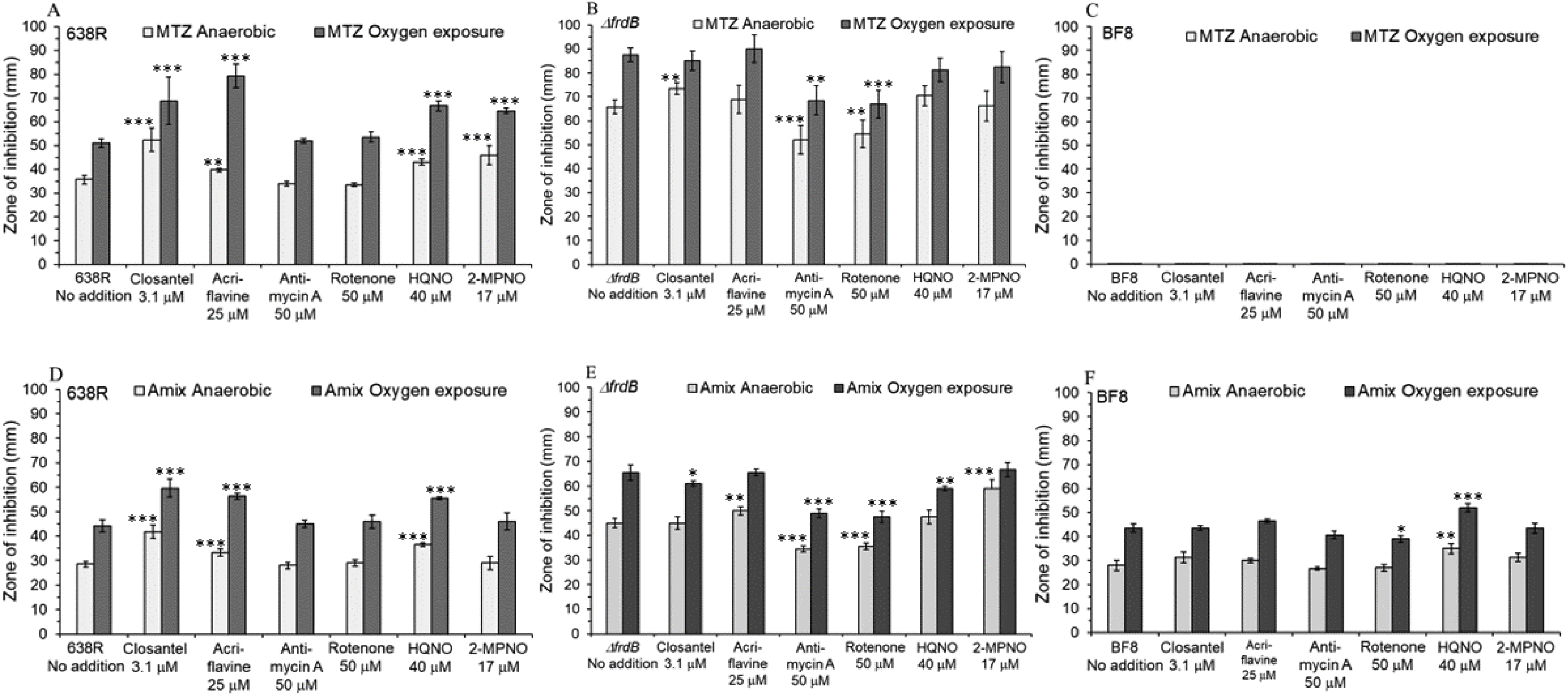
Disc diffusion assay sensitivity of *B. fragilis* strains to metronidazole (MTZ), panels A,B,C or for Amixicile (AMIX), panels D,E,F. A and D: Parent (*B. fragilis* 638R). B and E: *ΔfrdB* (derived from ADB77, isogenic BF638R *thy^−^* strain). C and F: *B. fragilis* BF8 strain (*nimB^+^*). Strains designations are depicted in each panel. For these experiments, 20 mM succinate and 50 μg/ml thymine were added into the BHIS media. Each bar represents the average zone of inhibition (mm) of at least three independent biological replicates. Vertical error bars denote standard deviation of the means from two independent experiments in triplicate. The significance of the *P* value was calculated between no addition control and each experimental group following an unpaired *t* test (parametric and two-tailed) two groups is shown above the horizontal bars. *p<0.05; **p<0.01; ***p<0.001.

In the presence of closantel, acriflavine, or HONO, the parent strain was significantly more sensitive to MTZ and AMIX compared to no addition control as determined by disc inhibition assays (Fig. 7A, D). A *ΔfrdB* strain was significantly more sensitive to MTZ and to AMIX than the parent strain with no addition of ETS inhibitors (Fig. 7A, B, D, E). However, sensitivity of the *ΔfrdB* strain to MTZ or AMIX, was not significantly altered in the presence of closantel, acriflavine, HQNO, or 2-MPNO but it was more sensitive in the presence of antimycin A and rotenone (Fig. 7B, E). The presence of ETS did not significantly affect the sensitivity to AMIX of the MTZ resistant strain BF8 strain compared to control except for an increase in AMIX sensitivity in the presence of HQNO (Fig. 7C, F).

### No crossed MTZ resistance and AMIX susceptibility

There is no report of AMIX resistant mutant strains. and this led us to test whether resistance to MTZ could alter susceptibility to AMIX. MTZ resistant strains were obtained by isolating random strains grown in increasing MTZ concentrations at 2, 4, 8, and 16 μg/ml. The findings showed that BF638R_(2 μg/ml)_ (MIC 4 μg/ml) showed no increase in AMIX resistance compared to the parent strain (Table 2). The BF638R_(4 μg/ml)_ (MIC 8 μg/ml), BF638R_(8 μg/ml)_, (MIC16 μg/ml), and BF638R_(16 μg/ml)_ (MIC 32 μg/ml) strains only showed 2-fold increase in AMIX MIC (4 ug/ml) (Table 2). In Fig. 8A, C, a comparison is shown of MTZ resistant strains to MTZ and to AMIX using disc inhibition assays. In addition, the BF8 strain carrying a *nimB* gene showed no change in AMIX susceptibility as determined by agar dilution and disc diffusion assays (Table 2, Fig. 8B, D).

**Fig. 8.**
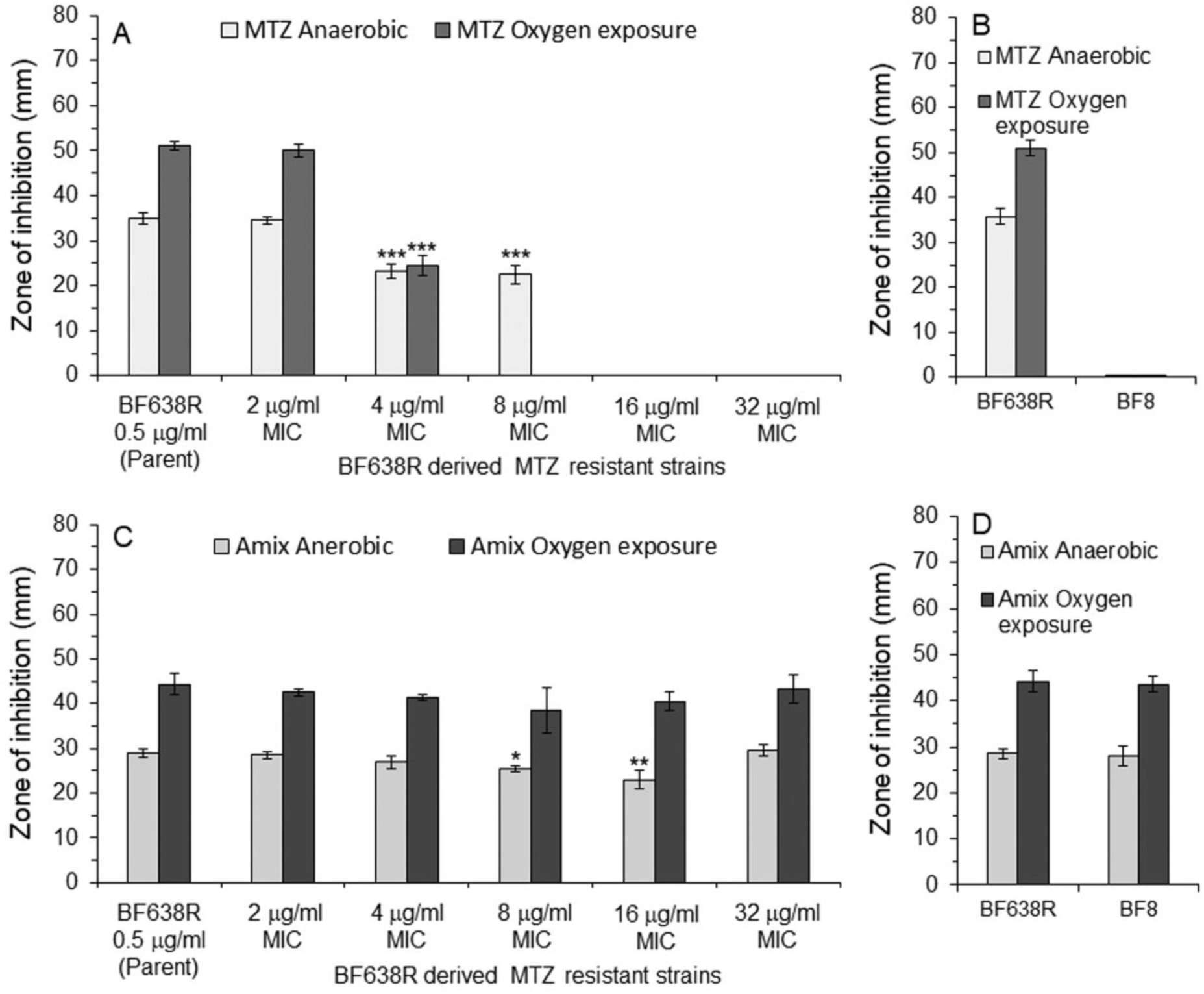
Disc diffusion assay sensitivity for A) metronidazole (MTZ) and B) Amixicile (AMIX). *B. fragilis* strains are depicted in each panel. Each bar represents the average zone inhibition (mm) of at least three independent biological replicates. Vertical error bars denote standard deviation of the means from two independent experiments in triplicate. The significance of the *P* value was calculated between parent strain control and each MTZ resistant strain following an unpaired *t* test (parametric and two-tailed) two groups is shown above the horizontal bars. *p<0.05; **p<0.01; ***p<0.001.

Additional evidence indicating that MTZ resistance does not interfere with AMIX susceptibility was the addition of ellagic acid, a polyphenolic compound that is a competitive inhibitor of *E. coli* nitroreductase, NfsA, *in vitro* (Chen et al., 2022). Ellagic acid added at 50 μM and 100 μM strongly increased resistance to MTZ while it had no apparent effect on AMIX susceptibility as determined by disk diffusion assay (Supplemental file S8). However, it remains to be defined how polyphenol compounds may affect MTZ resistance in *B. fragilis*.

### AMIX has antimicrobial activity against *B. fragilis* in an *in vivo* intra-abdominal infection model

The tissue cage model of intra-abdominal infection was used to examine if AMIX would have antimicrobial activity against *B. fragilis in vivo.* After inoculating *B. fragilis* 638R into the intra-abdominal tissue cage, the CFU/mL counts decreased by nearly 3 log-fold at day 4 post-infection and over 4 log-fold at day 8 post-infection in rats receiving AMIX IP compared to the CFU/mL of untreated rats receiving saline (Fig. 9A). However, the viability of *B. fragilis* in rats receiving intra-cage administration of AMIX was completely lost (below detection) at day 4 post-infection, and the CFU counts remained undetectable at day 8 post-infection compared to untreated control receiving saline (Fig. 9B). Taken together, these findings show that AMIX, a narrow spectrum antimicrobial designed to replace and inhibit thiamine-binding in ThPP-binding oxidoreductases, such as PFOR, is a potential alternative antimicrobial for *B. fragilis* infection.

**Fig. 9.**
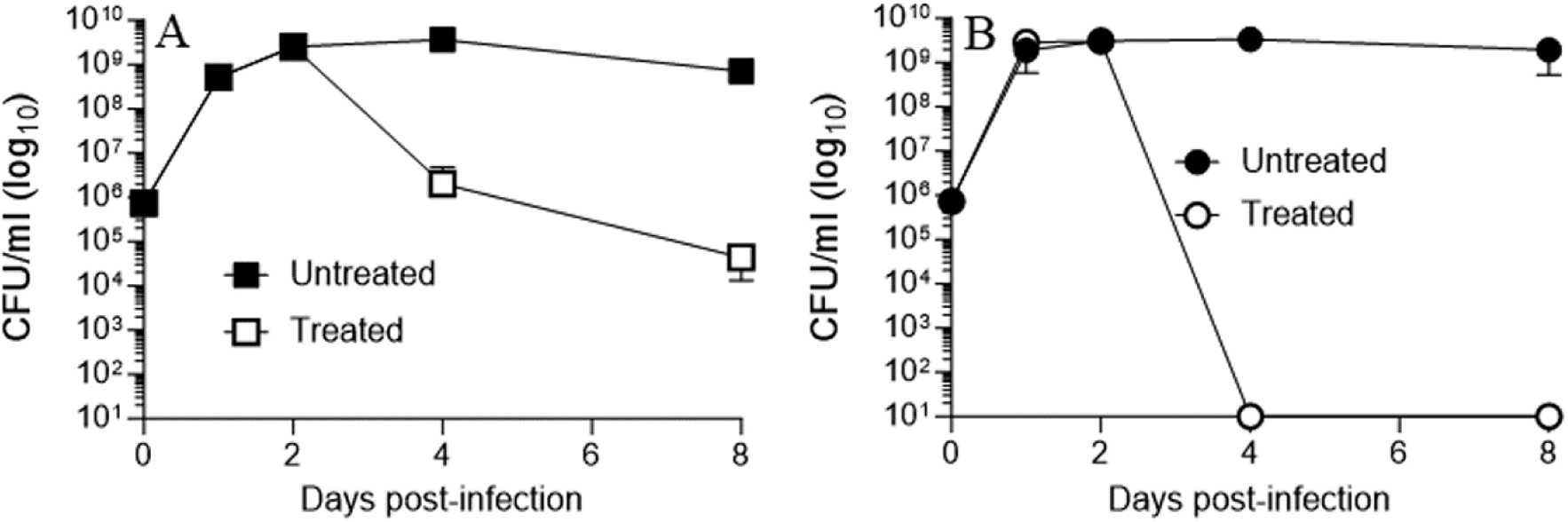
Survival of *B. fragilis* 638R in rat tissue cage infection following AMIX treatment at 20 mg/Kg/day via intraperitoneal (A) or 0.5 mg/intra-cage/day (B). AMIX administration started at day 1 through day 7 post-infection. Bacteria were grown overnight in BHIS medium and diluted in PBS to approximately 1 × 10^5^ CFU/mL. Four milliliters of the suspension were inoculated into the intraperitoneal tissue cage. Fluid samples were aspirated at time points for CFU counts as described in Materials and Methods.Tissue cage fluid was aspirated at day 1, 2, 4 and 8 post-infection. Data are expressed as the mean CFU per milliliter of intra-abdominal tissue cage fluid from three rats. The standard errors of the means (SEM) are denoted by vertical error bars. Detection limit of 1 × 10^1^ CFU/mL.

However, since lack of PFOR did not significantly alter *B. fragilis* growth nor susceptibility to AMIX *in vitro* as shown above in Table 2 and Fig. 5, experiments were carried out to find out if other members of the ThPP-binding 2-ketoacid oxidoreductases might play an essential role in *B. fragilis* anaerobic metabolism and physiology as described below.

### The ThPP-binding 2-ketoglutarate ferredoxin oxidoreductase subunits Kor2AEBG are essential for *B. fragilis* growth

Deletion of the *kor2AEBG* genes contained in the putative *kor2CDAEBG* operon caused a severe growth defect in rich media and completely abolished in minimally defined media (Fig. 10A, C). The *ΔPFOR::tetQ*, *ΔkorAB*, and *ΔpoxB*, single mutants, the *ΔPFOR::tetQ ΔpoxB*, *ΔkorAB ΔpoxB*, and *ΔPFOR::tetQ ΔkorAB* double mutants, and the *ΔPFOR::tetQ ΔkorAB ΔpoxB* triple mutant strains did not show significant growth defects indicating that the deficiency of *kor2AEBG* genes alone are responsible for the growth defect. The genetic complementation of the strains carrying the *Δkor2AEBG* deletion with the *kor2AEBG* native genes completely restored growth to parent strain levels in both BHIS media and in minimally defined media (Fig. 10B, D). However, the role of the *kor2AEBG* genes in the reductive formation of 2-KG *B. fragilis* TCA cycle is not well defined. In addition, we cannot rule out that *kor1AB* genes could be involved in the synthesis of 2KG because the *kor2AEBG* was shown to play a multifunctional role.

**Fig. 10.**
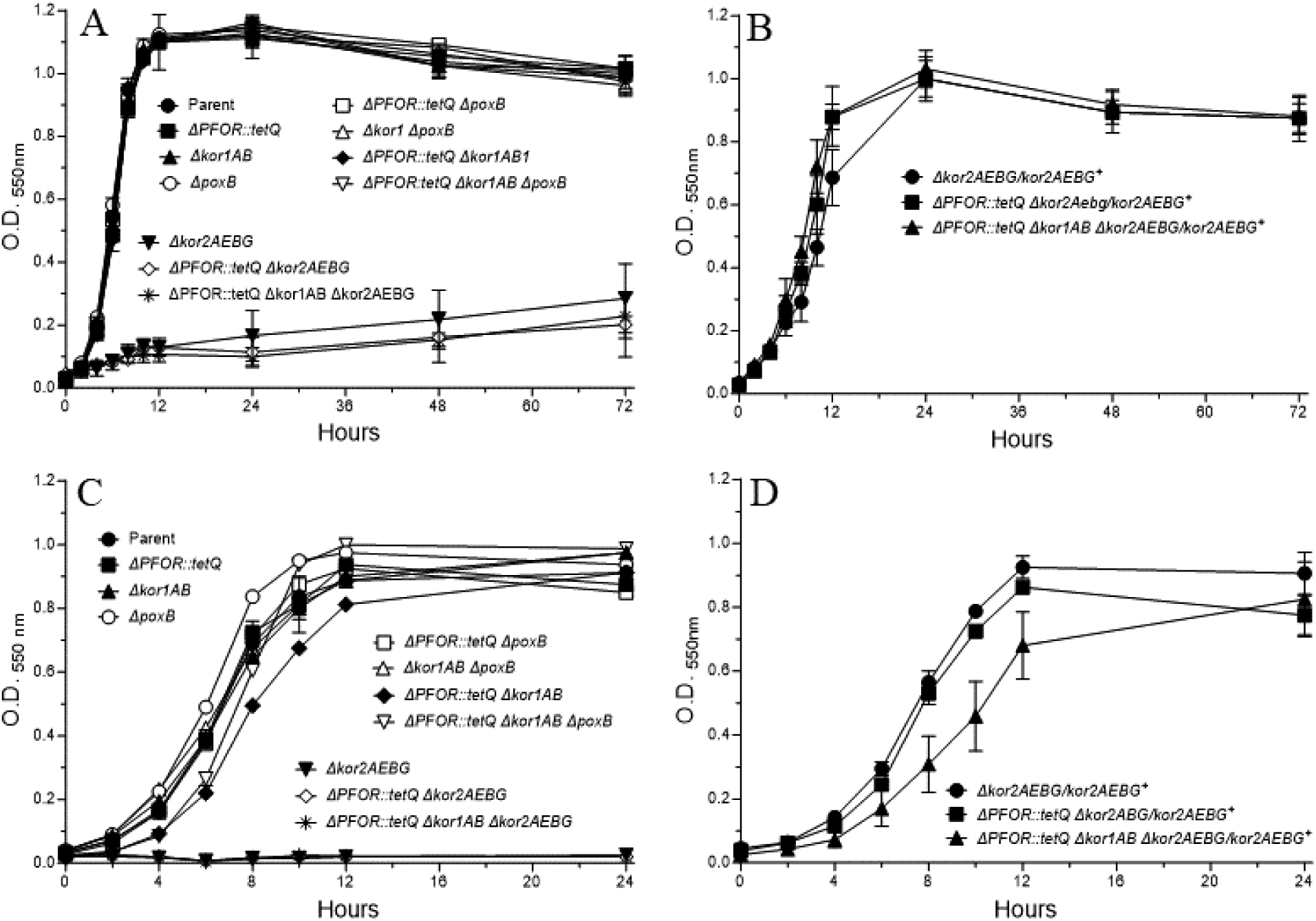
Growth of *B. fragilis* mutant strains in supplemented brain-heart infusion (BHIS) media (Panels A and B), or in chemically defined media with glucose (DM) (panels C and D). Strain designations are depicted for each panel. Media preparations and compositions are described in Materials and Methods section. Panels B and D show the *Δkor2AEBG* genetic complemented strains with pER-377 grown in BHIS (B) or defined medium (D), respectively.

### Supplementation with dimethyl-2-ketoglutarate (dM-2KG) restores growth of the *Δkor2AEBG* mutant in complex media but not in chemically defined media

Previous work with *B. thetaiotaomicron* has demonstrated that dM-2KG, a membrane-permeable precursor of 2-KG is transported across the cytoplasmic membrane and is hydrolysed to 2-KG (Schofield et al., 2018). Therefore, we hypothesized that dM-2KG could rescue the *Δkor2AEBG* phenotype. When 30 mM dM-2KG was added to BHIS media it stimulated the growth of the *Δkor2AEBG* single mutant and the *Δkor2AEBG ΔcitS Δicd ΔacnA* quadruple mutant strains compared to no addition culture control (Fig. 11A,B). The effect of dM-2KG from 1 mM to 40 mM on the growth of the *Δkor2AEBG* strain in BHIS media is shown in Supplemental file S9. In chemically defined media there was no growth stimulation of the *Δkor2AEBG* or the *Δkor2AEBG ΔcitS ΔacnA Δicd* quadruple mutant strain in the presence of 10 mM dM-2KG compared to no addition control (Fig. 11C,D), but in defined media containing 1% yeast extract, addition of 10 mM or 20 mM dM-2KG stimulated growth of the *Δkor2AEBG* mutant and the *Δkor2AEBG ΔcitS ΔacnA Δicd* quadruple mutant strain compared to control (Fig. 11E,F,G). No growth defect was observed in the *ΔcitS ΔacnA Δicd* triple mutant strain compared to the parent strain but growth was strongly inhibited at 30 mM dM-2KG in chemically defined media supplemented with yeast extract (Fig. 11H). The addition of 10 mM dM-2-KG in defined media containing 1% tryptone did not restore growth of the *Δkor2AEBG* mutant strain compared to no addition control (Supplemental file S10A, E, F). Addition of 30 mM dM-2KG into the minimally defined media or minimally defined media with 1% Tryptone was also highly inhibitory to growth (Supplemental file S10C, G). In contrast, it was required 1 mM dM-KG for *B. thetaiotaomicron* to synthesize L-glutamate or L-glutamine and to halt growth in defined media (Schofield et al., 2018). Since the membrane-permeable ester dM-2-KG is cleaved by intracellular esterases to form 2-KG in *E. coli* (Doucette et al., 2011), we cannot rule out that transport deficiency or low “esterase” activity in *B. fragilis* might account for these variations.

**Fig. 11.**
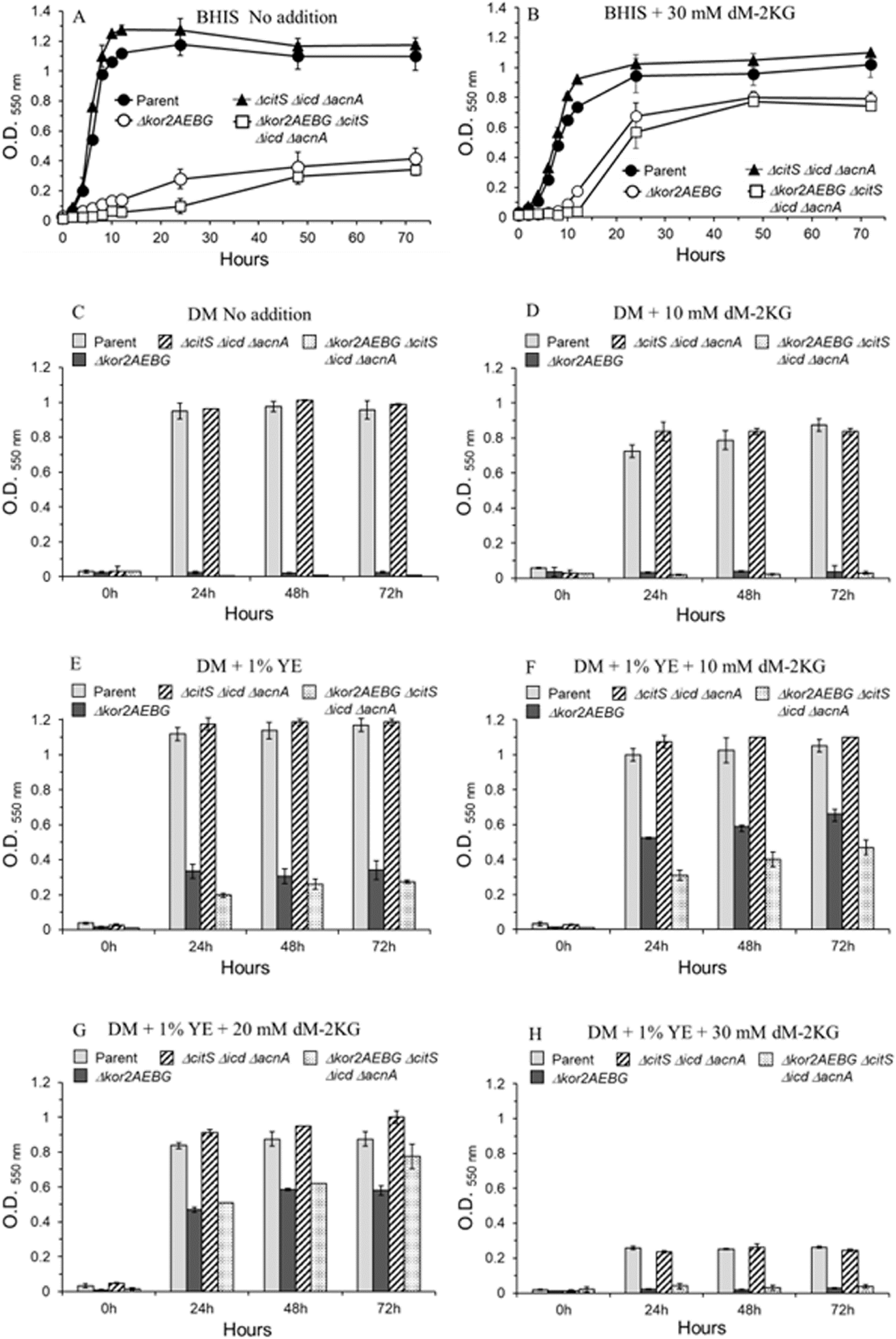
Growth of *B. fragilis* mutant strains in supplemented brain-heart infusion (BHIS) media (Panels A and B), or chemically defined media with glucose (DM) (panels C to H). DM supplemented with 1% yeast extract (YE) (Panels E to H). Dimethyl-2-ketoglutarate (dM-2KG) was added as indicated in the panels (Panels B, D, E to F). Strain designations are depicted for each panel. Media preparations and compositions are described in Materials and Methods section.

Overall, our results suggest that Kor2AEBG might have dual-function as determined by a) growth deficiency in rich media is rescued by dM-2KG supplementation suggesting a 2KG precursor by-pass requirement for this enzyme, b) an unknown metabolic function since addition of L-glutamate, L-glutamine, or tryptone (a glutamate-rich peptide source) did not compensate for the lack of the *kor2AEBG* genes (Supplemental file S10I, J, K, L) in defined media suggesting that formation of 2-KG as precursor for the synthesis of L-glutamate is not the only metabolic function of Kor2AEBG.

To better understand how Kor2AEBG functions in *B. fragilis* metabolism, we supplemented chemically defined media with soluble cecal content material or bile extract to investigate if nutrients available in the intestinal tract could rescue *Δkor2AEBG* mutant strain growth. When chemically defined media was supplemented with 10 % cell-free sterile rat cecum aqueous content or with 2 % ox bile, they strongly supported growth of the *Δkor2AEBG* strain compared to the parent strain (Supplemental file S10D, H). The growth of the *Δkor2AEBG* and *Δkor2AEBG ΔcitS Δicd ΔacnA* mutant strains in ox bile was restored in a dose-dependent manner compared to the parent strain (Fig. 12A-D). There was not a growth defect of the *ΔcitS Δicd ΔacnA* triple mutant strain. The addition of 0.2% taurodeoxycholic (TDCA) acid or 0.2% glycocholic acid (GCA) had no stimulatory growth effect on either *Δkor2AEBG* or *Δkor2AEBG ΔcitS Δicd ΔacnA* mutant strains compared to the parent strain (Fig. 12A-D). This suggests that bile components other than bile salts were involved in growth stimulation of the mutant strains lacking *kor2AEBG* genes.

**Fig. 12.**
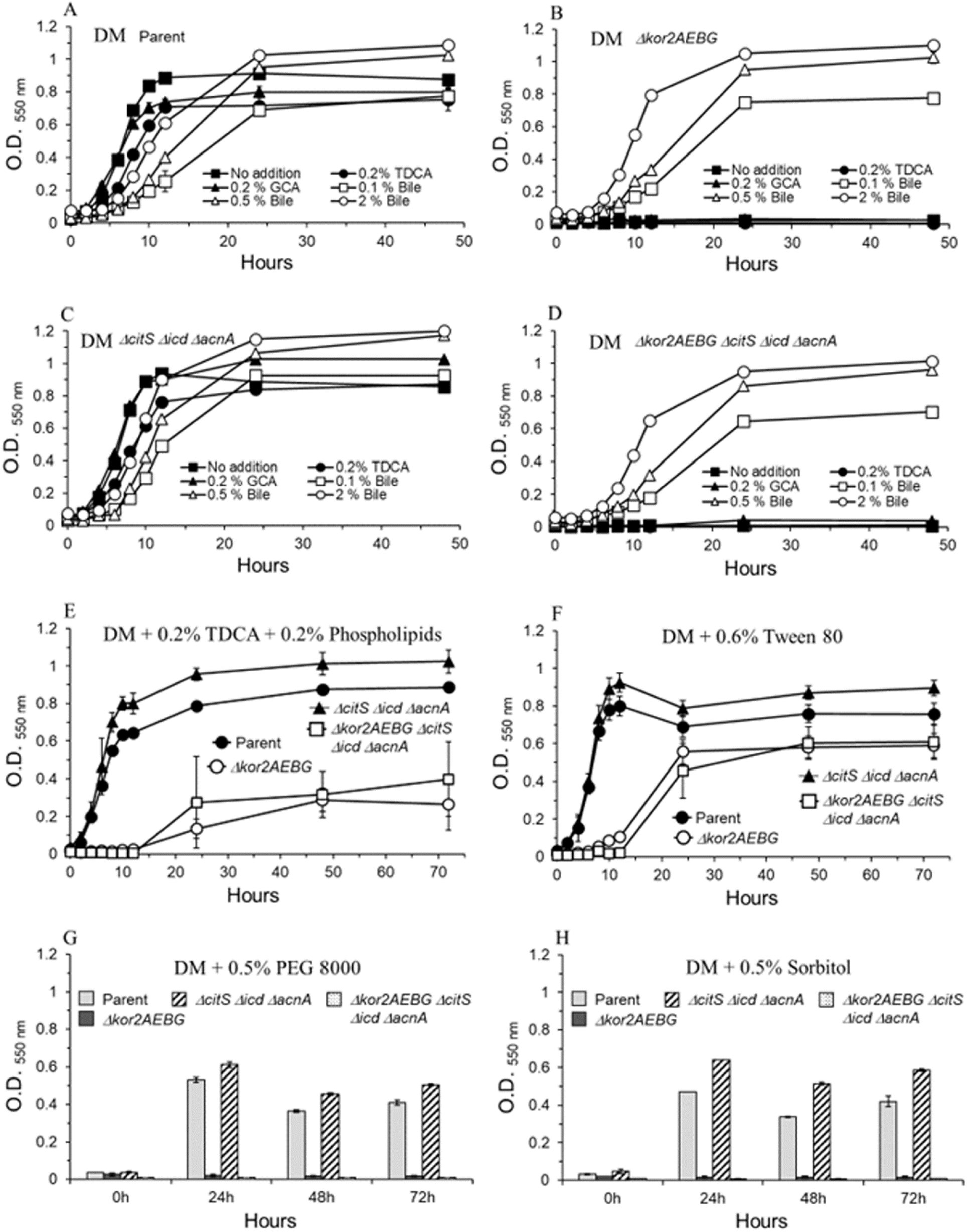
Growth of *B. fragilis* mutant strains in chemically defined media with glucose (DM). Panels A to D: DM with glucose was supplemented with 0.2% glycocholic acid (GCA), 0.2% taurodeoxycholic acid (TDCA), 0.1%, 0.5% or 2% ox-bile (Bile), or no addition. A: BF638R parent strain. B: *Δkor2AEBG* single mutant strain. C: *ΔcitS Δicd ΔacnA* triple mutant strain. D: *Δkor2AEBG ΔcitS Δicd ΔacnA* quadruple mutant strain. Panels E to H: The supplements added to DM are indicated in each panel. Phospholipids: Soy (Glycine max) phospholipids mixture, Sigma Aldrich Cat # 11145, Lot/Batch # BCCJ8356, contains roughly equal proportions of lecithin, cephalin, and phosphatidylinositol along with minor amounts of other phospholipids and polar lipids. It contains about 24% saturated fatty acids, 14% mono-unsaturated and 62% poly-unsaturated fatty acid. TDCA was mixed with phospholipids to solubilize micelles (Panel E). PEG: polyethylene glycol. Sorbitol was added instead of sorbitan (1,4-sorbitol cyclic ester derivative of sorbitol dehydration), a component of Tween 80, which is a polyethoxylated sorbitan with one oleic acid as a primary fatty acid. Strain designations are depicted in each panel.

### Addition of phospholipids or tween 80 restores growth of *Δkor2AEBG* mutant in chemically defined media

When these same mutant strains were grown in defined media containing 0.2% of a soy phospholipids mixture (in the presence of TDCA to solubilize micelles), it caused a partial growth stimulation of the *Δkor2AEBG* and *Δkor2AEBG ΔcitS Δicd ΔacnA* mutant strains (Fig. 12E) compared to no addition control (Fig. 12B,D). In addition, when the surfactant and emulsifier tween 80 was added to the media, it caused the *Δkor2AEBG* and *Δkor2AEBG ΔcitS Δicd ΔacnA* mutant strains to grow to levels comparable to the parent strain though with an extended lag-growth phase (Fig. 12F). The growth kinetics manner of the *Δkor2AEBG* and *Δkor2AEBG ΔcitS Δicd ΔacnA* mutant strains in defined media containing tween 80 was similar to growth kinetics observed in BHIS containing dM-2KG as shown above (Fig. 11B). The addition of PEG 8000 or sorbitan (components of tween 80) did not affect the growth of the *Δkor2AEBG* nor *Δkor2AEBG ΔcitS Δicd ΔacnA* mutant strain (Fig. 12G,H). Taken together, these findings indicate that the ThPP-binding Kor2CDAEBG enzyme complex has a novel metabolic function essential for *B. fragilis* growth that remains to be characterized. Collectively, our investigation reveals new information on *B. fragilis* central metabolism and its modulatory control by pirin proteins. (Fig. 13).

**Fig. 13.**
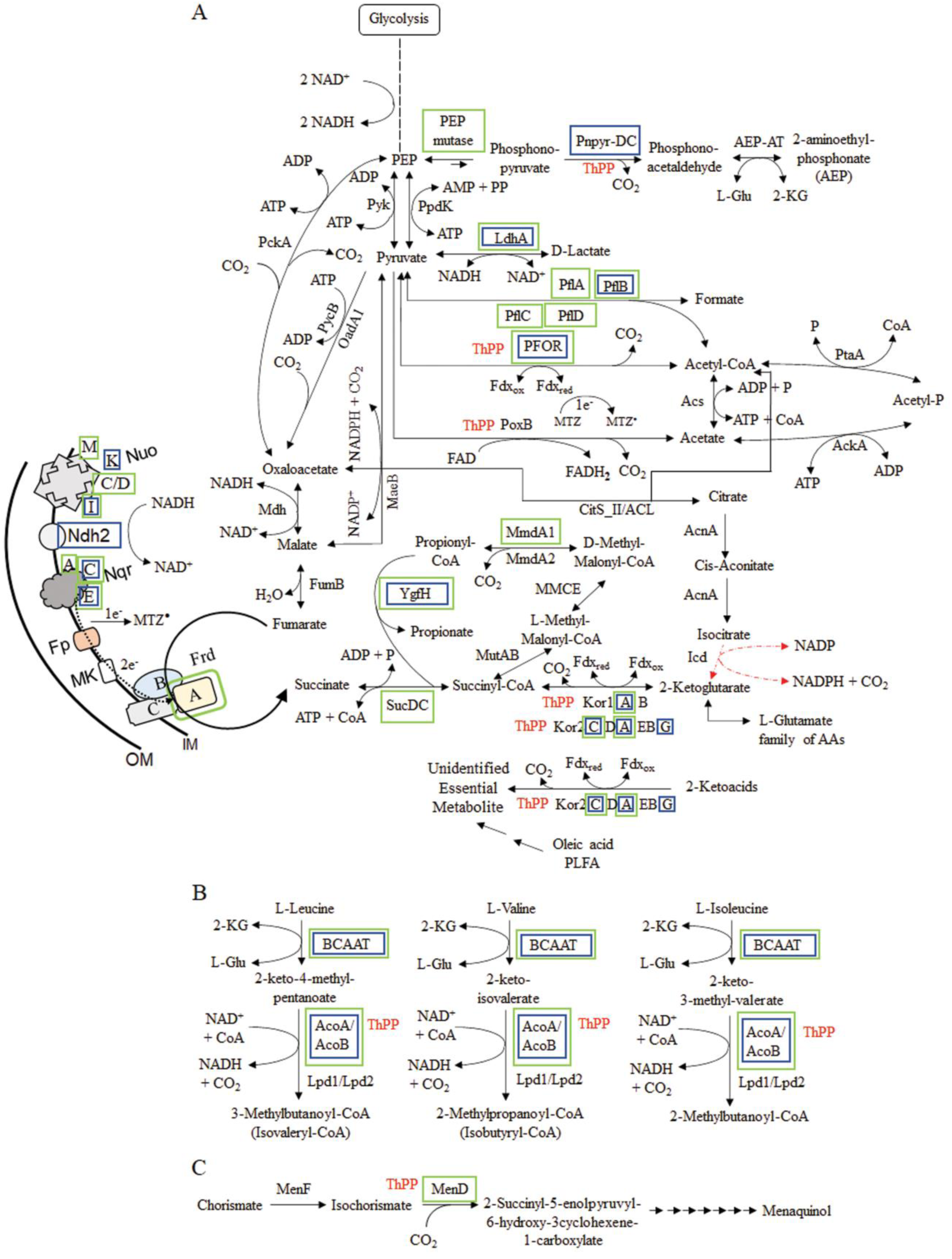
Schematic diagram of *B. fragilis* 638R (A) central metabolism, (B) degradation of branched amino acids pathway, and (C) initial steps of menaquinol biosynthetic pathway. The thiamine diphosphate binding enzymes are depicted with ThPP in red font beside enzyme designation. The enzyme subunits with predicted stable protein-protein interaction with Pir1 is highlighted with a blue box, and interaction with Pir2 with a green box. The compilation of the pathway reactions were based on public database https://www.genome.jp/pathway/bfg00020, /pathway/bfg00010, /pathway/bfg00620, /pathway/bfg00640, /pathway/bfg00280, /pathway/bfg00130), *B. fragilis* 638R genome (GenBank accession number FQ312004), and Refs.: Allison & Robinson, 1970; Allison et al., 1979; Butler et al., 2023; Harris & Reddy, 1977; Ito et al., 2020; Macy et al., 1978; Rios-Covian et al., 2015; Schofield et al., 2018; Sund et al., 2008;. Zhang et al., 2003. Orange dash/dot arrows indicate there is no report that 2-ketoglutrate or L-glutamate is formed from isocitrate dehydrogenase (Icd) pathway under anaerobic conditions (Allison & Robinson, 1970, Allison et al., 1979; Schofield et al., 2018). **AAs**: amino acids. **AckA**: Acetate kinase (BF638R_0490). **AcoA/AcoB**: 2-Ketoisovalerate dehydrogenase, single peptide containing alpha- and beta-subunit domains (BF638R_1637). **Acs**: Acetate:CoA ligase (BF638R_4449). **AcnA**: Aconitase (BF638R_3569). **AEP-AT**: 2-Aminoethylphosphonate aminotransferase (BF638R_1869) and (BF638R_3511). **BCAAT**: Branched-chain amino acid transferase (BF638R_3846). **CitS_II/ACL**: citrate synthase/2-methylcitrate synthase/ACL family (BF638R_3567). **Icd**: Isocitrate dehydrogenase (BF638R_3568). **Frd**: Fumarate reductase, FrdCAB (BF638R_4499-4501). **FumB**: Fumarate hydratase class I, anaerobic (BF638R_0646). **Kor1AB**: 2-Ketoglutarate ferredoxin oxidoreductase (BF638R_4321-4322). **Kor2CDAEBG**: 2-Ketoglutarate ferredoxin oxidoreductase (BF638R_1660-1665). **LdhA**: D-Lactate dehydrogenase (BF638R_1473). **Lpd1**: Dihydrolipoamide dehydrogenase-E3 (BF638R_0023). **Lpd2**: Dihydrolipoamide dehydrogenase-E3 (BF638R_1634). **MaeB**: Malate dehydrogenase [oxaloacetate-decarboxylating NADP^+^-dependent] (BF638R_3435). **Mdh**: Malate dehydrogenase (BF638R_0537). **MmdA1** (PccB): Propionyl-CoA carboxylase beta-chain (BF638R_1625). **MmdA2** (PccB): Propionyl-CoA carboxylase beta-chain (BF638R_3367). **MMCE**: Methylmalonyl-CoA epimerase (BF638R_3151). **MenD**: 2-Succinyl-5-enolpyruvyl-6-hydroxy-3-cyclohexene-1-carboxylic-acid synthase (BF638R_1316). **MenF**: Isochorismate synthase (BF638R_1315). **MutAB**: Methylmalonyl-CoA mutase, small (A) and large (B) subunits (BF638R_3616-3615). **Nad2**: NADH dehydrogenase, FAD-containing subunit (BF638R_1612). **Nqr**: Na+-translocating NADH-quinone oxidoreductase, NqrABCDE, (BF638R_2136_2140). **Nuo**: NADH:ubiquinone oxidoreductase, NuoABC/DHIJKLMN (BF638R_0850-0841). **OadA1**: Pyruvate/oxaloacetate carboxyltransferase (BF638R_2828). **PckA**: Phosphoenolpyruvate carboxykinase (BF638R_4326). **PEP mutase**: Phosphoenolpyruvate mutase (BF638R_1867). **PflA**: Pyruvate formate-lyase 1, activating enzyme (BF638R_1338). **PflB**: Pyruvate formate-lyase 1 [formate acetyltransferase 1] (BF638R_1339). **PflC**: Pyruvate formate-lyase 2, activating enzyme (BF638R_4262). **PflD**: Pyruvate formate-lyase 2 [formate acetyltransferase 2] 9BF638R_4263). **PFOR**: Pyruvate ferredoxin oxidoreductase (BF638R_3194). **Pnpyr-DC**: Phosphonopyruvate decarboxylase (BF638R_1868). **PoxB**: an inner membrane enzyme that catalyze oxidative decarboxylation of pyruvate to form acetate + CO_2_ (BF638R_3245). **PpdK**: Pyruvate orthophosphate dikinase (BF638R_2565). **PtaA**: Phosphate acetyltransferase (BF638R_0489). PycB: Pyruvate carboxylase biotin-containing subunit (BF638R_1927). **Pyk**: Pyruvate kinase (BF638R_4359). **SucDC**: Succinyl-CoA synthase alpha- and beta-chains (BF638R_2360-2361). **YgfH**: Succinate-CoA transferase subfamily (BF638R_0025). **PFLA**: Phospholipid fatty acids. **IM**: Inner cytoplasmic membrane. **OM**: Outer membrane. **Fp**: flavoprotein. MK: menaquinone.

## DISCUSSION

In this study, we show that Pir1 and Pir2 proteins are involved in modulating anaerobic fermentation pathways as demonstrated by changes in the SCFA products in different growth conditions and alteration of susceptibility to the antimicrobials MTZ and AMIX. These changes in metabolism correlate with protein-protein interactions of Pir1 or Pir2 with PFOR and/or Zn-ADH as demonstrated by bacterial THS assays and potentially with several other enzymes acting in the central carbon metabolism and energy preservation pathways as predicted by computational structural models (Supplemental File S5). A schematic description of the metabolic enzymes of the central metabolism and energy generating processes, degradation of branched amino acids, and synthesis of menadione that form protein-protein interaction with Pir1, Pir2 or both Pir1 and Pir2 is shown in Fig. 13. These findings agree with reports demonstrating a role for pirin in the control of TCA cycle switching to fermentation metabolism or to aerobic respiration in aerobic and facultatively anaerobic bacteria (Soo et al., 2007; Hansen et al., 2012; Tala et al., 2018; Young et al., 2023). It remains to be defined if *B. fragilis* pirins undergo Fe(II) to Fe(III) redox changes during oxygen exposure or iron-limiting conditions to modulate protein-protein interactions and conformational dynamics (Ahsan et al., 2023); Barman & Hamelberg, 2016; Liu et al., 2013). It is in the pirin active Fe(III)-form, but not in the inactive Fe(II)-form that human pirin coordinates binding to and regulates the nuclear factor NF-κB function (Liu et al., 2013). In this regard, we presume that the predicted stable protein-protein interactions of pirins with the *Bacteroides* aerotolerance protein BatC (Tang et al., 1999), and with oxygen sensitive enzymes such as PFOR, Pfl, Frd, aconitase, fumarase, and NADH dehydrogenases (Khademian & Imlay, 2020, Lu & Imlay, 2019; Lu & Imlay, 2021; Pan & Imlay, 2001) might play a role in protecting or regulating redox sensitive processes during aerotolerance when expression of *pir1* and *pir2* genes are up-regulated by oxygen (Fig. 13 and Supplemental file S5A).

Incidentally, constitutive expression of Pir1 or Pir2 significantly increased BF638R parent strain sensitivity to MTZ to similar levels of sensitivity found in the *ΔfrdB* mutant following oxygen exposure. Here we show that in a mutant lacking fumarate reductase activity, a backup/retention of NADH dehydrogenase coupled redox pair cycling electron transfer is assumed to occur. This is supported by our findings showing that inhibitors of the ETS such as acriflavine, that inhibits NADH dehydrogenase reduction of flavoprotein, and HQNO that inhibits electron flow of reduced quinone as electron donor for the reduction of fumarate to succinate by fumarate reductase (Harris & Reddy, 1997), increase MTZ sensitivity. Support for an NADH backup/retention occurrence is the fact that the level of MTZ susceptibility of the *ΔfrdB* deletion mutant strain was not further altered in the presence of ETS inhibitors compared to the parent strain (Fig. 7A and B). Moreover, whether the predicted stable protein-protein interactions of Pir2 with fumarate reductase FrdA subunit, and the interactions of Pir1 and Pir2 with subunits of all three NADH dehydrogenase types encoded in *B. fragilis* 638R genome; NADH:ubiquinone oxidoreductase (Nuo), NADH:quinone oxidoreductase (Nqr), and the NADH FAD-containing dehydrogenase II (Ndh2) (Butler et al., 2023; Ito et al., 2020) play a role in the modulation of NADH dehydrogenases redox activities remains to be defined (Fig. 13 and Supplemental file S5). Taken together, it is likely that dysregulation of the cellular membrane bioenergetics redox balance could account for an increase in reduction of MTZ.

In contrast to the effects of ETS inhibitors in increasing MTZ susceptibility, addition of the redox cycling agents 1,4-naphthoquinone (NQ) and 2-hydroxy-1,4-naphthoquinone (HNQ), which are analogues to menadione (2-methyl-1,4-naphthoquinone) to the culture media at 10 to 20 μM abolished *B. fragilis* 638R sensitivity to MTZ as determined by agar dilution and disc inhibition assays (Supplemental file S11, S12A, C). Plumbagin (5-hydroxy-2-methyl-1,4-naphthoquinone) and menadione caused lesser effect while there was no effect observed with 1,4-benzoquinone (BQ) as determined by disc inhibition assays (Supplemental file S12D, E, F). Addition of the redox cycling agents paraquat (methyl viologen) or benzyl viologen did not affect MTZ susceptibility compared to no addition control (Supplemental file S12G, H). The effect of redox cycling agents on MTZ susceptibility was not due to bacterial growth inhibition because they did not significantly affect the CFU counts in anaerobic conditions or in cultures exposed to oxygen (Supplementary file S13). Although it is unclear what redox mechanisms are affected by HNQ or NQ to cause high levels of resistance to MTZ, it appears to be linked to the electron transport mechanisms for the reduction of fumarate to succinate by fumarate reductase since the Δ*frdB* strain remained more susceptible to MTZ in the presence of HNQ than the parent strain (Supplemental file S12A, B). Moreover, the addition of HNQ did not alter *B. fragilis* 638R susceptibility to other antimicrobials such as nitrofurantoin, tetracycline, or chloramphenicol compared to no addition control (Supplemental file S12I, J). This indicates that the HNQ or NQ effect on MTZ susceptibility occurred possibly through the disruption of bioenergetics processes. Energy conservation mechanisms in commensal and pathogenic intestinal anaerobes are not well understood and among *Bacteroides* themselves there seem to have significant differences in bioenergetics which could be explored for novel antimicrobials. For example, the susceptibility to the antiparasitic and antibacterial agent closantel, a hydrogen ionophore that decouples oxidative phosphorylation and leads to inhibition of ATP synthesis (Gooyit & Janda, 2016; Hlasta et al., 1998; Rajamuthiah et al., 2014; Skuce & Fairweather, 1990; Stephenson et al., 2000; Tran et al., 2016) varied immensely when comparing *B. fragilis, B. thetaiotaomicron, and B. vulgatus* (Supplemental file S14). This also seems to be the condition among different anaerobic bacteria such as *Porphyromonas gingivalis*, *Prevotella melaninogenica*, and *Clostridioides difficile* as their susceptibility to MTZ in the presence of HNQ varies greatly compared to *B. fragilis* (Supplemental file S15).

Moreover, we show in this study that the *ΔPFOR::tetQ Δkor1AB Δkor2AEBG* triple mutant strain did not alter MTZ resistance compared to *ΔPFOR::tetQ* single mutant indicating that KFOR family members may not be the major generator of reductive power to reduce MTZ. However, it was the additional deletion of the *citS icd acnA* genes with *ΔPFOR::tetQ Δkor2ABG* background in the Δ*PFOR::tetQ Δkor2ABG ΔcitS Δicd ΔacnA* quintuple mutant strain that caused a 4-fold increase in MTZ MIC (4 μg/ml). It is not clear what role deletion of the *citS icd acnA* genes played in enhancing MTZ resistance. The *ΔcitS Δicd ΔacnA* triple mutant alone also had a 2-fold MIC increase in MTZ (1 μg/ml) and AMIX (4 μg/ml) resistance compared to the parent strain. Interestingly, no synthesis of 2-KG or L-glutamate has been demonstrated to occur via the citrate-isocitrate pathway in *Bacteroides* (Allison & Robinson 1970; Allison et al., 1979; Schofield et al., 2018) nor does it compensate for the lack of KGOR orthologs (this study). Although the regulation and functionality of the *citS icd acnA* operon in *B. fragilis* remains to be further explored, its deficiency may have altered other metabolic activities to maintain the redox carbon balance flow and energy generation. For example, MTZ resistance in *B. fragilis* is linked to high activity of lactate dehydrogenase which compensates for decreased activity of PFOR in the presence of MTZ (Narikawa et al., 1991).

Another aspect of *B. fragilis* metabolism revealed in this study was the role of KFOR orthologs Kor1AB and Kor2CDAEBG putative protein complexes. In the KEGG pathway database, the function of *kor1AB* and *kor2ABG* genes are indicated to perform reductive synthesis of 2-KG via carboxylation of succinyl-CoA (https://www.genome.jp/pathway/bfg00020 and https://www.genome.jp/pathway/bfg00620). This agrees with the fact that the *kor1AB* and *kor2ABG* gene orthologs in several microorganisms have been shown to synthesize 2-KG form succinyl-CoA (Chen et al., 2019; Dörner & Boll, 2002; Mai & Adams 1996; Yamamoto et al., 2006; Yamamoto et al., 2010). However, our findings indicate that Kor2AEBG may have an uncharacterized metabolic activity other than reductive synthesis of 2-KG since addition of L-glutamate, L-glutamine, or tryptone did not restore growth of the *Δkor2AEBG* mutant in chemically defined media (Supplemental file S10E,F,I,J,K).

In this regard, we showed in this study that rat cecal content or ox bile strongly restored growth deficiency of the *Δkor2AEBG* deletion strain in defined media (Supplemental file 10D,H). We ruled out that bile salts were involved (Fig. 12B,D) and showed that addition of a soy phospholipids mixture (mostly lecithin, cephalin and phosphatidylinositol, see Fig. 12 legend) or oleic acid component of tween 80 are sufficient for restoring growth of the *Δkor2AEBG* deletion strain. It remains unclear which metabolic pathway is used by *B. fragilis* to stimulate growth in the presence of oleic acid in the absence of *kor2ADBG* genes, however; we presume that oleic acid or other fatty acids might be a contributing factor present in bile for growth stimulation of bile-resistant anaerobic bacteria (Holdeman et al., 1977). We do not think that production of acetyl-CoA from fatty acid β-oxidation degradation is involved in compensating for the lack of Kor2AEBG since other pathways for acetyl-CoA production from decarboxylation of pyruvate such as PFOR or Pfl remain intact. Studies have shown that degradation of fatty acids in microorganisms occur via coordinated alternations between α-oxidation, β-oxidation, ω-oxidation [(ω1), (ω2), (ω3), (ω4)-type oxidation] and fatty acid hydroxylation by fatty acid hydratases mechanisms which can produce shorter dicarboxylic acids, shorter hydroxy-fatty acids, keto-fatty acids, and short branched-dicarboxylic acids (from long branched-fatty acids) (Child et al., 2019; Gatter et al., 2014; Miura & Fulco, 1974; Hagedoom et al., 2021; Kang et al., 2017a; Kang et al., 2017b; Miura & Fulco, 1975; Miura, 2013; Ruettinger et al., 1974; Shoun et al., 1985; Vanhanen et al., 2000). However, the product(s) of 2-ketoacid oxidoreductase metabolism from Kor2AEBG activities that intersects with oleic acid or other long fatty acids catabolism remains to be defined (Fig. 13).

In conclusion, we show that the central metabolism and bioenergetics of *B. fragilis* still have many features that are yet to be explored and are potential targets for the developments of novel narrow spectrum antibiotics. One of these targets is the ThPP-binding enzymes related to anaerobic bacteria metabolism to which AMIX was shown to be effective against *B. fragilis in vivo*. Although intraperitoneal administration of AMIX significantly decreased CFU counts, compared to elimination of *B. fragilis* with intra-cage administration, this difference may have been due to diminished systemic antimicrobial diffusion through the encapsulated barrier of the artificial tissue cage. Future studies using different experimental abscess models would clarify this matter. Nonetheless, our findings agree with previous studies showing the effectiveness of AMIX against infection caused by *C. difficile*, *H. pylori*, *C. jejuni*, and by oral anaerobic pathogens in infections such as gingivitis, periodontitis, and in biofilms in *in vivo* animal models (Gui et al., 2019; Gui et al., 2020; Hoffman et al., 2014; Hoffman, 2020; Hutcherson et al., 2017; Kennedy et al., 2016; Reed et al., 2018; Warren et al., 2012). Taken together, these findings show that AMIX is a potential antimicrobial for *B. fragilis* extra-intestinal infection. Lastly, we show evidence that the α*-*ketoglutarate ferredoxin oxidoreductase Kor2CDADBG is an essential enzyme for *B. fragilis* growth and plays a novel function in anaerobic central metabolism which remains to be completely characterized. We believe that understanding the *B. fragilis* central carbon metabolism and energy-conservation pathways may lead to the development of novel narrow-spectrum selective antimicrobials for the inhibition of essential metabolic targets (Baek et al., 2011; Bunik et al., 2013; Cook et al., 2014; Feng et al. 2015; Gil-Gil et al. 2020; Murina et al., 2014; Stokes et al., 2019).

## Data availability

The data that supports the findings of this work are available at https://www.ncbi.nlm.nih.gov/gds/. GEO Datasets GSE241210 and GSE241676.

## Ethics statement

All procedures involving animals followed the guidelines given by the National Research Council’s *Guide for the Care and Use of Laboratory Animals* (National Research Council, 2011) and approved by the Institutional Animal Care and Use Committee of East Carolina University.

## ACKNOWLEDGMENTS

This work was carried out with the support from the funds available from the Dept. of Microbiology and Immunology at Brody School of Medicine to which ERR is very thankful.

## REFERENCES

Adamsson I, Nord CE, Lundquist P, Sjöstedt S, Edlund C. 1999. Comparative effects of omeprazole, amoxycillin plus metronidazole versus omeprazole, clarithromycin plus metronidazole on the oral, gastric and intestinal microflora in *Helicobacter pylori*-infected patients. J Antimicrob Chemother. 44(5):629–640. doi: 10.1093/jac/44.5.629. PMID: 10552979.

Ahsan T, Shoily SS, Ahmed T, Sajib AA. 2023. Role of the redox state of the Pirin-bound cofactor on interaction with the master regulators of inflammation and other pathways. PLoS One. 18:e0289158. doi: 10.1371/journal.pone.0289158. PMID: 38033031.

Alauzet C, Lozniewski A, Marchandin H. 2019. Metronidazole resistance and *nim* genes in anaerobes: A review. Anaerobe. 55:40–53. doi: 10.1016/j.anaerobe.2018.10.004. PMID: 30316817.

Allison MJ, Robinson IM. 1970. Biosynthesis of alpha-ketoglutarate by the reductive carboxylation of succinate in *Bacteroides ruminicola*. J Bacteriol. 104:50–56. doi: 10.1128/jb.104.1.50-56.1970. PMID: 5473908.

Allison MJ, Robinson IM, Baetz AL. 1979. Synthesis of alpha-ketoglutarate by reductive carboxylation of succinate in *Veillonella, Selenomonas*, and *Bacteriodes* species. J Bacteriol. 140(3):980–6. doi: 10.1128/jb.140.3.980-986.1979. PMID: 533772.

An J, Sun JY, Yuan Q, Tian HY, Qiu WL, Guo W, Zhao FK. 2004. Proteomics analysis of differentially expressed metastasis-associated proteins in adenoid cystic carcinoma cell lines of human salivary gland. Oral Oncol. 40:400–408. doi: 10.1016/j.oraloncology.2003.09.014. PMID: 14969819.

Agarwal G, Rajavel M, Gopal B, Srinivasan N. 2009. Structure-based phylogeny as a diagnostic for functional characterization of proteins with a cupin fold. PLoS One 4:e5736. doi: 10.1371/journal.pone.0005736. PMID: 19478949.

Bäckhed F, Fraser CM, Ringel Y, Sanders ME, Sartor RB, Sherman PM, Versalovic J, Young V, Finlay BB. 2012. Defining a healthy human gut microbiome: current concepts, future directions, and clinical applications. Cell Host Microbe. 12:611–622. doi: 10.1016/j.chom.2012.10.012. PMID: 23159051.

Baek SH, Li AH, Sassetti CM. 2011. Metabolic regulation of mycobacterial growth and antibiotic sensitivity. PLoS Biol. 9:e1001065. doi: 10.1371/journal.pbio.1001065. Epub 2011 May 24. PMID: 21629732.

Ballard TE, Wang X, Olekhnovich I, Koerner T, Seymour C, Salamoun J, Warthan M, Hoffman PS, Macdonald TL. 2011. Synthesis and antimicrobial evaluation of nitazoxanide-based analogues: identification of selective and broad spectrum activity. Chem Med Chem. 6:362–367. PMID: 21275058.

Barman A, Hamelberg D. 2016. Fe(II)/Fe(III) Redox Process Can Significantly Modulate the Conformational Dynamics and Electrostatics of Pirin in NF-κB Regulation. ACS Omega. 1:837–842. doi: 10.1021/acsomega.6b00231. PMID: 31457166.

Baughn AD, Malamy MH. 2002. A mitochondrial-like aconitase in the bacterium *Bacteroides fragilis*: implications for the evolution of the mitochondrial Krebs cycle. Proc Natl Acad Sci U S A. 99:4662–4667. doi: 10.1073/pnas.052710199. PMID: 11880608.

Baughn AD, Malamy MH. 2003. The essential role of fumarate reductase in haem-dependent growth stimulation of *Bacteroides fragilis*. Microbiology (Reading). 149(:1551–1558. doi: 10.1099/mic.0.26247-0. PMID: 12777495.

Betteken MI, Rocha ER, Smith CJ. 2015. Dps and DpsL mediate survival *in vitro* and *in vivo* during the prolonged oxidative stress response in *Bacteroides fragilis*. J Bacteriol 197:3329–3338. doi:10.1128/JB.00342-15.

Bock AK, Kunow J, Glasemacher J, Schönheit P. 1996. Catalytic properties, molecular composition and sequence alignments of pyruvate: ferredoxin oxidoreductase from the methanogenic archaeon *Methanosarcina barkeri* (strain Fusaro). Eur J Biochem. 237:35–44. PMID: 8620891.

Buckel W, Thauer RK. 2018. Flavin-Based Electron Bifurcation, Ferredoxin, Flavodoxin, and Anaerobic Respiration With Protons (Ech) or NAD+ (Rnf) as Electron Acceptors: A Historical Review. Front Microbiol. 9:401. doi: 10.3389/fmicb.2018.00401. PMID: 29593673.

Buckel W, Thauer RK. 2018. Flavin-Based Electron Bifurcation, A New Mechanism of Biological Energy Coupling. Chem Rev. 118:3862–3886. doi: 10.1021/acs.chemrev.7b00707. PMID: 29561602.

Bunik VI, Tylicki A, Lukashev NV. 2013. Thiamin diphosphate-dependent enzymes: from enzymology to metabolic regulation, drug design and disease models. FEBS J. 280:6412–6442. doi: 10.1111/febs.12512. PMID: 24004353.

Butler NL, Ito T, Foreman S, Morgan JE, Zagorevsky D, Malamy MH, Comstock LE, Barquera B. 2023. *Bacteroides fragilis* Maintains Concurrent Capability for Anaerobic and Nanaerobic Respiration. J Bacteriol. 205:e0038922. doi: 10.1128/jb.00389-22. PMID: 36475831.

Byun JH, Kim M, Lee Y, Lee K, Chong Y. 2019. Antimicrobial Susceptibility Patterns of Anaerobic Bacterial Clinical Isolates From 2014 to 2016, Including Recently Named or Renamed Species. Ann Lab Med. 39:190–199. doi: 10.3343/alm.2019.39.2.190. PMID: 30430782.

Chen M, Wolin MJ. 1981. Influence of heme and vitamin B_12_ on growth and fermentations of *Bacteroides* species. J Bacteriol. 145:466–471. doi: 10.1128/jb.145.1.466-471.1981. PMID: 7462148.

Chen PY, Li B, Drennan CL, Elliott SJ. 2019. A reverse TCA cycle 2-oxoacid:ferredoxin oxidoreductase that makes C-C bonds from CO_2_. Joule. 3:595–611. doi: 10.1016/j.joule.2018.12.006. PMID: 31080943.

Chen L, Chen X, Bai Y, Zhao ZN, Cao YF, Liu LK, Jiang T, Hou J. 2022. Inhibition of *Escherichia coli* nitroreductase by the constituents in *Syzygium aromaticum*. Chin J Nat Med. 20:506–517. doi: 10.1016/S1875-5364(22)60163-8. PMID: 35907649.

Child SA, Rossi VP, Bell SG. 2019. Selective ω-1 oxidation of fatty acids by CYP147G1 from *Mycobacterium marinum*. Biochim Biophys Acta Gen Subj. 1863:408–417. doi: 10.1016/j.bbagen.2018.11.013. PMID: 30476524.

Cook GM, Greening C, Hards K, Berney M. 2014. Energetics of pathogenic bacteria and opportunities for drug development. Adv Microb Physiol. 65:1–62. doi: 10.1016/bs.ampbs.2014.08.001. PMID: 25476763.

Delday M, Mulder I, Logan ET, Grant G. 2019. *Bacteroides thetaiotaomicron* Ameliorates Colon Inflammation in Preclinical Models of Crohn’s Disease. Inflamm Bowel Dis. 25:85–96. doi: 10.1093/ibd/izy281. PMID: 30215718.

Diniz CG, Farias LM, Carvalho MA, Rocha ER, Smith CJ. 2004. Differential gene expression in a *Bacteroides fragilis* metronidazole-resistant mutant. J Antimicrob Chemother. 54:100–108. doi: 10.1093/jac/dkh256. PMID: 15150173.

Dingsdag SA, Hunter N. 2018. Metronidazole: an update on metabolism, structure-cytotoxicity and resistance mechanisms. J Antimicrob Chemother. 73:265–279. doi: 10.1093/jac/dkx351. PMID: 29077920.

Dörner E, Boll M. 2002. Properties of 2-oxoglutarate:ferredoxin oxidoreductase from *Thauera aromatica* and its role in enzymatic reduction of the aromatic ring. J Bacteriol. 184:3975–3983. doi: 10.1128/JB.184.14.3975-3983.2002. PMID: 12081970.

Doucette CD, Schwab DJ, Wingreen NS, Rabinowitz JD. 2011. α-Ketoglutarate coordinates carbon and nitrogen utilization via enzyme I inhibition. Nat Chem Biol. 7:894–901. doi: 10.1038/nchembio.685. PMID: 22002719.

Dunwell JM, Khuri S, Gane PJ. 2000. Microbial relatives of the seed storage proteins of higher plants: conservation of structure and diversification of function during evolution of the cupin superfamily. Microbiol Mol Biol Rev. 64:153–179. doi: 10.1128/MMBR.64.1.153-179.2000. PMID: 10704478.

Dunwell JM, Culham A, Carter CE, Sosa-Aguirre CR, Goodenough PW. 2001. Evolution of functional diversity in the cupin superfamily. Trends Biochem Sci. 26:740–746. doi: 10.1016/s0968-0004(01)01981-8. PMID: 11738598.

Dunwell JM, Purvis A, Khuri S. 2004. Cupins: the most functionally diverse protein superfamily? Phytochemistry. 65(1):7–17. doi: 10.1016/j.phytochem.2003.08.016. PMID: 14697267.

Evans R, O’Neill, Pritzel A. et al., 2021. Protein complex prediction with AlphaFold-Multimer. bioRxiv 2021.10.04.463034. 10.1101/2021.10.04.463034.

Farr SB, Kogoma T. 1991. Oxidative stress responses in *Escherichia coli* and *Salmonella typhimurium*. Microbiol Rev. 55:561–585. doi: 10.1128/mr.55.4.561-585.1991. PMID: 1779927.

Feng X, Zhu W, Schurig-Briccio LA, Lindert S, Shoen C, Hitchings R, Li J, Wang Y, Baig N, Zhou T, Kim BK, Crick DC, Cynamon M, McCammon JA, Gennis RB, Oldfield E. 2015. Antiinfectives targeting enzymes and the proton motive force. Proc Natl Acad Sci U S A. 112:E7073–82. doi: 10.1073/pnas.1521988112. PMID: 26644565.

Gatter M, Förster A, Bär K, Winter M, Otto C, Petzsch P, Ježková M, Bahr K, Pfeiffer M, Matthäus F, Barth G. 2014. A newly identified fatty alcohol oxidase gene is mainly responsible for the oxidation of long-chain ω-hydroxy fatty acids in *Yarrowia lipolytica*. FEMS Yeast Res. 14:858–872. doi: 10.1111/1567-1364.12176. PMID: 24931727.

Gauss GH, Reott MA, Rocha ER, Young MJ, Douglas T, Smith CJ, Lawrence CM. 2012. Characterization of the *Bacteroides fragilis bfr* gene product identifies a bacterial DPS-like protein and suggests evolutionary links in the ferritin superfamily. J Bacteriol. 194:15–27. doi: 10.1128/JB.05260-11.. PMID: 22020642.

Ghotaslou R, Bannazadeh Baghi H, Alizadeh N, Yekani M, Arbabi S, Memar MY. 2018. Mechanisms of *Bacteroides fragilis* resistance to metronidazole. Infect Genet Evol. 64:156–163. doi: 10.1016/j.meegid.2018.06.020. PMID: 29936037.

Gibson MI, Chen PY, Drennan CL. 2016. A structural phylogeny for understanding 2-oxoacid oxidoreductase function. Curr Opin Struct Biol. 41:54–61. doi: 10.1016/j.sbi.2016.05.011. PMID: 27315560.

Gil-Gil T, Corona F, Martínez JL, Bernardini A. 2020. The Inactivation of Enzymes Belonging to the Central Carbon Metabolism Is a Novel Mechanism of Developing Antibiotic Resistance. mSystems. 5(3):e00282–20. doi: 10.1128/mSystems.00282-20. PMID: 32487742.

Gooyit M, Janda KD. 2016. Reprofiled anthelmintics abate hypervirulent stationary-phase *Clostridium difficile*. Sci Rep. 6:33642. doi: 10.1038/srep33642. PMID: 27633064.

Gui Q, Hoffman PS, Lewis JP. 2019. Amixicile targets anaerobic bacteria within the oral microbiome. J Oral Biosci. 61:226–235. doi: 10.1016/j.job.2019.10.004. Erratum in: J Oral Biosci. 2020 Sep;62(3):297. PMID: 31706024.

Gui Q, Ramsey KW, Hoffman PS, Lewis JP. 2020. Amixicile depletes the ex vivo periodontal microbiome of anaerobic bacteria. J Oral Biosci. 62:195–204. doi: 10.1016/j.job.2020.03.004. PMID: 32278683.

Gui Q, Lyons DJ, Deeb JG, Belvin BR, Hoffman PS, Lewis JP.2021. Non-human Primate Macaca mulatta as an Animal Model for Testing Efficacy of Amixicile as a Targeted Anti-periodontitis Therapy. Front Oral Health. 2:752929. doi: 10.3389/froh.2021.752929. PMID: 35048063.

Guiney DG, Hasegawa P, Davis CE. 1984. Plasmid transfer from *Escherichia coli* to *Bacteroides fragilis*: differential expression of antibiotic resistance phenotypes. Proc Natl Acad Sci U S A. 81:7203–7206. doi: 10.1073/pnas.81.22.7203. PMID: 6095273.

Hagedoorn PL, Hollmann F, Hanefeld U. 2021. Novel oleate hydratases and potential biotechnological applications. Appl Microbiol Biotechnol. 105:6159–6172. doi: 10.1007/s00253-021-11465-x. PMID: 34350478.

Haggoud A, Reysset G, Azeddoug H, Sebald M. 1994. Nucleotide sequence analysis of two 5-nitroimidazole resistance determinants from *Bacteroides strains* and of a new insertion sequence upstream of the two genes. Antimicrob Agents Chemother. 38:1047–51. doi: 10.1128/AAC.38.5.1047. PMID: 8067736.

Hansen GA, Ahmad R, Hjerde E, Fenton CG, Willassen NP, Haugen P. 2012. Expression profiling reveals Spot 42 small RNA as a key regulator in the central metabolism of *Aliivibrio salmonicida*. BMC Genomics.13:37. doi: 10.1186/1471-2164-13-37. PMID: 22272603.

Harris MA, Reddy CA. 1977. Hydrogenase activity and the H_2_-fumarate electron transport system in *Bacteroides fragilis*. J Bacteriol. 131:922–928. doi: 10.1128/jb.131.3.922-928.1977. PMID: 893348.

Hartmeyer GN, Sóki J, Nagy E, Justesen US. 2012. Multidrug-resistant *Bacteroides fragilis* group on the rise in Europe? J Med Microbiol. 61(Pt 12):1784–1788. doi: 10.1099/jmm.0.049825-0. PMID: 22956754.

Hihara Y, Muramatsu M, Nakamura K, Sonoike K. 2004. A cyanobacterial gene encoding an ortholog of Pirin is induced under stress conditions. FEBS Lett. 574:101–105. doi: 10.1016/j.febslet.2004.06.102. PMID: 15358547.

Hlasta DJ, Demers JP, Foleno BD, Fraga-Spano SA, Guan J, Hilliard JJ, Macielag MJ, Ohemeng KA, Sheppard CM, Sui Z, Webb GC, Weidner-Wells MA, Werblood H, Barrett JF. 1998. Novel inhibitors of bacterial two-component systems with gram positive antibacterial activity: pharmacophore identification based on the screening hit closantel. Bioorg Med Chem Lett. 8(14):1923–1928. doi: 10.1016/s0960-894x(98)00326-6. PMID: 9873460.

Hoffman PS, Goodwin A, Johnsen J, Magee K, Veldhuyzen van Zanten SJ. 1996. Metabolic activities of metronidazole-sensitive and -resistant strains of *Helicobacter pylori*: repression of pyruvate oxidoreductase and expression of isocitrate lyase activity correlate with resistance. J Bacteriol. 178:4822–4829. doi: 10.1128/jb.178.16.4822-4829.1996. PMID: 8759844.

Hoffman PS, Sisson G, Croxen MA, Welch K, Harman WD, Cremades N, Morash MG. 2007. Antiparasitic drug nitazoxanide inhibits the pyruvate oxidoreductases of *Helicobacter pylori*, selected anaerobic bacteria and parasites, and *Campylobacter jejuni*. Antimicrob Agents Chemother. 51:868–876. PMID: 17158936.

Hoffman PS, Bruce AM, Olekhnovich I, Warren CA, Burgess SL, Hontecillas R, Viladomiu M, Bassaganya-Riera J, Guerrant RL, Macdonald TL. 2014. Preclinical studies of amixicile, a systemic therapeutic developed for treatment of *Clostridium difficile* infections that also shows efficacy against *Helicobacter pylori*. Antimicrob Agents Chemother. 58:4703–4712. doi: 10.1128/AAC.03112-14. PMID: 24890599.

Hoffman PS. 2020. Amixicile: A Concept Therapeutic for Treatment of Chronic Anaerobic Infections. Br J Gastroenterol. 2:138–142. doi: 10.31488/bjg.1000108. PMID: 37346897.

Holdeman LV, Cato EP, Moore, WEC. 1977. Anaerobe Laboratory Manual. 4^th^ ed. Anaerobe Laboratory Virginia Polytechnic Institute and University, Blacksburg, Virginia.

Hromada S, Venturelli OS. 2023. Gut microbiota interspecies interactions shape the response of *Clostridioides difficile* to clinically relevant antibiotics. PLoS Biol. 21:e3002100. doi: 10.1371/journal.pbio.3002100. PMID: 37167201.

Huang C, Feng S, Huo F, Liu H. 2022. Effects of Four Antibiotics on the Diversity of the Intestinal Microbiota. Microbiol Spectr. 10:e0190421. doi: 10.1128/spectrum.01904-21. PMID: 35311555.

Hutcherson JA, Sinclair KM, Belvin BR, Gui Q, Hoffman PS, Lewis JP. 2017. Amixicile, a novel strategy for targeting oral anaerobic pathogens. Sci Rep. 7:10474. PMID: 28874750.

Ito T, Gallegos R, Matano LM, Butler NL, Hantman N, Kaili M, Coyne MJ, Comstock LE, Malamy MH, Barquera B. 2020. Genetic and Biochemical Analysis of Anaerobic Respiration in *Bacteroides fragilis* and Its Importance *In Vivo*. mBio. 11:e03238–19. doi: 10.1128/mBio.03238-19. PMID: 32019804.

Jain E, Zaenker EI, Hoffman PS, Warren CA. 2022. *In vitro* activity of amixicile against *T. vaginalis* from clinical isolates. Parasitol Res. 121:2453–2455. doi: 10.1007/s00436-022-07567-8. Epub 2022 Jun 9. PMID: 35676563.

Jakobsson HE, Jernberg C, Andersson AF, Sjölund-Karlsson M, Jansson JK, Engstrand L. 2010. Short-term antibiotic treatment has differing long-term impacts on the human throat and gut microbiome. PLoS One. 5:e9836. doi: 10.1371/journal.pone.0009836. PMID: 20352091.

Jasemi S, Emaneini M, Ahmadinejad Z, Fazeli MS, Sechi LA, Sadeghpour Heravi F, Feizabadi MM. 2021. Antibiotic resistance pattern of *Bacteroides fragilis* isolated from clinical and colorectal specimens. Ann Clin Microbiol Antimicrob. 20:27. doi: 10.1186/s12941-021-00435-w. PMID: 33892721.

Jumper J, Evans R, Pritzel A, et al., 2021. Highly accurate protein structure prediction with AlphaFold. Nature 596:583–589. doi: 10.1038/s41586-021-03819-2. PMID: 34265844.

Kang WR, Seo MJ, Shin KC, Park JB, Oh DK. 2017a. Comparison of Biochemical Properties of the Original and Newly Identified Oleate Hydratases from *Stenotrophomonas maltophilia*. Appl Environ Microbiol. 83:e03351–16. doi: 10.1128/AEM.03351-16. PMID: 28235876.

Kang WR, Seo MJ, Shin KC, Park JB, Oh DK. 2017b. Gene cloning of an efficiency oleate hydratase from *Stenotrophomonas nitritireducens* for polyunsaturated fatty acids and its application in the conversion of plant oils to 10-hydroxy fatty acids. Biotechnol Bioeng. 114(1):74–82. doi: 10.1002/bit.26058. PMID: 27474883.

Kennedy AJ, Bruce AM, Gineste C, Ballard TE, Olekhnovich IN, Macdonald TL, Hoffman PS. 2016. Synthesis and Antimicrobial Evaluation of Amixicile-Based Inhibitors of the Pyruvate-Ferredoxin Oxidoreductases of Anaerobic Bacteria and Epsilonproteobacteria. Antimicrob Agents Chemother. 60:3980–39877. doi: 10.1128/AAC.00670-16. PMID: 27090174.

Kennedy AJ, Bruce AM, Gineste C, Ballard TE, Olekhnovich IN, Macdonald TL, Hoffman PS. 2016. Synthesis and Antimicrobial Evaluation of Amixicile-Based Inhibitors of the Pyruvate-Ferredoxin Oxidoreductases of Anaerobic Bacteria and Epsilonproteobacteria. Antimicrob Agents Chemother. 60:3980–3987. doi: 10.1128/AAC.00670-16. PMID: 27090174.

Khademian M, Imlay JA.2020. Do reactive oxygen species or does oxygen itself confer obligate anaerobiosis? The case of *Bacteroides thetaiotaomicron*. Mol Microbiol. 114:333–347. doi: 10.1111/mmi.14516. PMID: 32301184.

Koropatkin NM, Martens EC, Gordon JI, Smith TJ. 2008. Starch catabolism by a prominent human gut symbiont is directed by the recognition of amylose helices. Structure. 16:1105–1015. doi: 10.1016/j.str.2008.03.017. PMID: 18611383.

Krissinel E, Henrick K. 2007. Inference of macromolecular assemblies from crystalline state. J Mol Biol. 372:774–797. doi: 10.1016/j.jmb.2007.05.022. PMID: 17681537.

Lapik YR, Kaufman LS. 2003. The Arabidopsis cupin domain protein AtPirin1 interacts with the G protein alpha-subunit GPA1 and regulates seed germination and early seedling development. Plant Cell. 15:1578–7590. doi: 10.1105/tpc.011890. PMID: 12837948.

Liu F, Rehmani I, Esaki S, Fu R, Chen L, de Serrano V, Liu A. 2013. Pirin is an iron-dependent redox regulator of NF-κB. Proc Natl Acad Sci U S A. 110:9722–9727. doi: 10.1073/pnas.1221743110. PMID: 23716661.

Lobo LA, Jenkins AL, Jeffrey Smith C, Rocha ER. 2013. Expression of *Bacteroides fragilis* hemolysins *in vivo* and role of HlyBA in an intra-abdominal infection model. Microbiologyopen. 2:326–337. doi: 10.1002/mbo3.76. PMID: 23441096.

Lu Z, Imlay JA. 2019. A conserved motif liganding the [4Fe-4S] cluster in [4Fe-4S] fumarases prevents irreversible inactivation of the enzyme during hydrogen peroxide stress. Redox Biol. 26:101296. doi: 10.1016/j.redox.2019.101296. PMID: 31465957.

Lu Z, Imlay JA. 2021. When anaerobes encounter oxygen: mechanisms of oxygen toxicity, tolerance and defence. Nat Rev Microbiol. 19:774–785. doi: 10.1038/s41579-021-00583-y. PMID: 34183820.

Macy J, Probst I, Gottschalk G. 1975. Evidence for cytochrome involvement in fumarate reduction and adenosine 5’-triphosphate synthesis by *Bacteroides fragilis* grown in the presence of hemin. J Bacteriol. 123:436–442. doi: 10.1128/jb.123.2.436-442.1975. PMID: 1150622.

Macy JM, Ljungdahl LG, Gottschalk G. 1978. Pathway of succinate and propionate formation in *Bacteroides fragilis*. J Bacteriol. 134:84–91. doi: 10.1128/jb.134.1.84-91.1978. PMID: 148460.

Mai X, Adams MW. 1996. Characterization of a fourth type of 2-keto acid-oxidizing enzyme from a hyperthermophilic archaeon: 2-ketoglutarate ferredoxin oxidoreductase from *Thermococcus litoralis*. J Bacteriol. 178:5890–5896. doi: 10.1128/jb.178.20.5890-5896.1996. PMID: 8830683.

Martens EC, Chiang HC, Gordon JI. 2008. Mucosal glycan foraging enhances fitness and transmission of a saccharolytic human gut bacterial symbiont. Cell Host Microbe 4:447–457. doi: 10.1016/j.chom.2008.09.007. PMID: 18996345.

Mirdita M, Schütze K, Moriwaki Y, Heo L, Ovchinnikov S, Steinegger M. 2022. ColabFold: making protein folding accessible to all. Nat Methods. 19:679–682. doi: 10.1038/s41592-022-01488-1. PMID: 35637307.

Miura Y, Fulco AJ. 1974. (ω-2) hydroxylation of fatty acids by a soluble system from *Bacillus megaterium*. J Biol Chem. 249:1880–1888. PMID: 4150419.

Miura Y, Fulco AJ. 1975. ω-1, ω-2 and ω-3 hydroxylation of long-chain fatty acids, amides and alcohols by a soluble enzyme system from *Bacillus megaterium*. Biochim Biophys Acta. 388:305–317. doi: 10.1016/0005-2760(75)90089-2. PMID: 805599.

Miura Y. 2013. The biological significance of ω-oxidation of fatty acids. Proc Jpn Acad Ser B Phys Biol Sci. 89:370–382. doi: 10.2183/pjab.89.370. PMID: 24126285.

Murima P, McKinney JD, Pethe K. 2014. Targeting bacterial central metabolism for drug development. Chem Biol. 21:1423–1432. doi: 10.1016/j.chembiol.2014.08.020. Epub 2014 Oct 16. PMID: 25442374.

Nagy E, Urbán E, Nord CE. 2011. ESCMID Study Group on Antimicrobial Resistance in Anaerobic Bacteria. Antimicrobial susceptibility of *Bacteroides fragilis* group isolates in Europe: 20 years of experience. Clin Microbiol Infect. 17:371–379. doi: 10.1111/j.1469-0691.2010.03256.x. PMID: 20456453.

Narikawa S, Suzuki T, Yamamoto M, Nakamura M. 1991. Lactate dehydrogenase activity as a cause of metronidazole resistance in *Bacteroides fragilis* NCTC 11295. J Antimicrob Chemother. 28:47–53. doi: 10.1093/jac/28.1.47. PMID: 1769942.

National Research Council. 2011. Guide for the care and use of laboratory animals, 8th ed. National Academies Press, Washington, DC.

Orzaez D, de Jong AJ, Woltering EJ. 2001. A tomato homologue of the human protein PIRIN is induced during programmed cell death. Plant Mol Biol. 46:459–468. doi: 10.1023/a:1010618515051. PMID: 11485202.

Pan N, Imlay JA. 2001. How does oxygen inhibit central metabolism in the obligate anaerobe *Bacteroides thetaiotaomicron*. Mol Microbiol. 39:1562–1671. doi: 10.1046/j.1365-2958.2001.02343.x. PMID: 11260473.

Pang H, Bartlam M, Zeng Q, Miyatake H, Hisano T, Miki K, Wong LL, Gao GF, Rao Z. 2004. Crystal structure of human pirin: an iron-binding nuclear protein and transcription cofactor. J Biol Chem. 279:1491–1498. doi: 10.1074/jbc.M310022200. PMID: 14573596.

Parker AC, Seals NL, Baccanale CL, Rocha ER. 2022. Analysis of Six tonB Gene Homologs in *Bacteroides fragilis* Revealed That *tonB3* is Essential for Survival in Experimental Intestinal Colonization and Intra-Abdominal Infection. Infect Immun. 90:e0046921. doi: 10.1128/IAI.00469-21. PMID: 34662212.

Paunkov A, Hummel K, Strasser D, Sóki J, Leitsch D. 2023. Proteomic analysis of metronidazole resistance in the human facultative pathogen *Bacteroides fragilis*. Front Microbiol. 14:1158086. doi: 10.3389/fmicb.2023.1158086. PMID: 37065137.

Pfaffl MW. 2001. A new mathematical model for relative quantification in real-time RT-PCR. Nucleic Acids Res. 29:e45. doi: 10.1093/nar/29.9.e45. PMID: 11328886.

Prajapati S, Rabe von Pappenheim F, Tittmann K. 2022. Frontiers in the enzymology of thiamin diphosphate-dependent enzymes. Curr Opin Struct Biol. 76:102441. doi: 10.1016/j.sbi.2022.102441. PMID: 35988322.

Privitera G, Dublanchet A, Sebald M. 1979. Transfer of multiple antibiotic resistance between subspecies of *Bacteroides fragilis*. J Infect Dis. 139:97–101. doi: 10.1093/infdis/139.1.97. PMID: 108340.

Ragsdale SW. 2003. Pyruvate ferredoxin oxidoreductase and its radical intermediate. Chem Rev. 103:2333–23346. doi: 10.1021/cr020423e. PMID: 12797832.

Rajamuthiah R, Fuchs BB, Jayamani E, Kim Y, Larkins-Ford J, Conery A, Ausubel FM, Mylonakis E. 2014. Whole animal automated platform for drug discovery against multi-drug resistant *Staphylococcus aureus*. PLoS One. 9:e89189. doi: 10.1371/journal.pone.0089189. PMID: 24586584.

Reed GH, Ragsdale SW, Mansoorabadi SO. 2012. Radical reactions of thiamin pyrophosphate in 2-oxoacid oxidoreductases. Biochim Biophys Acta. 1824:1291–1298. doi: 10.1016/j.bbapap.2011.11.010. PMID: 22178227.

Reed LA, O’Bier NS, Oliver LD, Hoffman PS, Marconi RT. 2018. Antimicrobial activity of amixicile against *Treponema denticola* and other oral spirochetes associated with periodontal disease. J Periodontol. doi: 10.1002/JPER.17-0185. PMID: 29958324.

Reott MA, Parker AC, Rocha ER, Smith CJ. 2009. Thioredoxins in redox maintenance and survival during oxidative stress of *Bacteroides fragilis*. J Bacteriol. 191:3384–3391. doi: 10.1128/JB.01665-08. PMID: 19286811.

Rios-Covian D, Sánchez B, Salazar N, Martínez N, Redruello B, Gueimonde M, de Los Reyes-Gavilán CG. 2015. Different metabolic features of *Bacteroides fragilis* growing in the presence of glucose and exopolysaccharides of bifidobacteria. Front Microbiol. 6:825. doi: 10.3389/fmicb.2015.00825. PMID: 26347720.

Robertson KP, Smith CJ, Gough AM, Rocha ER. 2006. Characterization of *Bacteroides fragilis* hemolysins and regulation and synergistic interactions of HlyA and HlyB. Infect Immun. 74:2304–2316. doi: 10.1128/IAI.74.4.2304-2316.2006. PMID: 16552061.

Rocha ER, Smith CJ. 1995. Biochemical and genetic analyses of a catalase from the anaerobic bacterium *Bacteroides fragilis*. J Bacteriol. 177:3111–3119. doi: 10.1128/jb.177.11.3111-3119.1995. PMID: 7768808.

Rocha ER, Selby T, Coleman JP, Smith CJ. 1996. Oxidative stress response in an anaerobe, *Bacteroides fragilis*: a role for catalase in protection against hydrogen peroxide. J Bacteriol. 178:6895–6903. doi: 10.1128/jb.178.23.6895-6903.1996. PMID: 8955312.

Rocha ER, Smith CJ. 1997. Regulation of *Bacteriodes fragilis katB* mRNA by oxidative stress and carbon limitation. J Bacteriol. 179:7033–7039. doi: 10.1128/jb.179.22.7033-7039.1997. PMID: 9371450.

Rocha ER, Smith CJ. 1998. Characterization of a peroxide-resistant mutant of the anaerobic bacterium *Bacteroides fragilis*. J Bacteriol. 180:5906–5612. doi: 10.1128/JB.180.22.5906-5912.1998. PMID: 9811648.

Rocha ER, Smith CJ. 1999. Role of the alkyl hydroperoxide reductase (*ahpCF*) gene in oxidative stress defense of the obligate Anaerobe *Bacteroides fragilis*. J Bacteriol. 181:5701–5710. doi: 10.1128/JB.181.18.5701-5710.1999. PMID: 10482511.

Rocha ER, Smith CJ. 2004. Transcriptional regulation of the *Bacteroides fragilis* ferritin gene (*ftnA*) by redox stress. Microbiology (Reading). 2004 150:2125–2134. doi: 10.1099/mic.0.26948-0. PMID: 15256555.

Rocha ER, Tzianabos AO, Smith CJ. 2007. Thioredoxin reductase is essential for thiol/disulfide redox control and oxidative stress survival of the anaerobe *Bacteroides fragilis*. J Bacteriol. 2007 Nov;189(22):8015–23. doi: 10.1128/JB.00714-07. PMID: 17873045.

Roy AA, Dhawanjewar AS, Sharma P, Singh G, Madhusudhan MS. 2019. Protein Interaction Z Score Assessment (PIZSA): an empirical scoring scheme for evaluation of protein-protein interactions. Nucleic Acids Res. 47:W331–W337. doi: 10.1093/nar/gkz368. PMID: 31114890.

Ruettinger RT, Olson ST, Boyer RF, Coon MJ. 1974. Identification of the omega-hydroxylase of *Pseudomonas oleovorans* as a nonheme iron protein requiring phospholipid for catalytic activity. Biochem Biophys Res Commun. 57:1011–1017. doi: 10.1016/0006-291x(74)90797-9. PMID: 4830742.

Schofield WB, Zimmermann-Kogadeeva M, Zimmermann M, Barry NA, Goodman AL. 2018. The Stringent Response Determines the Ability of a Commensal Bacterium to Survive Starvation and to Persist in the Gut. Cell Host Microbe. 24:120–132.e6. doi: 10.1016/j.chom.2018.06.002. PMID: 30008292.

Schuetz AN. 2014. Antimicrobial resistance and susceptibility testing of anaerobic bacteria. Clin Infect Dis. 59:698–705. doi: 10.1093/cid/ciu395. PMID: 24867792.

Shafquat Y, Jabeen K, Farooqi J, Mehmood K, Irfan S, Hasan R, Zafar A. 2019. Antimicrobial susceptibility against metronidazole and carbapenem in clinical anaerobic isolates from Pakistan. Antimicrob Resist Infect Control. 8:99. doi: 10.1186/s13756-019-0549-8. PMID: 31210928.

Shilnikova II, Dmitrieva NV. 2015. Evaluation of antibiotic susceptibility of *Bacteroides*, *Prevotella* and *Fusobacterium* species isolated from patients of the N. N. Blokhin Cancer Research Center, Moscow, Russia. Anaerobe. 31:15–18. doi: 10.1016/j.anaerobe.2014.08.003. PMID: 25157873.

Shoun H, Sudo Y, Beppu T. 1985. Subterminal hydroxylation of fatty acids by a cytochrome P-450-dependent enzyme system from a fungus, *Fusarium oxysporum*. J Biochem. 97:755–763. doi: 10.1093/oxfordjournals.jbchem.a135115. PMID: 4019433.

Simon R, Priefer U, Pühler A. 1983. A broad range mobilization system for in vivo genetic engineering: transposon mutagenesis in Gram-negative bacteria. Nat Biotechnol 1:784 –791. 10.1038/nbt1183-784.

Sisson G, Jeong JY, Goodwin A, Bryden L, Rossler N, Lim-Morrison S, Raudonikiene A, Berg DE, Hoffman PS. 2000. Metronidazole activation is mutagenic and causes DNA fragmentation in *Helicobacter pylori* and in *Escherichia coli* containing a cloned *H. pylori* RdxA(+) (Nitroreductase) gene. J Bacteriol. 182:5091–5096. doi: 10.1128/JB.182.18.5091-5096.2000. PMID: 10960092.

Skuce PJ, Fairweather I. 1990. The effect of the hydrogen ionophore closantel upon the pharmacology and ultrastructure of the adult liver fluke *Fasciola hepatica*. Parasitol Res. 76:241–250. doi: 10.1007/BF00930821. PMID: 2315284.

Smith CJ, Rogers MB, McKee ML. 1992. Heterologous gene expression in *Bacteroides fragilis*. Plasmid 27:141–154. 10.1016/0147-619X(92)90014-2. PMID: 1615064.

Smith CJ, Rollins LA, Parker AC. 1995. Nucleotide sequence determination and genetic analysis of the Bacteroides plasmid, pBI143. Plasmid 34:211–222. 10.1006/plas.1995.0007. PMID: 8825374

Snydman DR, Jacobus NV, McDermott LA, Goldstein EJ, Harrell L, Jenkins SG, Newton D, Patel R, Hecht DW. 2017. Trends in antimicrobial resistance among *Bacteroides* species and *Parabacteroides* species in the United States from 2010-2012 with comparison to 2008-2009. Anaerobe. 43:21–26. doi: 10.1016/j.anaerobe.2016.11.003. PMID: 27867083.

Soo PC, Horng YT, Lai MJ, Wei JR, Hsieh SC, Chang YL, Tsai YH, Lai HC. 2007. Pirin regulates pyruvate catabolism by interacting with the pyruvate dehydrogenase E1 subunit and modulating pyruvate dehydrogenase activity. J Bacteriol. 189:109–118. doi: 10.1128/JB.00710-06. PMID: 16980458.

Stephenson K, Yamaguchi Y, Hoch JA. 2000. The mechanism of action of inhibitors of bacterial two-component signal transduction systems. J Biol Chem. 275:38900–38904. doi: 10.1074/jbc.M006633200. PMID: 10978341.

Stokes JM, Lopatkin AJ, Lobritz MA, Collins JJ. 2019. Bacterial Metabolism and Antibiotic Efficacy. Cell Metab. 30:251–259. doi: 10.1016/j.cmet.2019.06.009. Epub 2019 Jul 3. PMID: 31279676.

Sund CJ, Rocha ER, Tzianabos AO, Wells WG, Gee JM, Reott MA, O’Rourke DP, Smith CJ. 2008. The *Bacteroides fragilis* transcriptome response to oxygen and H_2_O_2_: the role of OxyR and its effect on survival and virulence. Mol Microbiol. 67:129–142. doi: 10.1111/j.1365-2958.2007.06031.x. PMID: 18047569.

Talà A, Damiano F, Gallo G, Pinatel E, Calcagnile M, Testini M, Fico D, Rizzo D, Sutera A, Renzone G, Scaloni A, De Bellis G, Siculella L, De Benedetto GE, Puglia AM, Peano C, Alifano P. 2018. Pirin: A novel redox-sensitive modulator of primary and secondary metabolism in *Streptomyces*. Metab Eng. 48:254–268. doi: 10.1016/j.ymben.2018.06.008. PMID: 29944936.

Tang YP, Dallas MM, Malamy MH. 1999. Characterization of the Batl (Bacteroides aerotolerance) operon in *Bacteroides fragilis*: isolation of a *B. fragilis* mutant with reduced aerotolerance and impaired growth in *in vivo* model systems. Mol Microbiol. 32:139–149. doi: 10.1046/j.1365-2958.1999.01337.x. PMID: 10216867.

Tran TB, Cheah SE, Yu HH, Bergen PJ, Nation RL, Creek DJ, Purcell A, Forrest A, Doi Y, Song J, Velkov T, Li J. 2016. Anthelmintic closantel enhances bacterial killing of polymyxin B against multidrug-resistant *Acinetobacter baumannii*. J. Antibiot. 69:415–421. doi: 10.1038/ja.2015.127. PMID: 26669752.

Turrens JF, Newton CL, Zhong L, Hernandez FR, Whitfield J, Docampo R. 1999. Mercaptopyridine-N-oxide, an NADH-fumarate reductase inhibitor, blocks *Trypanosoma cruzi* growth in culture and in infected myoblasts. FEMS Microbiol Lett. 175:217–221. doi: 10.1111/j.1574-6968.1999.tb13623.x. PMID: 10386371.

Van Den Bossche H, Verhoeven H, Vanparijs O, Lauwers H, Thienpont D. 1979. Closantel, a new antiparasitic hydrogen ionophore. Arch Int Physiol Biochim. 87:851–853. PMID: 93944.

Vanhanen S, West M, Kroon JT, Lindner N, Casey J, Cheng Q, Elborough KM, Slabas AR. 2000. A consensus sequence for long-chain fatty-acid alcohol oxidases from *Candida* identifies a family of genes involved in lipid omega-oxidation in yeast with homologues in plants and bacteria. J Biol Chem. 275:4445–4452. doi: 10.1074/jbc.275.6.4445. PMID: 10660617.

Warren CA, van Opstal E, Ballard TE, Kennedy A, Wang X, Riggins M, Olekhnovich I, Warthan M, Kolling GL, Guerrant RL, Macdonald TL, Hoffman PS. 2012. Amixicile, a novel inhibitor of pyruvate: ferredoxin oxidoreductase, shows efficacy against *Clostridium difficile* in a mouse infection model. Antimicrob Agents Chemother. 56:4103–4111. PMID: 22585229.

Wendler WM, Kremmer E, Förster R, Winnacker EL. 1997. Identification of pirin, a novel highly conserved nuclear protein. J Biol Chem. 272:8482–8489. doi: 10.1074/jbc.272.13.8482. PMID: 9079676.

Williamson RL, Metcalf RL. 1967. Salicylanilides: a new group of active uncouplers of oxidative phosphorylation. Science. 158:1694–1695. doi: 10.1126/science.158.3809.1694. PMID: 4228706.

Yamamoto M, Arai H, Ishii M, Igarashi Y. 2006. Role of two 2-oxoglutarate:ferredoxin oxidoreductases in *Hydrogenobacter thermophilus* under aerobic and anaerobic conditions. FEMS Microbiol Lett. 263:189–193. doi: 10.1111/j.1574-6968.2006.00415.x. PMID: 16978355.

Yamamoto M, Ikeda T, Arai H, Ishii M, Igarashi Y. 2010. Carboxylation reaction catalyzed by 2-oxoglutarate:ferredoxin oxidoreductases from *Hydrogenobacter thermophilus*. Extremophiles. 14:79–85. doi: 10.1007/s00792-009-0289-4. PMID: 19894084.

Yoshikawa R, Yanagi H, Hashimoto-Tamaoki T, Morinaga T, Nakano Y, Noda M, Fujiwara Y, Okamura H, Yamamura T. 2004. Gene expression in response to anti-tumour intervention by polysaccharide-K (PSK) in colorectal carcinoma cells. Oncol Rep. 12:1287–1293. PMID: 15547752.

Young M, Chojnacki M, Blanchard C, Cao X, Johnson WL, Flaherty D, Dunman PM. 2023. Genetic Determinants of *Acinetobacter baumannii* Serum-Associated Adaptive Efflux-Mediated Antibiotic Resistance. Antibiotics (Basel). 12:1173. doi: 10.3390/antibiotics12071173. PMID: 37508269.

Yun NR, Arai H, Ishii M, Igarashi Y. 2001. The genes for anabolic 2-oxoglutarate: ferredoxin oxidoreductase from *Hydrogenobacter thermophilus* TK-6. Biochem Biophys Res Commun. 282:589–594. doi: 10.1006/bbrc.2001.4542. PMID: 11401501.

Yun NR, Yamamoto M, Arai H, Ishii M, Igarashi Y. 2002. A novel five-subunit-type 2-oxoglutalate:ferredoxin oxidoreductases from *Hydrogenobacter thermophilus TK*-6. Biochem Biophys Res Commun. 292:280–286. doi: 10.1006/bbrc.2002.6651. PMID: 11890705.

Zhang G, Dai J, Lu Z, Dunaway-Mariano D. 2003. The phosphonopyruvate decarboxylase from *Bacteroides fragilis*. J Biol Chem. 278(42):41302–41308. doi: 10.1074/jbc.M305976200. PMID: 12904299.

