## Supplemental File S5.xlsx for "Novel roles for pirin proteins and a 2-ketoglutarate: ferredoxin oxidoreductase ortholog in *Bacteroides fragilis* central metabolism and comparison of metabolic mutants susceptibility to metronidazole and amixicile"

**Supplemental File S1**

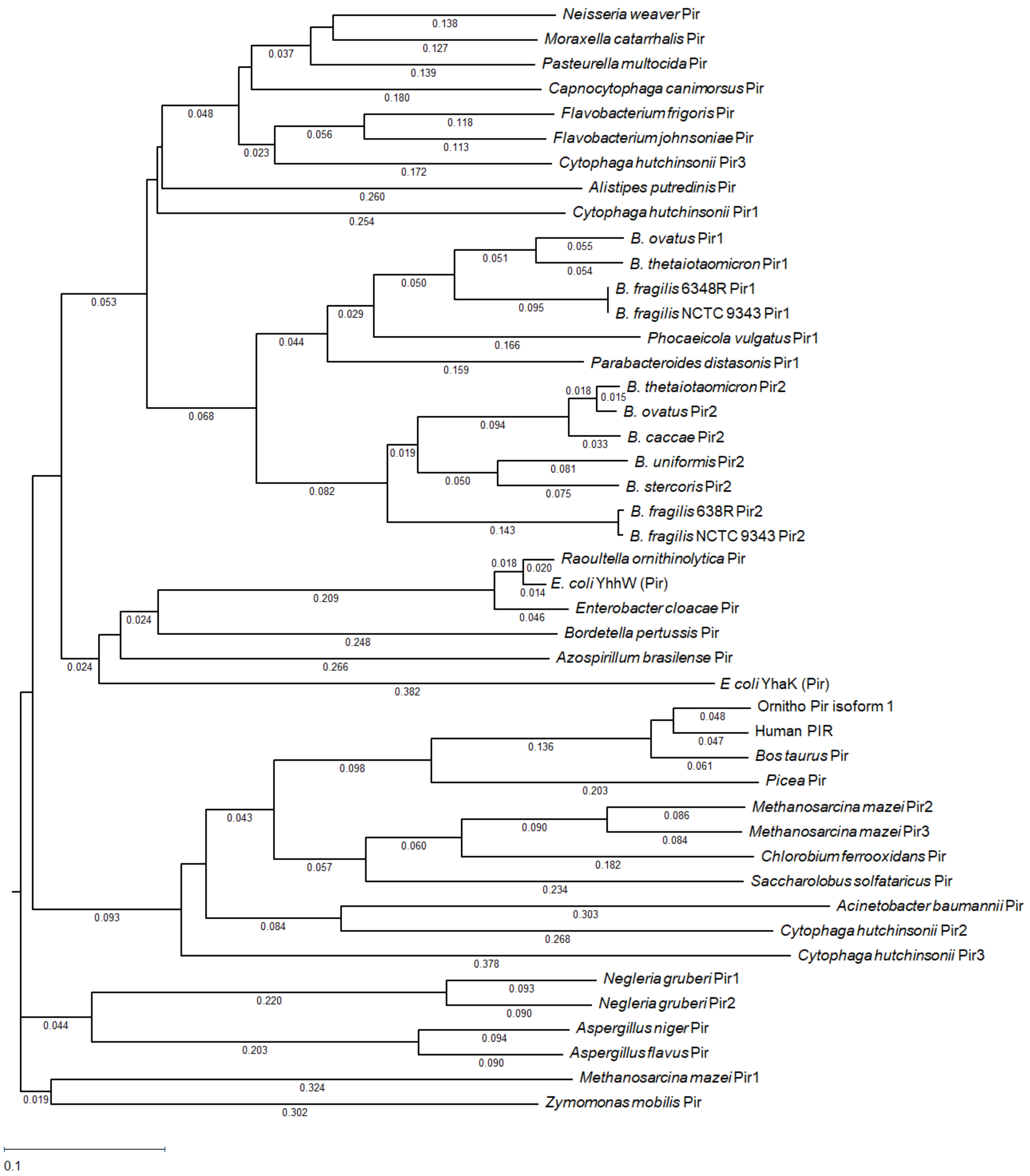

**Supplemental File S1**. Phylogenetic relationship of 49 Pirin-like protein homologues representatives from Bacteroidetes, and from other prokaryotes, eukaryotes, and Archaea. The unrooted phylogenetic tree was constructed from multiple amino acid sequences alignment based on ClustalW algorithm in MegAlign Pro program of Lasergene Version 17.5.0 (DNASTAR, Inc). The built-in Neighbor Joining method was used to calculate the distances between all pairs and the calculated relative distance values are depicted in parenthesis following each tree entry name. GenBank accession numbers and abbreviations are as follow: Acinetobacter baumanii AC12 Pir (AHX26982). *Alistipes putredinis* DSM 17216 Pir (EDS02825). *Aspergillus flavus* AF70 Pir (KOC10185). *Aspergillus niger* ATCC 1015 Pir (EHA28466). *Azospirillum brasilense* Pir (TWA73044). *Bacteroides fragilis* 638R Pir1 (CBW23516), Pir2 (CBW22007). *Bacteroides fragilis* NCTC 9343 Pir1 (CAH08727), Pir2 (CAH07099). *Bacteroides ovatus* ATCC 8483 Pir1 (EDO13012), Pir2 (EDO09940). *Bacteroides stercoris* ATCC 43183 Pir2 (EDS16761). *Bacteroides thetaiotaomicron* VPI-5482 Pir1 (AAO75294), Pir2 (AAO76683). *Bacteroides uniformis* ATCC 8492 Pir2 (EDO51902). *Bordetella pertussis* CS Pir (AEE66951). *Bos taurus* Pir (NC_037357). *Capnocytophaga canimorsus* Cc5 Pir (AEK22609). *Chlorobium ferrooxidans* DSM 1303 Pir (EAT59973). *Cytophaga hutchinsonii* ATCC 33406 Pir1 (ABG57630), Pir2 (ABG58934), Pir3 (ABG59359), and Pir4 (ABG60251). *Enterobacter cloacae* subsp. cloacae ATCC 13047 Pir (ADF64329). *Escherichia coli* str K12 MG1655 YahK (NP_417577), YhhW (NC_000913). *Flavobacterium frigoris* PS1 Pir (EIA09520). *Flavobacterium johnsoniae* UW101 Pir (ABQ04767). Human Pir: *Homo sapiens* (NP_003653). *Methanobactins mazei* Go1 Pir1 (AAM32493), Pir2 (AAM32593) and Pir3 (AAM32603). *Moraxella catarrhalis* 7169 Pir (EGE09966). *Naegleria gruberi* Pir1 (EFC43129), Pir2 (EFC40196). *Neisseria weaveri* ATCC 51223 Pir (EGV38178). Ornitho Pir Isoform 1: *Ornithorhynchus anatinus* pirin isoform X1 (XP_003431123). *Parabacteroides distasonis* ATCC 8503 Pir1 (ABR41886). *Pasteurella multocida* 36950 Pir (AET16501). *Phocaeicola vulgatus* ATCC 8482 Pir1 (ABR40270). Picea: *Picea sitchensis* Pir (ABK27098). *Raoultella ornithinolytica* 10-5246 Pir (EHT04992). *Saccharolobus solfataricus* Pir (AKA74406). *Zymomonas mobilis* subsp. mobilis ZM4 Pir (AAV89961).

### Supplemental File S2

```
                        1 ........10........20........30........40........50........60........70........80
E. coli YhaK            1 -MITTRTARQCGQADYGWLQARYTFSFGHYFDPKLLGYASLRVLNQEVLAPGAAFQPRTYPKVDILNVILDG-EAEYRDS
BF638R_1469 Pir2        1 MKKVVHRSDTRGRSVYDWLDSHHSFSFDEYYNPERVHFGALRVLNDDRVAPGEGFQPHPHKNMEIVSIPLKG-LLAHGDS
BF638R_3039 Pir1        1 MKTVVDKASSRGYFNHGWLKTHHTFSFANYYNPSRMHFGVLRVLNDDSVDPEMGFDTHPHQNMEVISIPLKG-YLRHGDS
BT_0187 Pir1            1 MKKVIDRASSRGYFNHGWLKTHHTFSFANYYNPERIHFGALRVLNDDSVDPSMGFDTHPHKNMEVISIPLKG-YLRHGDS
BT_1576 Pir2            1 MKKVIHKADTRGHSQYDWLDSYHTFSFDEYFDSDRINFGALRVLNDDKVAPGEGFQTHPHKNMEIISIPLKG-HLQHGDS
E. coli YhhW            1 -MIYLRKANERGHANHGWLDSWHTFSFANYYDPNFMGFSALRVINDDVIEAGQGFGTHPHKDMEILTYVLEG-TVEHQDS
Human_Pir               1 --MGSSKKVTLSVLSREQSEGVGARVRRSIGRPELKNLDPFLLFDEFKGGRPGGFPDHPHRGFETVSYLLEGGSMAHEDF
                                         R        R        E K                     * *
BF_Pir1 binding sites                   NHG         FAN Y P  M     RV          G DTHPHQNM
BF_Pir2 binding sites                              SFDEYY P  V     R            FQPHPHKNM
                       81 ........90.......100.......110.......120.......130.......140.......150.......160
E. coli YhaK           79 EGNHVQASAGEALLLSTQPGVSYSEHNLSKDKPLTRMQLWLDAC--PQRENPLIQKLALNMGK-QQLIASPEGAMG--SL
BF638R_1469 Pir2       80 KKNSRTITVGDIQVMSAGTGIYHSEMNGSKSEPVEFLQIWIIPK--ERNTHPLYQDYDIRGLLKKDELAFILSPNGSTPA
BF638R_3039 Pir1       80 VKNTRTITPGDIQVMSTGKGIFHSEYNGSDKEQLEFLQIWVFPR--IENTEPEYNNYDIRPLLKRNELALIISPDGKVPA
BT_0187 Pir1           80 VQNTKTITPGDIQVMSTGSGIYHSEYNDSKEEQLEFLQIWVFPR--IENTKPEYNNFDIRPLLKPNELSLFISPNGKTPA
BT_1576 Pir2           80 KKNSRIITVGEIQTMSAGTGIFHSEVNASPVEPVEFLQIWIMPR--ERNTHPVYKDFSIKELERPNELAVIVSPDGSTPA
E. coli YhhW           79 MGNKEQVPAGEFQIMSAGTGIRHSEYNPNSTERLHLYQIWIMPE--ENGITPRYEQRRFDAVQGKQLVLSPDARDG--SL
Human_Pir              79 CGHTGKMNPGDLQWMTAGRGILHAEMPCSE-EPAHGLQLWVNLRSSEKMVEPQYQELKSEEIPKPSKDGVTVAVISGEAL
                                                * *

BF_Pir1 binding sites                          F                 W F
BF_Pir2 binding sites                       T  Y                    I     R
                      161 .......170.......180.......190.......200.......210.......220.......230.......240
E. coli YhaK          154 QLRQQVWLHHIVLDKGESANF-----QLHGPR--AYLQSIHGKFHALTH-HEEKAALTCGDGAFIRDEANITLVADSPLR
BF638R_1469 Pir2      158 KLLQDTWFSMGEIGAGKTVEY-----TLHGTDMGVYVFLIEGEVKIDD------VILTRRDGLGISEIENFEIETLKDSK
BF638R_3039 Pir1      158 SIKQDAWFSMGTFDAGKSFEY-----KLHQEGNGVYLFIIEGDVEVAG------NRLSRRDGIGLWDTKSFKVEITQEAT
BT_0187 Pir1          158 SIKQDAWFSMGDFDTERTIEY-----CMHQEGNGAYLFVIEGEISVAD------EHLAKRDGIGIWDTKSFSIRATKGTK
BT_1576 Pir2          158 SLLQDTWFSIGKVEAGKKLGY-----HLHQSHGGVYIFLIEGEIVVDG------EVLKRRDGMGVYDTKSFELETLKDSH
E. coli YhhW          155 KVPQDMELYRWALLKDEQSVH-----QIAAERR-VWIQVVKGNVTIN------GVKASTSDGLAIWDEQAISIHADSDSE
Human_Pir             158 GIKSKVYTRTPTLYLDFKLDPGAKHSQPIPKGWTSFIYTISGDVYIGPDDAQQKIEPHHTAVLGEGDSVQVENKDPKRSH

                      241 .......250.......260.......270.......280.......290...
E. coli YhaK          226 ALLIDLPV---------------------------------------------
BF638R_1469 Pir2      227 ILLIEVPM---------------------------------------------
BF638R_3039 Pir1      227 LLLMEVPMR--------------------------------------------
BT_0187 Pir1          227 LLVMEVPM---------------------------------------------
BT_1576 Pir2          227 ILLIEVPM---------------------------------------------
E. coli YhhW          223 VLLFDLPPV--------------------------------------------
Human_Pir             238 FVLIAGEPLREPVIQHGPFVMNTNEEISQAILDFRNAKNGFERAKTWKSKIGN
```

**Supplemental File S2**. Multiple alignment of predicted pirin (Pir) proteins from *Bacteroides fragilis* 636R (BF638R) Pir1 (CBW23516) and Pir2 (CBW22007), *Bacteroides thetaiotaomicron* VPI 5482 (BT), Pir1 (AAO75294) and Pir2 (AAO76683) with pirin proteins from *Escherichia coli* str. K-12 substr. MG1655 YhaK (NP_417577) and YhhW (NP_417896), and Human Pir (NP_003653). The conserved amino acid residues (>50 %) are labeled with black boxes. Semiconserved amino acid substitutions are depicted by grey boxes. Alignment of the protein sequences was performed using ClustalW algorithm from the MegAlign Pro component of the DNASTAR Lasergene software Version: 17.5.0 (DNASTAR, Inc. Madison, WI). The N-terminal domain residues His56, His58, His101, and Glu103 ligands of the iron centre of human pirin (Pang et al., 2004; Liu et al., 2013) are depicted by an asterisk below the sequence. The amino acids interacting with >50% of the binder proteins are denoted in blue bold-face font below the sequence. Orange bold-face font denotes binding interaction with <49% to >25% of binder proteins, and black regular font below the sequence represents amino acids binding <25% to >10% of binder proteins. The green-font represents human pirin amino acids Arg14, Arg23, Glu32, and Lys34 interacting with nuclear factor. NF-κB (Liu et al., 2013).

#### References

Liu F, Rehmani I, Esaki S, Fu R, Chen L, de Serrano V, Liu A. 2013. Pirin is an iron-dependent redox regulator of NF-κB. Proc Natl Acad Sci U S A. 110:9722-9727. doi: 10.1073/pnas.1221743110. PMID: 23716661.

Pang H, Bartlam M, Zeng Q, Miyatake H, Hisano T, Miki K, Wong LL, Gao GF, Rao Z. 2004. Crystal structure of human pirin: an iron-binding nuclear protein and transcription cofactor. J Biol Chem. 279:1491-1498. doi: 10.1074/jbc.M310022200. PMID: 14573596.

### Supplemental File S3

**Deduced peptide sequences of pTRG/genomic library clones interacting with Pir1.**

| pTRG clone number | Deduced Peptide Sequence | Homology to: | Locus-tag |
|---|---|---|---|
| 1 | DQLANGFHYTVGFNLSDYQSEVTKFDNESKELGNWYVGQKQGEI | TonB-dependent receptor | BF638R_3226 |
| 3 | SPAPVGYYSTGSWLLFGRSYFVCLKSDCPYTYRDFEKTVFPNQEQI | Lipopolysaccharide kinase InaA family protein | BF638R_0173 |
| 6 | DRTKEPGANGEPLYLDVKDCFYGAENAPVIVGGRYGLGSKDTTPAQIIAV FKNLAMPMPKNHFTIGIVDDVTFTSLPQEAEIALGGEGMFEAKFYGLGAD GTVGANKNSVKIIGDNTDKHCQAYFSYDSKKSGGFTCSHLRFGDDRSAKK FDHGILKFKWLQQIVDCIYLETLDGIFRIGGCKHNQGLHH | Pyruvate:ferredoxin (flavodoxin) oxidoreductase | BF638R_3194 |
| 11 | DRLEDTQFFEEEIHAAVKAAEDAGTYVMVHVYVPRAIQRAIHAGVKSIEH GHLIDEPTMQLI | amidohydrolase family protein | BF638R_1629 |
| 12 | DLLLMSEIKGYVNNPQYYFESRDDTHRKAI | Hypothetical protein | BF638R_1398 |
| 13 | DRTKEPGANGEPLYLDVKDCFYGAENAPVIVGGRYGLGSKDTTPAQIIAV FKNLAMPMPKNHFTIGIVDDVTFTSLPQEAEIALGGEGMFEAKFYGLGAD GTVGANKNSVKIIGDNTDKHCQAYFSYDSKKSGGFTCSHLRFGDDRSAKK FDHGILKFKWLQQIVDCIYLETLDGIFRIGGCKHNQGLHH | Pyruvate:ferredoxin (flavodoxin) oxidoreductase | BF638R_3194 |
| 15 | DRTKEPGANGEPLYLDVKDCFYGAENAPVIVGGRYGLGSKDTTPAQIIAV FKNLAMPMPKNHFTIGIVDDVTFTSLPQEAEIALGGEGMFEAKFYGLGAD GTVGANKNSVKIIGDNTDKHCQAYFSYDSKKSGGFTCSHLRFGDDRSAKK FDHGILKFKWLQQIVDCIYLETLDGIFRIGGCKHNQGLHH | Pyruvate:ferredoxin (flavodoxin) oxidoreductase | BF638R_3194 |
| 19 | DLLLMSEIKGYVNNPQYYFESRDDTHRKAI | Hypothetical protein | BF638R_1398 |
| 21 | SVGRYTVLRGRDSCCR | No significant similarity found | Non-canonical peptide |
| 22 | SINSTATPIASITASRRASGSPIRVTTSRL | No significant similarity found | Non-canonical peptide |
| 24 | TFCGECFFCRHGYVNNCTDPDGGWALGCRIDGGQAEYVRV | zinc-binding alcohol dehydrogenase | BF638R_1292 |
| 25 | SFINVRNKRICQQSPILFRIPGRYTPQGNRSPI | No significant similarity found | Non-canonical peptide |

**Deduced peptide sequences of pTRG/genomic library clones interacting with Pir2. First experiment**

| pTRG clone number | Deduced Peptide Sequence | Homology to: | Locus-tag |
|---|---|---|---|
| 1 | SICILFLF | No significant similarity found | Non-canonical peptide |
| 2 | SICILFLF | No significant similarity found | Non-canonical peptide |
| 3 | TFCGECFFCRHGYVNNCTDPDGGWALGCRIDGGQAEYVRV | zinc-binding alcohol dehydrogenase | BF638R_1292 |
| 12 | SFFILLNVFSA | No significant similarity found | Non-canonical peptide |
| 18 | SICILFLF | No significant similarity found | Non-canonical peptide |
| 22 | SFFILLNVFSA | No significant similarity found | Non-canonical peptide |

**Deduced peptide sequences of pTRG/genomic library clones interacting with Pir2. Second experiment**

| pTRG clone number | Deduced Peptide Sequence | Homology to: | Locus-tag |
|---|---|---|---|
| 2 | TFCGECFFCRHGYVNNCTDPDGGWALGCRIDGGQAEYVRV | zinc-binding alcohol dehydrogenase | BF638R_1292 |
| 14 | SLVLLLSVKQW | No significant similarity found | Non-canonical peptide |
| 20 | TFCGECFFCRHGYVNNCTDPDGGWALGCRIDGGQAEYVRV | zinc-binding alcohol dehydrogenase | BF638R_1292 |
| 21 | TFCGECFFCRHGYVNNCTDPDGGWALGCRIDGGQAEYVRV | zinc-binding alcohol dehydrogenase | BF638R_1292 |
| 31 | TFCGECFFCRHGYVNNCTDPDGGWALGCRIDGGQAEYVRV | zinc-binding alcohol dehydrogenase | BF638R_1292 |

**Supplemental file S3**: Deduced amino acid sequences of Two-Hybrid System partial genomic library from *B. fragilis* 638R genome inserted in the pTRG plasmid interacting with pirin 1 (Pir1) or Pirin 2 (Pir2). The pTRG forward primer (Table S1) was used in the sequencing reactions to obtain nucleotide sequence of the inserted DNA cloned into the BamHI site downstream of the RNAP α-subunit.. Non-canonical peptide: in-frame translation of non-annotated DNA reading frame. The ligation of Sau3AI partial library into the pTRG BamHI site contains multiple random self-ligated Sau3AI DNA fragments. Only the deduced peptide expressed in-frame with RNA-α-subunit is shown.

**Supplemental file S4**

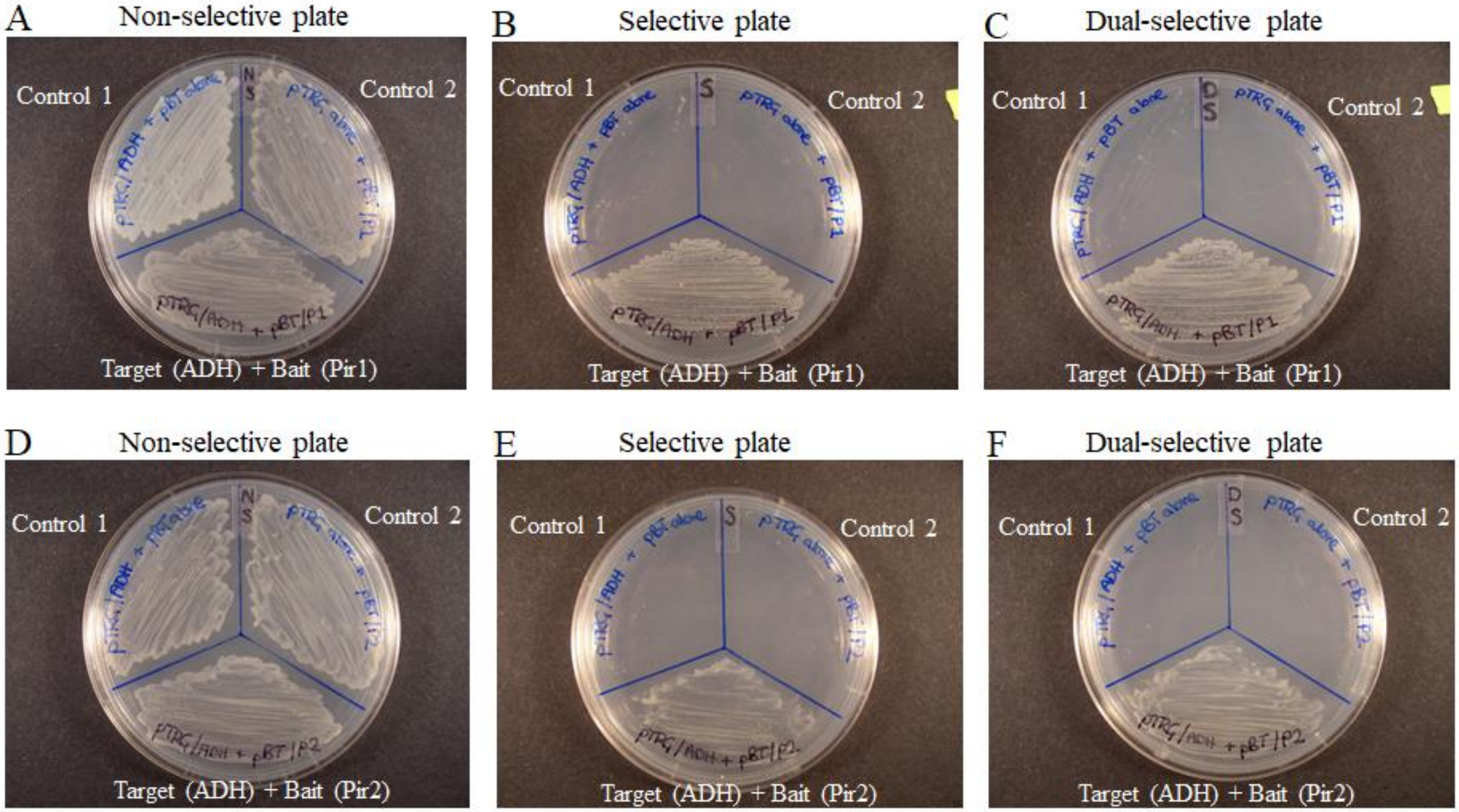

**Supplemental file S4.** Bacterial two-hybrid system assay showing protein-protein interaction between Pir1 and Zn-ADH (A, B, C), and between Pir2 and Zn-ADH (D, E, F). *E. coli* reporter strain was cotransformed with pBT/Pir1 (bait) and pTRG/ADH (prey) or pBT/Pir2 (bait) and pTRG/ADH (prey) constructs and they were grown on control non-selective plate (A, D), selective plate (B, E), and dual selective plate(C, F). Self-activation controls are *E. coli* two-hybrid system reporter strain carrying the following constructs: control 1, empty "bait" (pBT alone) cotransformed with loaded "prey" (pTRG/ADH); control 2, loaded "bait" (pBT/Pir1 or pBT/Pir2) cotransformed with empty "prey" (pTRG alone). See Materials and Methods for details on the bacterial two-hybrid system assay.

### Supplemental File S6.

Growth conditions/strains — Short chain fatty acids concentration (mM)

| Anaerobic mid-log | Succinate | Lactate | Formate | Acetate | Propionate | Isobutyrate | Butyrate | Isovalerate | Phenylacetate | % Glucose used |
|---|---|---|---|---|---|---|---|---|---|---|
| 638R | 1.42 (± 0.50) | 0.00 (± 0.00) | 0.84 (± 1.07) | 1.83 (± 0.37) | 0.00 (± 0.00) | 1.08 (± 1.53) | 0.21 (± 0.30) | 0.67 (± 0.15) | 0.35 (± 0.01) | 8.88 (± 1.46) |
| Δ*pir1* | 1.30 (± 0.21) | 0.06 (± 0.08) | 0.79 (± 1.11) | 0.93 (± 0.98) | 0.51 (± 0.72) | 1.56 (± 2.20) | 0.22 (± 0.30) | 0.07 (± 0.01) | 0.00 (± 0.00) | 8.30 (± 2.64) |
| Δ*pir2* | 1.82 (± 0.18) | 0.15 (± 0.22) | 1.44 (± 0.85) | 1.68 (± 1.12) | 0.14 (± 0.09) | 1.32 (± 1.86) | 0.04 (± 0.06) | 0.04 (± 0.05) | 0.00 (± 0.00) | 9.71 (± 3.08) |
| Δ*pir1* Δ*pir2* | 1.67 (± 0.64) | 0.31 (± 0.43) | 0.92 (± 0.82) | 0.98 (± 0.92) | 0.18 (± 0.25) | 0.99 (± 1.38) | 0.72 (± 0.38) | 0.04 (± 0.04) | 0.00 (± 0.00) | 10.43 (± 0.10) |

| Anaerobic mid-log + DP | Succinate | Lactate | Formate | Acetate | Propionate | Isobutyrate | Butyrate | Isovalerate | Phenylacetate | % Glucose used |
|---|---|---|---|---|---|---|---|---|---|---|
| 638R | 1.12 (± 0.15) | 0.32 (± 0.30) | 0.95 (± 1.34) | 0.87 (± 0.46) | 0.51 (± 0.71) | 0.00 (± 0.00) | 0.00 (± 0.00) | 0.29 (± 0.41) | 0.00 (± 0.00) | 8.19 (± 2.26) |
| Δ*pir1* | 1.00 (± 0.01) | 0.26 (± 0.24) | 1.35 (± 1.91) | 1.64 (± 0.71) | 0.14 (± 0.19) | 0.15 (± 0.21) | 0.00 (± 0.00) | 0.00 (± 0.00) | 0.00 (± 0.00) | 10.49 (± 6.45) |
| Δ*pir2* | 1.14 (± 0.22) | 0.42 (± 0.45) | 1.29 (± 1.82) | 1.24 (± 1.12) | 0.02 (± 0.02) | 2.28 (± 1.87) | 0.00 (± 0.00) | 0.03 (± 0.04) | 0.00 (± 0.00) | 8.92 (± 0.72) |
| Δ*pir1* Δ*pir2* | 0.86 (± 0.27) | 0.29 (± 0.36) | 0.92 (± 1.30) | 0.78 (± 0.78) | 0.25 (± 0.35) | 2.16 (± 2.58) | 0.47 (± 0.66) | 0.00 (± 0.00) | 0.00 (± 0.00) | 7.41 (± 1.62) |

| Oxygen for 1h | Succinate | Lactate | Formate | Acetate | Propionate | Isobutyrate | Butyrate | Isovalerate | Phenylacetate | % Glucose used |
|---|---|---|---|---|---|---|---|---|---|---|
| 638R | 1.93 (± 0.64) | 0.22 (± 0.31) | 0.95 (± 0.69) | 1.23 (± 0.99) | 0.14 (± 0.20) | 1.15 (± 1.62) | 0.66 (± 0.38) | 0.02 (± 0.02) | 0.01 (± 0.02) | 12.46 (± 1.94) |
| Δ*pir1* | 1.81 (± 0.47) | 0.23 (± 0.33) | 0.85 (± 0.84) | 1.17 (± 1.15) | 0.00 (± 0.00) | 0.87 (± 1.24) | 0.20 (± 0.29) | 0.05 (± 0.03) | 0.01 (± 0.14) | 12.16 (± 3.50) |
| Δ*pir2* | 2.04 (± 0.11) | 0.45 (± 0.23) | 1.07 (± 1.03) | 0.87 (± 0.77) | 0.14 (± 0.20) | 1.63 (± 2.29) | 0.53 (± 0.66) | 0.03 (± 0.03) | 0.01 (± 0.02) | 12.73 (± 3.35) |
| Δ*pir1* Δ*pir2* | 1.67 (± 0.37) | 0.26 (± 0.12) | 1.01 (± 0.54) | 0.94 (± 1.32) | 0.30 (± 0.43) | 1.10 (± 1.46) | 0.64 (± 0.19) | 0.02 (± 0.02) | 0.01 (± 0.02) | 11.10 (± 3.65) |

| Anaerobic for 24 h | Succinate | Lactate | Formate | Acetate | propionate | isobutyrate | butyrate | isovalerate | phenylacetate | % Glucose used |
|---|---|---|---|---|---|---|---|---|---|---|
| 638R | 14.01 (± 1.08) | 3.50 (± 0.45) | 6.00 (± 0.07) | 15.38 (± 0.48) | 3.57 (± 0.02) | 0.94 (± 0.01) | 0.00 (± 0.00) | 0.29 (± 0.01) | 0.14 (± 0.00) | 79.67 (± 0.25) |
| Δ*pir1* | 13.71 (± 0.59) | 4.40 (± 0.27) | 6.07 (± 0.07) | 15.36 (± 0.50) | 3.48 (± 0.02) | 0.97 (± 0.04) | 0.00 (± 0.00) | 0.30 (± 0.00) | 0.14 (± 0.00) | 79.56 (± 0.51) |
| Δ*pir2* | 13.24 (± 1.01) | 5.33 (± 0.62) | 6.16 (± 0.08) | 15.02 (± 0.59) | 3.87 (± 0.50) | 1.96 (± 1.43) | 0.00 (± 0.00) | 0.22 (± 0.08) | 0.09 (± 0.08) | 78.02 (± 1.24) |
| Δ*pir1* Δ*pir2* | 13.72 (± 0.71) | 4.54 (± 0.29) | 4.96 (± 0.41) | 14.74 (± 0.04) | 3.82 (± 0.53) | 1.36 (± 0.53) | 0.16 (± 0.22) | 0.26 (± 0.05) | 0.09 (± 0.08) | 80.47 (± 0.16) |

| Oxygen for 24h | Succinate | Lactate | Formate | Acetate | propionate | isobutyrate | butyrate | isovalerate | phenylacetate | % Glucose used |
|---|---|---|---|---|---|---|---|---|---|---|
| 638R | 3.31 (± 0.30) | 9.92 (± 1.68) | 0.00 (± 0.00) | 2.88 (± 0.00) | 1.65 (± 0.23) | 0.82 (± 0.19) | 0.00 (± 0.00) | 0.00 (± 0.00) | 0.05 (± 0.03) | 26.20 (± 0.30) |
| Δ*pir1* | 3.05 (± 0.40) | 10.47 (± 1.57) | 0.00 (± 0.00) | 2.64 (± 0.83) | 1.53 (± 0.99) | 0.90 (± 0.46) | 0.62 (± 0.32) | 0.00 (± 0.00) | 0.18 (± 0.18) | 24.78 (± 2.16) |
| Δ*pir2* | 3.02 (± 0.15) | 11.17 (± 1.29) | 0.00 (± 0.00) | 3.15 (± 0.81) | 1.36 (± 1.05) | 0.67 (± 0.40) | 0.00 (± 0.00) | 0.34 (± 0.58) | 0.15 (± 0.09) | 24.41 (± 5.66) |
| Δ*pir1* Δ*pir2* | 2.72 (± 0.27) | 10.90 (± 1.55) | 0.00 (± 0.00) | 2.69 (± 0.46) | 1.15 (± 1.01) | 1.15 (± 0.77) | 0.63 (± 0.54) | 0.00 (± 0.00) | 0.17 (± 0.09) | 28.02 (± 1.27) |

| Oxygen for 24h + DP | Succinate | Lactate | Formate | Acetate | propionate | isobutyrate | butyrate | isovalerate | phenylacetate | % Glucose used |
|---|---|---|---|---|---|---|---|---|---|---|
| 638R | 2.21 (± 1.43) | 2.62 (± 3.66) | 0.12 (± 0.17) | 2.62 (± 1.70) | 0.00 (± 0.00) | 0.04 (± 0.06) | 0.00 (± 0.00) | 0.00 (± 0.00) | 0.00 (± 0.00) | 11.46 (± 5.96) |
| Δ*pir1* | 2.07 (± 1.38) | 2.67 (± 3.77) | 0.00 (± 0.00) | 1.90 (± 2.29) | 0.00 (± 0.00) | 0.03 (± 0.05) | 0.19 (± 0.27) | 0.00 (± 0.00) | 0.14 (± 0.17) | 11.42 (± 6.46) |
| Δ*pir2* | 2.23 (± 1.50) | 2.34 (± 3.10) | 0.00 (± 0.00) | 2.46 (± 1.52) | 0.00 (± 0.00) | 0.32 (± 0.20) | 0.11 (0.16) | 0.00 (± 0.00) | 0.07 (± 0.10) | 9.44 (± 3.58) |
| Δ*pir1* Δ*pir2* | 2.18 (± 1.47) | 2.93 (± 3.05) | 0.00 (± 0.00) | 2.54 (± 1.23) | 0.00 (± 0.00) | 0.13 (± 0.19) | 0.37 (± 0.53) | 0.00 (± 0.00) | 0.00 (± 0.00) | 10.14 (± 4.20) |

**Supplemental file S6**. Short chain fatty acids concentrations (mM) in the culture supernatant of *B. fragilis* strains grown in PYG broth in conditions described in the panel, respectively. Anaerobic mid-log cultures were split, and one half was exposed to atmospheric air for 1h or 24h in aerobic shaker incubator at 37° C. After pelleting cultures, the clear supernatants were passed through 0.22 mm filters. Samples were analyzed for SCFAs using a BioRad HPLC organic acid system with AMINEX 87H, 300x7mm column with 5 mM sulfuric acid eluant at 0.6 ml/min, 65° C, with refractive index detector. Uninoculated media were used as blank and media background was subtracted except for glucose peak. DP: 2,2'-dipyridyl added at 50 μM. Values are means (mM) ± SE of the mean.

### Supplemental File S7

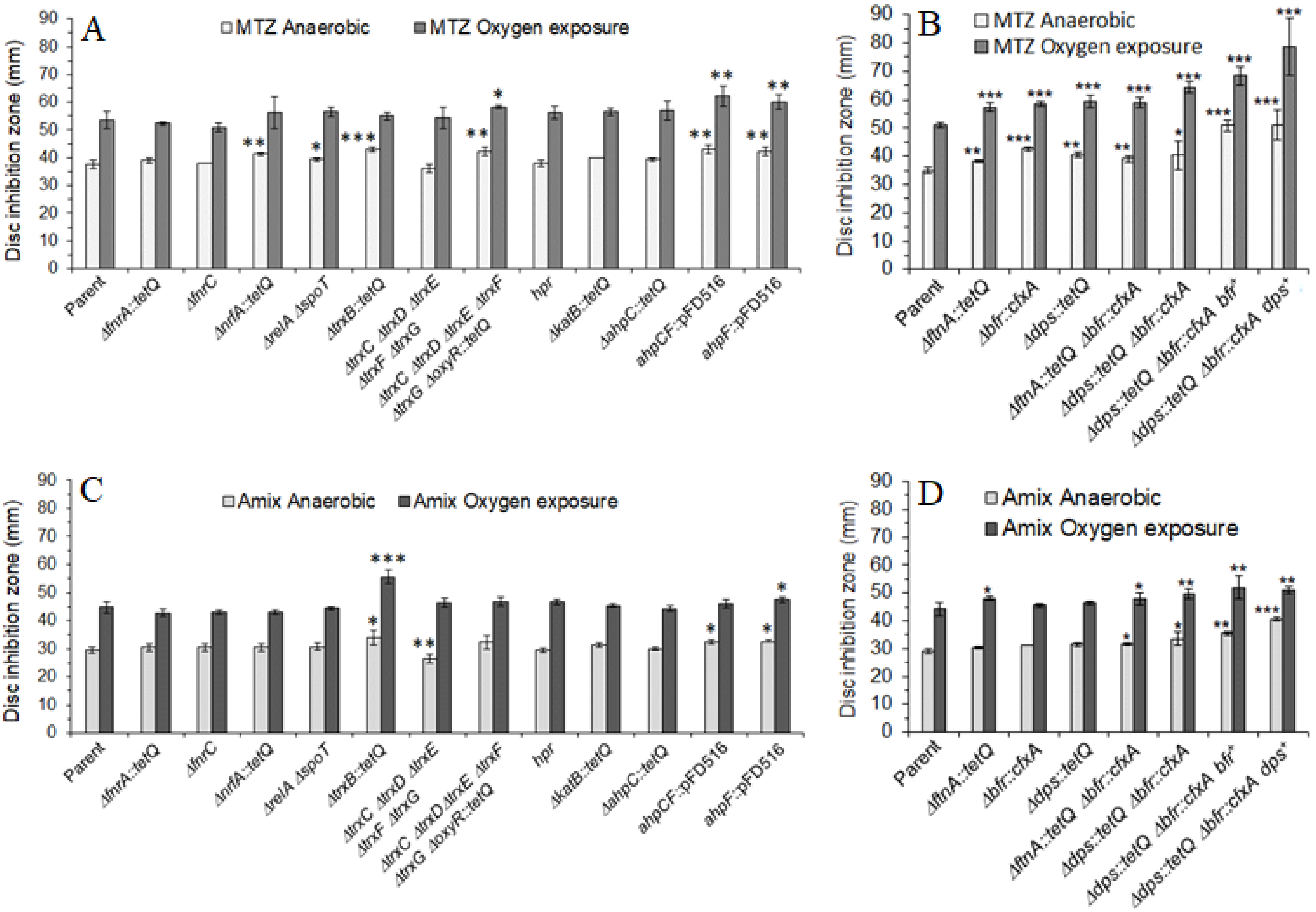

**Supplemental file S7**. Disc diffusion assay sensitivity for metronidazole (MTZ), panels A and B, and for Amixicile (Amix), panels C and D. Parent (*B. fragilis* 638R) and its isogenic mutant strains are depicted in each panel, respectively. Panels A and C) show mutant strains in the oxidative and redox stress responses, stringent response (*relA/spoT*), nitrite reductase (*nrfA*), and anaerobic regulators (*fnrA* and *fnrC*). Panels B and D) show mutants in the iron-storage family proteins *ftnA*, *bfr*, and *dps*, and genetic complemented strains, respectively. Strains designations are shown in each panel. Each bar represents the average zone inhibition (mm) of at least three independent biological replicates. Vertical error bars denote standard deviation of the means from two independent experiments in triplicate. The significance of the *P* value was calculated between parent strain control and each mutant strain following an unpaired *t* test (parametric and two-tailed) two groups is shown above the horizontal bars. *p<0.05; **p<0.01; ***p<0.001.

### Supplemental File S8.

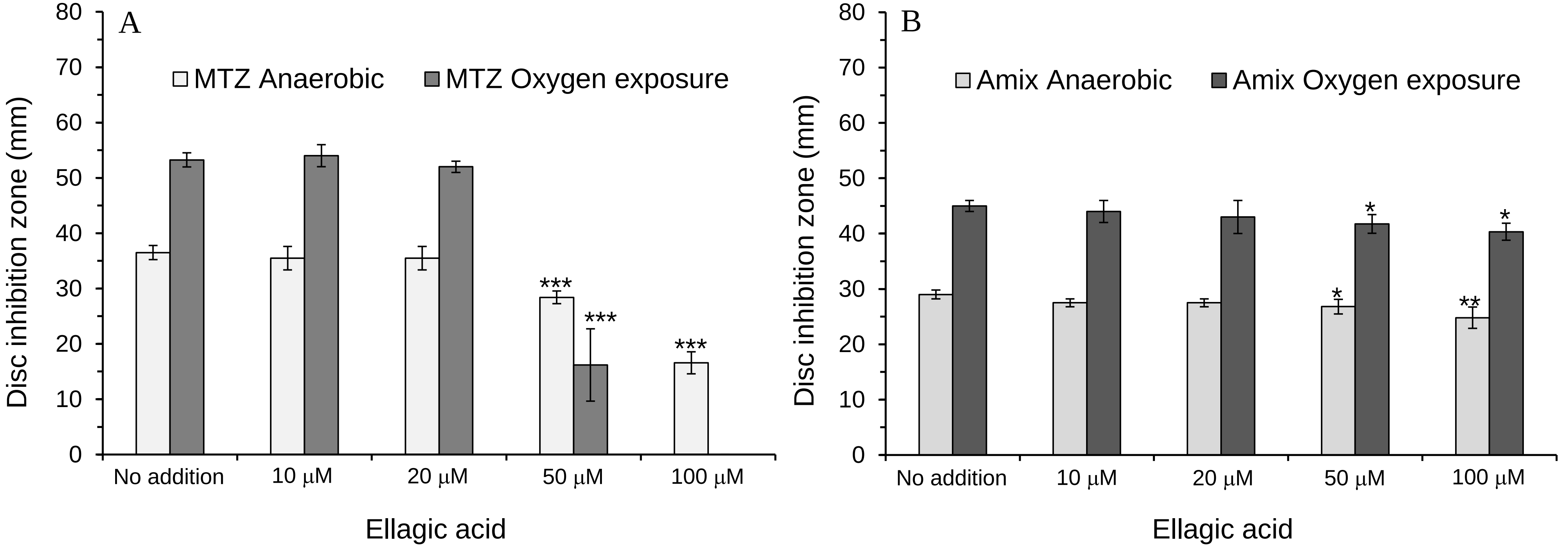

**Supplemental file S8**. Disc diffusion assay sensitivity of *B. fragilis* 638R to A) metronidazole (MTZ) or to B) Amixicile (Amix) in the presence of Ellagic acid. Ellagic acid was added to the BHI media at concentration described in each panel. Each bar represents the average zone inhibition (mm) of at least three independent biological replicates. Vertical error bars denote standard deviation of the means. The significance of the *P* value was calculated between no addition control and each experimental group following an unpaired *t* test (parametric and two-tailed) two groups is shown above the horizontal bars. *$p<0.05$; **$p<0.01$; ***$p<0.001$.

### Supplemental File S9

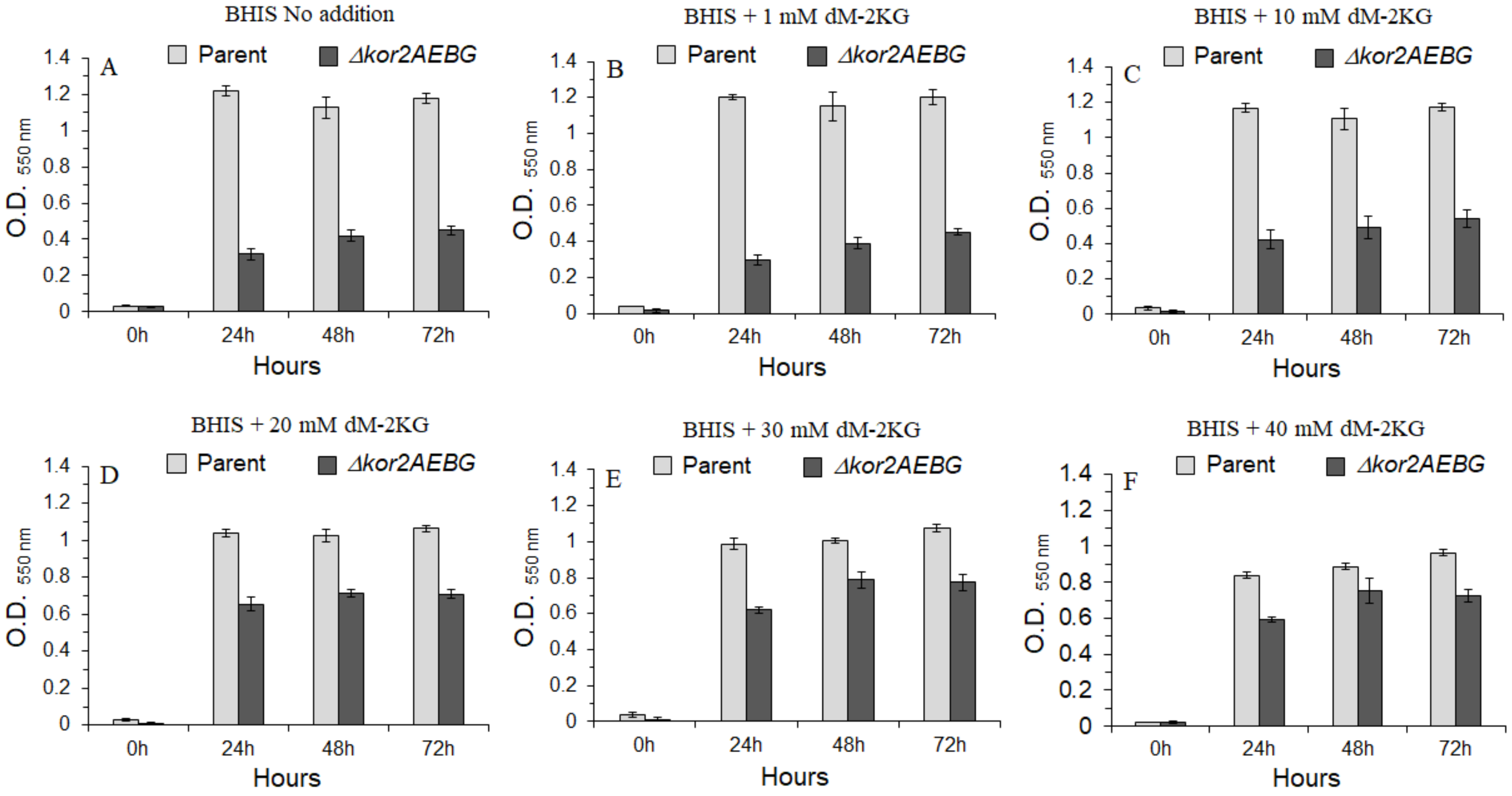

**Supplemental file S9.** Growth of *B. fragilis* 638R (parent) strain and the isogenic *Δkor2AEBG* mutant strain in supplemented brain-heart infusion (BHIS) media. Dimethyl-2-ketoglutarate (dM-2KG) was added as indicated in the panels. Strain designations are depicted for each panel. Media preparations and compositions are described in Materials and Methods section. Each bar represents the average of at least two independent experiments in triplicate. Vertical error bars denote standard deviation of the means.

### Supplemental File S10

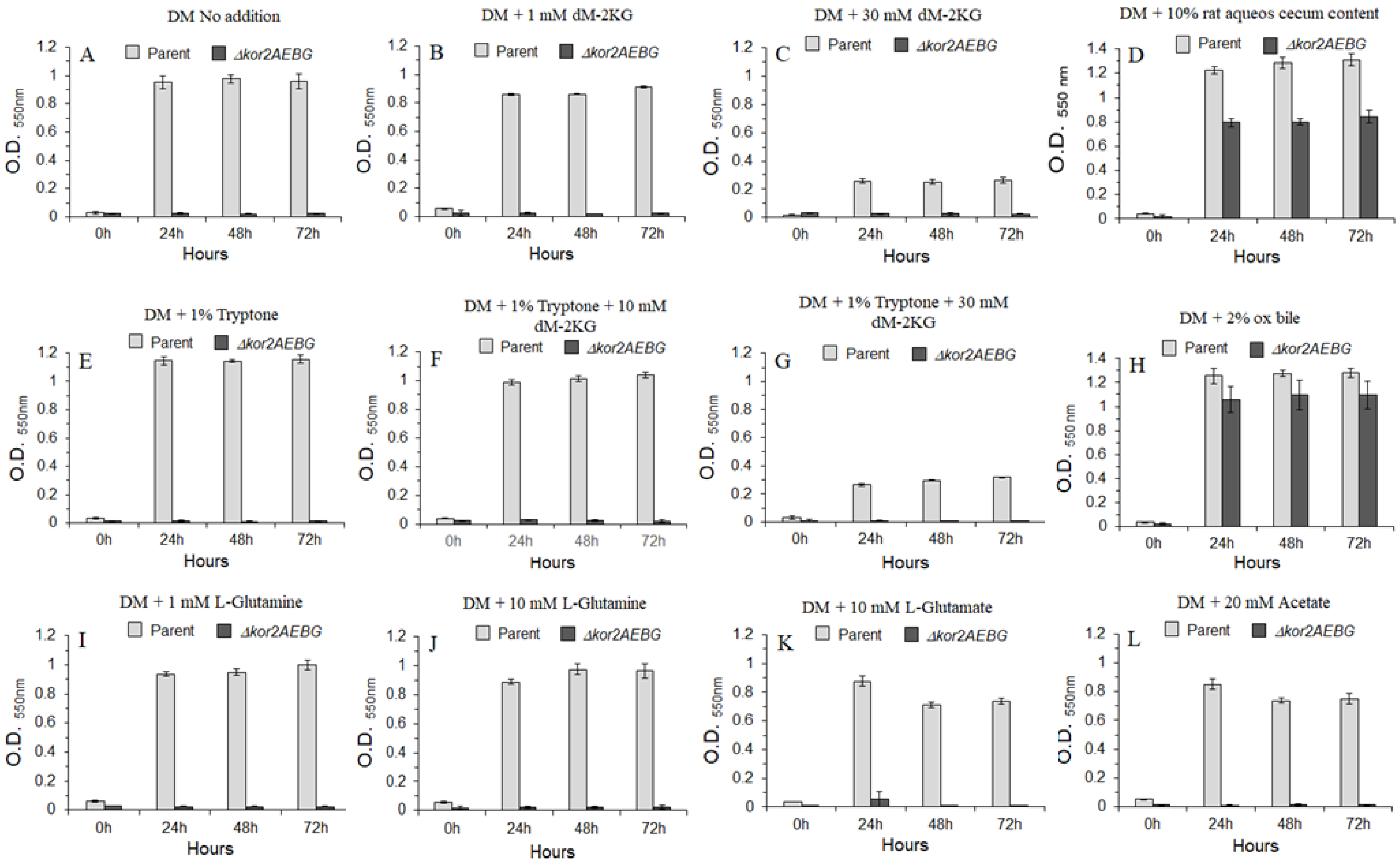

**Supplemental file S10.** Growth of *B. fragilis* 638R (parent) strain and the isogenic *Δkor2AEBG* mutant strain in chemically defined media with glucose (DM). DM was supplemented with 1% Tryptone, Dimethyl-2-ketoglutarate (dM-2KG), L-glutamine, L-glutamate, acetate, rat cell-free sterile aqueous cecum content, or ox bile as indicated in each panel. Strain designations are depicted for each panel. Media preparations and compositions are described in Materials and Methods section. Each bar represents the average of at least two independent experiments in triplicate. Vertical error bars denote standard deviation of the means.

#### Supplemental File S11

**Supplemental file S11**. Metronidazole minimal Inhibitory concentration (MIC µg/ml) for *B. fragilis* 638R in BHIS media supplemented with 2-hydroxy-1,4-naphthoquinone

| Media supplement/ MTZ MIC µg/ml | Anaerobic | $O_2$ exposure |
|---|---|---|
| Control (No addition) | 0.5 | 0.5 |
| 2-hydroxy-1,4-Naphthoquinone 10 µM | 8 | 4 |
| 2-hydroxy-1,4-Naphthoquinone 50 µM | 16 | 8 |
| 2-hydroxy-1,4-Naphthoquinone 100 µM | 32 | 32 |
| 1,4-Naphthoquinone 10 µM | 2 | 2 |
| 1,4-Naphthoquinone 50 µM | 16 | 16 |
| 1,4-Naphthoquinone 100 µM | 32 | 32 |

### Supplemental File S12

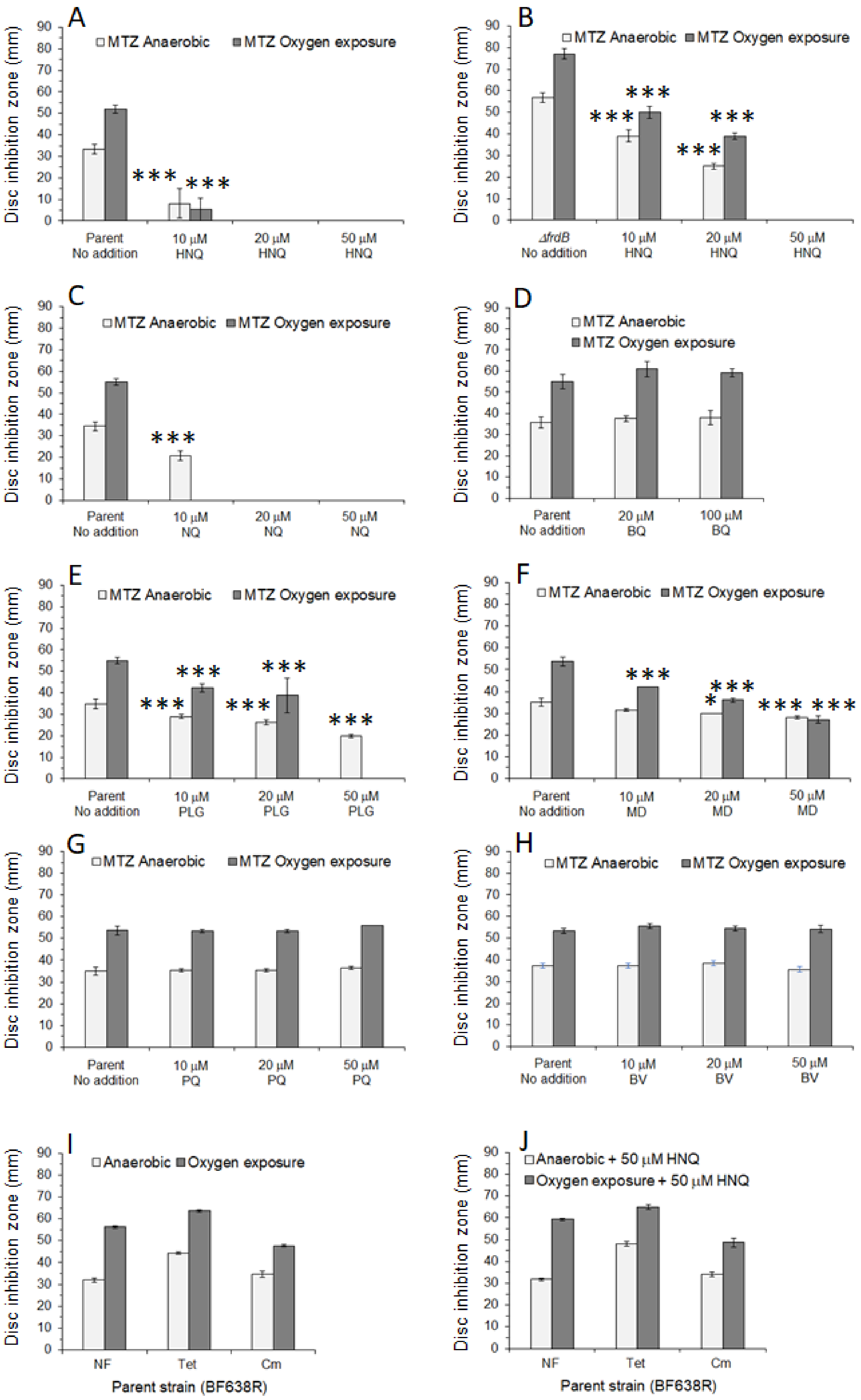

**Supplemental file S12**. Disc diffusion assays. Susceptibility of *B. fragilis* 638R strain (Parent) and its isogenic *ΔfrdB* deletion mutant to metronidazole (MTZ). Panels A and B: in the presence of 2-hydroxy-1,4-napththoquinone (HNQ). Panel C: in the presence of 1,-4-naphthonquinone (NQ). Panel D: in the presence of 1,4-benzoquinone (BQ). Panel E: in the presence of plumbagin (PLG). Panel F: in the presence of menadione (MD). Panel G: in the presence of paraquat (PQ). Panel H: in the presence of benzyl-viologen (BV). . Panels I and J: Susceptibility of *B. fragilis 638R* strain to nitrofurantoin (NT), tetracycline (Tet), or chloramphenicol (Cm) exposed to oxygen with no addition (I) or addition of 50 μM HNQ (J). Each bar represents the average zone inhibition (mm) of at least three independent biological replicates. Vertical error bars denote standard deviation of the means from two independent experiments in triplicate. The significance of the *P* value was calculated between no addition control and each experimental group following an unpaired *t* test (parametric and two-tailed) two groups is shown above the horizontal bars. *p<0.05; **p<0.01; ***p<0.001.

### Supplemental File S13

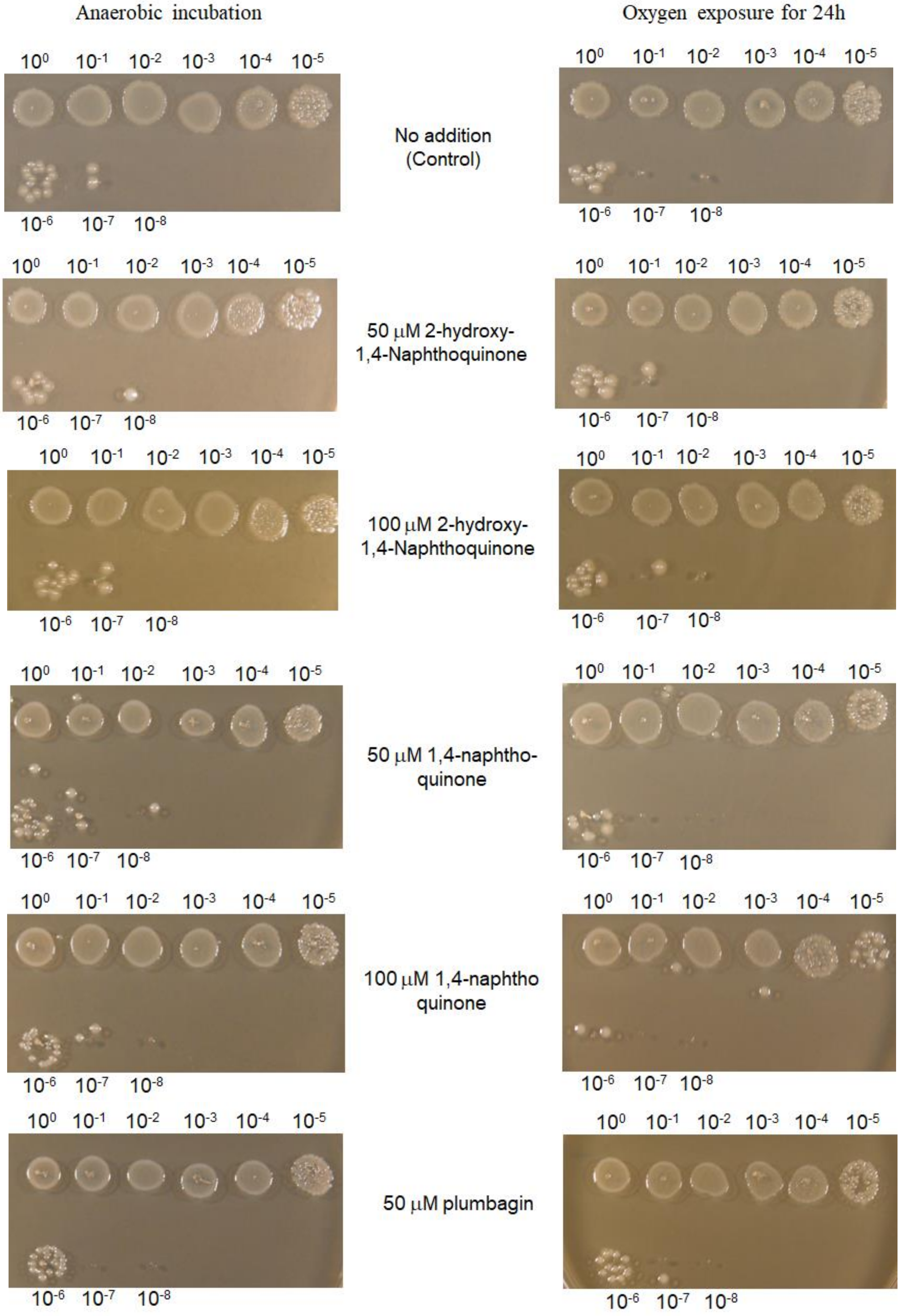

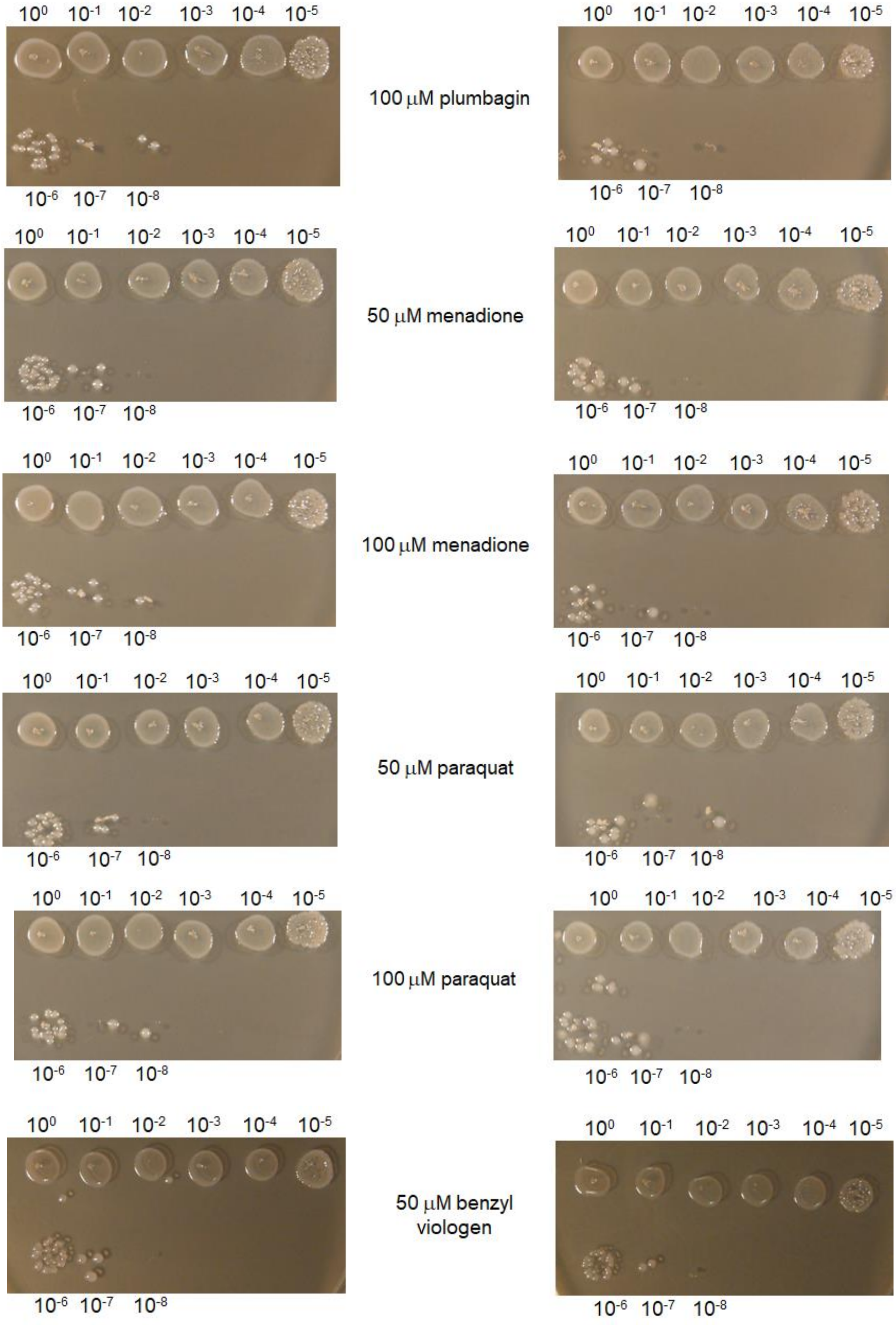

100 μM plumbagin
50 μM menadione
100 μM menadione
50 μM paraquat
100 μM paraquat
50 μM benzyl viologen

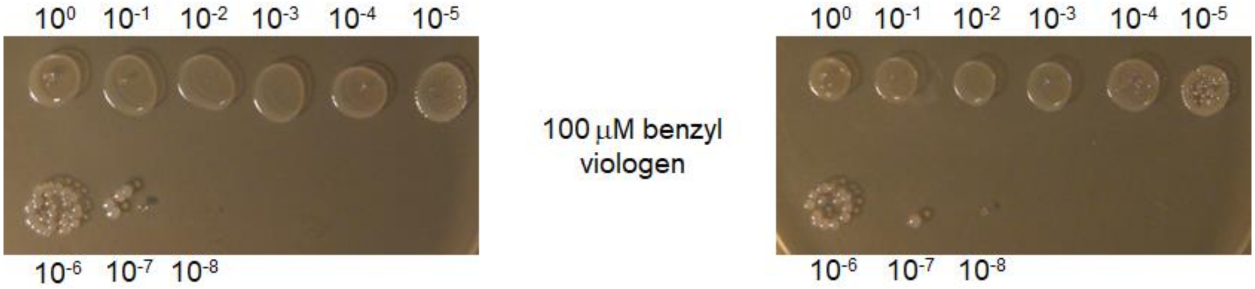

**Supplemental file S13**. Survival of *B. fragilis* 638R on BHIS supplemented with redox cycling agents as shown in each panel. Bacteria were grown in BHIS broth overnight and serially diluted in PBS, pH 7.4. Five μl aliquot from each dilution was place on top of the plate as indicated, respectively. Plates were inoculated in duplicated, and one set was immediately incubated in an anaerobic chamber incubator at 37° C. The second set was placed in an aerobic incubator at 37° C for 20-24 h prior to incubation in an anaerobic chamber incubator. Dilution factor is indicated above and below the panels.

### Supplemental File S14

**Supplemental file S14.** Closantel minimal inhibitory concentration (MIC) for *Bacteroides* species in BHIS media.

| Strains/Closantel MIC μg/ml (μM) | Anaerobic | $O_2$ exposure |
|---|---|---|
| *B. fragilis* 638R | 8.6 (12.5) | 8.6 (12.5) |
| *B. fragilis* BF8 | 4.3 (6.25) | 4.3 (6.25) |
| *B. fragilis* ATCC 25285 | 4.3 (6.25) | 4.3 (6.25) |
| *B. thetaiotaomicron* VPI 5482 | 274.4 (400) | 34.3 (50) |
| *B. vulgatus* ATCC 8482 | 2.15 (3.12) | <0.01325 (<0.1953) |

### Supplemental File S15

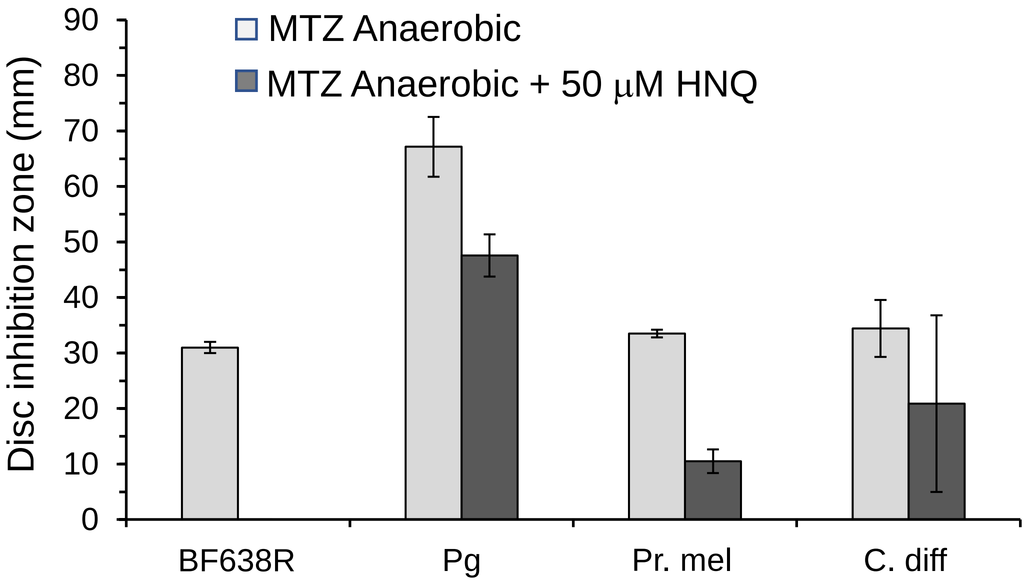

**Supplemental file S15**. Disc diffusion assays for MTZ susceptibility in the presence of 50 μM 2-hydroxy-1,4-naphthoquinone (HNQ). BF638R: *B. fragilis* 638R. Pg: *Porphyromonas gingivalis* ATCC 33277. Pr. mel: *Prevotella melaninogenica* ATCC 25845. *Clostridioides difficile* ATCC 43255. For this experiment BHI was supplemented with 5% defibrinated laked sheep blood. Each bar represents the average zone inhibition (mm) of at least three independent biological replicates. Vertical error bars denote standard deviation of the means from two independent experiments in triplicate.

**Supplemental Table S1**. List of oligonucleotide primers used in this study.

| Primer name | Nucleotide Sequence |
|---|---|
| PIR1-pBT (EcoRI) | GAAAGGAATTCTATGAAAACAG |
| PIR1-pBT (BamHI) | GAATACGGATCCTTATCGCATTGG |
| PFOR-pTRG-FOR | GACTAAACAGAAGGGATCCATC |
| PFOR-pTRG-REV | GTTATTATCGGATCCAAGAT |
| PIR1-FOR-RTPCR | CAATCATGGTTGGCTGAAAA |
| PIR1-REV-RTPCR | GAATCCCATTTCAGGGTCAA |
| PIR1mut-BglII-FOR | GGACGCTTCAAGATCTCATTCC |
| PIR1mut-BamHI-REV | CCGACGGATCCAACTGTTACC |
| PIR1mut-SstI-FOR | GATACAAAGAGCTCTAAAGTGG |
| PIR1mut-SstI-REV | GTAGAGAAGAGCTCAGGTGCCG |
| PIR1-pFD-BamHI-FOR | CGTTTGGGATCCAGATGTTC |
| PIR1-pFD-SstI-REV | AAGGGCTGAGCTCGCTATTGAAGG |
| PIR2-pBT (EcoRI) | GAAAGGAATTCCATCATGAAA |
| PIR2-pBT (BamHI) | GCGGATCCTTTAAATTTACATCGG |
| ADH-pTRG-FOR | ATTAGGAGGATCCATAATGC |
| ADH-pTRG-REV | GTACTGAGGATCCCTACTCCGGG |
| PIR2-FOR-RTPCR | GGGTTCATTTCGGTGCTCT |
| PIR2-REV-RTPCR | CATGTGCCAGAAGACCCTTT |
| PIR2mut-BglII-FOR | CCCACCTGAAGATCTGTCGGAG |
| PIR2mut-BamHI-REV | CATATACGGATCCTCCGCGGG |
| PIR2mut-EcoRI-FOR | GGAGTCTACGAATTCCTGATCG |
| PIR2mut-EcoRI-REV | GGGATACTGAATTCGCGGCTG |
| PIR2-pFD-BamHI-FOR | GGTTTATAGGATCCATTTATAAAAG |
| PIR2-pFD-SstI-REV | CACTCCCGAGCTCGTTTCCGTG |
| pTRG-Forward | TGGCTGAACAACTGGAAGCT |
| pTRG-Reverse | ATTCGTCGCCCGCCATAA |
| pBT-Forward | TCCGTTGTGGGGAAAGTTATC |
| pBT-Reverse | GGGTAGCCAGCAGCATCC |
| Pyc-InsF | CGATGAATTCCGCAGGCAACTTTGTGCA |
| Pyc-InsR | CGATGGATCCAACGGACCGGGTAAAC |
| FNR_1017-NT-PstI | GACCTGCAGGCCATAAATGCTGCCATGGCC |
| FNR_1017-NT-SOE | CCACATCTCTGCAATACGTTCGGG |
| FNR_1017-CT-SOE | CCCGAACGTATTGCAGAGATGTGGCGACTGATCACCATCGACGGACG |
| FNR_1017-CT-XbaI | GCATCTAGAGAAGCAATGGCTGCTATTGC |
| FNR_1017-check-F | CTGTTATTCACGAAGTAAGCC |

| | |
|---|---|
| FNR_1017-check-R | GCAATGATTTGTACGTCCGG |
| Kor1-BamHI-FOR3 | GAAAGT<u>GGATCC</u>AAACAAAAAACGC |
| Kor1-REV-SEW2 | CTGCTCGGCGACTGCATCCGGAATCTGTGATGACCAGCC |
| Kor1-FOR-SEW | GGCTGGTCATCACAGATTCCGGATGCAGTCGCCGAGCAG |
| Kor1-SalI-REV | GATCTGTTT<u>GTCGAC</u>AGTATGTGGTGC |
| Kor1-mutcheck-FOR | CACTAACACCGGTTCATGTTAATCC |
| Kor1-mutcheck-REV | GACTATGCGGTTACGACAGATTCG |
| Kor2-PstI-FOR | GATCGGGTTCA<u>CTGCAG</u>ATTAGCG |
| Kor2-REV-SEW | ACCGGGACTTGATGATGAGG |
| Kor2-FOR-SEW | CCTCATCATCAAGTCCCGGTCCGAACGCCACCATCACC |
| Kor2-BamHI-REV | GACTGAGTGTTACTGAT<u>GGATCC</u>GCG |
| Kor2-mutcheck-FOR | GTAATATCCCTCGCCAAAGAGG |
| Kor2-mutcheck-REV | GGCTTTCGCTGACGTGCC |
| Kor2comp-BamHI-FOR | GATC<u>GGATCC</u>GCAACCGTATGTCCGGACGG |
| Kor2comp-SacI-REV | GATCGAGCTCCCGCTATCTCTGCCTGTGGC |
| PoxB-BamHI-FOR | GATC<u>GGATCC</u>GACGGATCTGCTCCATCATCCG |
| PoxB-REV-SEW | GGACCTTCCAGCTCGAAAGCGCCGGTCAGTTGCGCTTCTGC |
| PoxB-FOR-SEW | GCAGAAGCGCAACTGACCGGCGCTTTCGAGCTGGAAGGTCC |
| PoxB-PstI-REV | GATCCTGCAGGGAATCGGATCGACAGACCG |
| PoxB-mutcheck-FOR | TTTCCGGCATTCAAGCCG |
| PoxB-mutcheck-REV | CGCGAATGAAAGGTTTGAGG |
| Acn-BamHI-FOR | GATC<u>GGATCC</u>ACAATTCTGTTTCCGGGTTGG |
| Acn-SEW-REV | GCCAAGTCACGTGCGAGTTCCCTTCCATCTTCCGGGGATAACC |
| CitS-SEW-FOR | GGTTATCCCCGGAAGATGGAAGGGAACTCGCACGTGACTTGGC |
| CitS-SalI-REV | GATC<u>GTCGAC</u>CCCAAACTCTTCCCGATAGCG |
| Acn-mutcheck-FOR | GGTCCGTTTACTCCGGACGC |
| CitS-mutcheck-REV | TCTGTGCAATCTGTAAATCTGCGG |
| AcnBglIIcompl-FOR | GATCAGATCTAATAAATAGAAGAACAGATAAAACCGAAA |
| CitSacIcompl-REV | GATCGAGCTCCCACCACAGATTGGCAGACGG |
| FrdC-BamHI-FOR | GATC<u>GGATCC</u>GCACAATGAGAAAGAAGGCTCG |
| FrdC-BamHI-REV | GATC<u>GGATCC</u>GGCAAGTCCGGTACCTACAACG |
| FrdB-BamHI-FOR | GATC<u>GGATCC</u>CCGGTGAAAATTCAGAACCGG |
| FrdB-SacI-REV | GATC<u>GAGCTC</u>GTTCCTTGCGAGCCTGACGG |

---

The underlined nucleotide sequences denote restriction sites used for cloning into appropriate plasmid vector restriction sites respectively.
